# Observations on Paleospecies Determination, With Additional Data on *Tyrannosaurus* Including Its Highly Divergent Species Specific Supraorbital Display Ornaments That Give *T. rex* a New and Unique Life Appearance

**DOI:** 10.1101/2022.08.02.502517

**Authors:** Gregory S. Paul

## Abstract

Intrageneric dinosaur species have been being named for decades without either significant examination of the methods and standards used to do so, or widely publicized controversy over the results. The long standing assumption that all large known specimens of the iconic North American *Tyrannosaurus* consisted of just the one popular species *T. rex* was recently challenged with the first comprehensive test of the question. The result was the diagnosing and naming of two additional taxa, *T. imperator* and *T. regina,* based on a number of species levels characters regarding robustness and tooth proportions in the context of their stratigraphic distribution. In association a rare in-depth look was taken at the current state of naming vertebrate paleospecies, which it turns out are not highly rigorous because of inherent problems with the species concept and other matters. The results of the paper were severely criticized in in a manner never seen before for new dinosaur species even when based on less evidence. This study takes another look as the determination of paleospecies, and shows that many of the claims made in the criticisms regarding the *Tyrannosaurus* species work were inaccurate. New data on the proportions of strength bars in *Tyrannosaurus* skulls reinforces the basing of the three species in part on robustness factors, and allows all but one skull to be assigned to one of the species. These results allow the first detailed systematic examination of the supraorbital display bosses of the genus. They sort out as visually distinctive species specific ornaments based on both stratigraphic and taxonomic factors, strongly affirm that *Tyrannosaurus* was multispecific, and the species probably dimorphic. New skulls of *T. rex* show that the species sported – males probably -- striking display bosses not yet observed in other tyrannosaurids.

## INTRODUCTION

From around the turn of the century until now a number of academic studies have examined and sometimes named new sibling paleospecies within dinosaur genera, in all but one case with little immediate post publication criticism in the news media (Barrett et al. 2005; Evans and Reisz 2007; Sereno 2010; Scannella et al. 2014; Tschopp et al.; Campbell et al. 2016; MacDonald & Currie 2018; Fowler and Freedman 2020). Most of these and other papers on dinosaur species have not explicitly addressed at length the methods used within to determine fossil species, Paul (2006) and Carpenter (2010) being rare exceptions. It appears that most of those investigating issues concerning paleospecies have done so in an ad-hoc manner, which is the general situation regarding extinct tetrapods. While assessing the species contained within *Tyrannosaurus,* the long term study by Paul et al. (2022) included (mainly in the supplement) an extended look at the state of paleospecies work on dinosaurs and other groups to better determine whether or not the study met the standards on hand. The analysis concluded that Tyrannosaurus included three species, *T. rex*, *T. regina*, and *T. imperator*.

In direct response to Paul et al. (2022), Carr et al. (2022) produced a technical reply that concluded that only *T. rex* is valid. Carr et al. (2022) was unusual in the quickness of the submission after Paul et al. (2022). Also atypical is that a major portion of Carr et al. (2022) repeats arguments made in an itself uncommon wave of criticisms of Paul et al. (2022) by a number of paleozoologists in the news and social media, such having not been not seen before regarding other dinosaur taxa conducted under broadly similar circumstances (Ashworth 2022; Barras 2022; Carr 2022; Davis 2022; Dunham 2022; Elbein 2022; Greshko 2020; Hernandez 2022; Hunt, 2022; Kim 2022; Kruger and Ricci 2022; Lawrence 2022; Witton 2022). Many of the comments directly claimed or implied that certain and strict methods must be used to determine paleospecies. That was done despite many of the claimed minimal criteria being contrary to those documented as being commonly used in the previous paleozoological literature in Paul et al. (2022).

This analysis in part a response to Carr et al. (2022). Building on Paul et al. (2022), the below investigation also takes the opportunity to further discuss what needs to be done, as well what is not necessary, when testing and determining the number of closely related sibling species proposed to be present within paleogenera. After those standards are detailed new data that further tests the status of species within *Tyrannosaurus* is offered, including the first demonstration that the three species each sported visually distinctive species style cranial display decorations of the type expected between and used to diagnose species, and the diagnoses of the three named species are revised and expanded. A follow up to Paul et al. (2022), the great majority of this science research paper was produced prior to Carr et al. (2022), in parallel, Persons and Van Raalte are preparing a more statistical oriented analysis of the cumulative data for *Tyrannosaurus* that will be presented when finalized.

## SYSTEMATICS

### General Systematic Diagnostic Procedures

The below are differential, contrasting minimal character diagnoses. Diagnostic characters are sometimes overlapping nonbimodal between taxa at a given level, and not entirely consistent within taxon as per Maisch (2008), Maxwell (2012), Scannella et al. (2014), MacDonald & Currie (20180, Harvati & Ackermann (2022), Paul et al. (2022), Carr et al. (2022), for more information see further discussions below.

### Derived Tyrannosaurines

Revised diagnoses for Asian and North American derived tyrannosaurine genera and species. Characters are from large specimens. Those for North American *Tyrannosaurus* are based primarily on TT-zone specimens, with some specimens from geographically lateral latest Maastrichtian formations, further detailed and expanded from Paul et al. (2022). MGE, most gracile example/s in the genus; MRE, most robust example/s in the genus, autapomorphies relative to other tyrannosaurid species indicated by asterisks. Including all specimens available for this analysis, it is expected that the diagnoses will be further modified with the inclusion of new specimens and any revisions in the data base. A few specimens have been reassigned or their status modified.

### Incorporating Three Species of TT-zone *Tyrannosaurus*

This first set of diagnoses for all pertinent taxa is for three species of TT-zone

*Tyrannosaurus* Tyrannosaurinae Osborn, 1906

*Tarbosaurus* Maleev, 1955

*Diagnosis*: Temporal region breadth well under twice that of rostrum and less than half length of skull, orbits and lateral face of jugal do not face substantially anteriorly, nasal is well over half length of skull and strongly domed, anterodorsal process on anterior ramus of lacrimal absent, lacrimals do not nearly meet at midline and lateral swelling on supraorbital process of lacrimal is present, vomer has extensive contact with premaxilla and a deep ventral flange, 12-13 maxillary and 14-15 dentary teeth, large teeth not as robust, usually or always two slender functional anterior incisiform dentary teeth; pubic boot moderate in size, lower hindlimb elements longer relative to femur.

*Tarbosaurus bataar* Maleev, 1955

*Holotype*: PIN 551-1.

*Referred specimens*: As listed in Hurum and Sabath (2003).

*Diagnosis*: Gigantic at 4-5 tonnes; interfenestral pillar broad, postorbital bosses are fairly subtle subcircular, knob-like discs limited to the frontal process that do not project well above the dorsal rim of the skull; femurs are robust relative to general tyrannosaurid curve.

#### Tyrannosaurus Osborn, 1905

*Diagnosis*: Temporal region about twice as broad as rostrum and over half length of skull, orbits and lateral face of jugal face substantially anteriorly, nasal a little over half length of skull and not as strongly domed, presence of anterodorsal process on anterior ramus of lacrimal projects into nasal, lacrimals nearly meet at midline, sublunate in dorsal shape partly because lateral swelling on supraorbital process is absent, vomer sometimes has anterior spear point, contact with premaxilla not as extensive and deep ventral flange is absent, 11-12 maxillary and 12-14 dentary teeth, large teeth robust; pubic boot massive, lower hindlimb elements not as long relative to femur.

#### Tyrannosaurus imperator Paul et al., 2022

*Holotype*: FMNH PR2081.

*Referred specimens*: BHI 4182, 6248/?AMNH 3892, <u>HMN MB.R.91216,</u> MOR 1125, 1128, RGM 792.000, SDSM 12047, TCM 2001.90.1, TMT v2222, MOR 008?, NMNNH P-3698?.

*Age and Stratigraphy*: Late Maastrichtian, lower, lower middle and possibly middle Hell Creek and Lance, Laramie, Arapahoe, McRae?, North Horn?, Javelina?.

*Geographic distribution*: Montana, Dakotas, Wyoming, New Mexico?, Texas?, Utah?.

*Diagnosis*: Gigantic at 6-8 tonnes; generally robust, always or usually so regarding maxilla all being more robust than *T. regina* but not all *T. rex*, interfenestral pillar (MRE) all being more robust than for *T. rex* and especially *T. regina*, lacrimal (MRE), postorbital process of jugal (MRE), quadratojugal (MRE), dentary, humerus, ilium (MRE) all being more robust than for *T. rex* and *T. regina*, metatarsals 2 and 4, with length/circumference ratios of 2.4 or less for the femur (MRE); usually two slender functional anterior incisiform dentary teeth; very rugose nasals sometimes present, postorbital bosses are sometimes large, prominent horizontally extended *spindles that extend posteriorly to close to or on the anterior squamosal process, do not project much above dorsal rim of the skull presumably among males, antero-medial processes can be fairly well developed.

#### Tyrannosaurus rex Osborn, 1905

*Holotype*: CM 9380.

*Referred specimens*: BHI 6230, 6233, 6435, 6436, UWBM 99000, RSM 2523.8, BHI 4100?, NHMUK R7994?, UCMP 118742?.

*<u>Age and stratigraphy</u>*: Latest Maastrichtian, upper and possibly middle Hell Creek and Lance, Ferris, Denver, Frenchman, Willow Creek, Scollard.

*<u>Geographic distribution</u>*: Montana, <u>Colorado,</u> Dakotas, Wyoming, Alberta, Saskatchewan.

*Diagnosis*: Gigantic at 6-8 tonnes; generally robust, but overall less so than *T. imperator*, always or usually so regarding maxilla (MRE) all being more robust than for *T. regina* but not for all *T. imperator*, interfenestral pillar all being more robust than for *T. regina* while being more gracile than for *T. imperator,* postorbital process of jugal all being more robust than for *T. regina* but not all *T. imperator*, quadratojugal, dentary (MRE), and in some cases metatarsals, with length/circumference ratios of 2.4 or less for the femur; usually one slender functional anterior incisiform dentary tooth and no examples with a truly small second dentary tooth yet observed; very rugose nasals sometimes present, postorbital bosses are sometimes prominent subcircular, knob-like discs limited to the frontal process that *project well above the dorsal rim of the skull presumably among males, antero-medial processes weakly developed.

#### Tyrannosaurus regina Paul et al., 2022

*Holotype*: USNM 555000.

*Referred specimens*: NHMAD “S”, MOR 980, LACM 23485, LL 12823, TMP 81.6.1,

LACM 23844?, 23845?.

*<u>Age and stratigraphy</u>*: Latest Maastrichtian, upper and possibly <u>middle</u> Hell Creek and Lance, Ferris, Denver, Frenchman, Willow Creek, Scollard.

*Geographic distribution*: Montana, Colorado, Dakotas, Wyoming, Alberta, Saskatchewan.

*Diagnosis*: Gigantic at 6-8 tonnes; generally gracile, more so than *T. rex* and much more than *T. imperator*, always or usually so regarding maxilla (MGE) all more gracile than *T. rex* or *T. imperator*, interfenestral pillar (MGE) all more gracile than *T. rex* or especially *T. imperator*, lacrimal (MGE), postorbital process of jugal (MGE) all more gracile than *T. rex* but not all *T. imperator*, quadratojugal (MGE), dentary (MGE), humerus (MGE), ilium all more gracile than *T. imperator*, *femur gracile relative to scaling norm for tyrannosaurids with length/circumference ratios of 2.4 or higher; one slender functional anterior incisiform dentary tooth and no examples with a truly small second dentary tooth yet observed; very rugose nasals not yet observed, postorbital bosses are neither spindles nor knobs, are not as posteriorly limited as in *T. rex*. do not project much above dorsal rim of the skull, are sometimes *hat shaped presumably among mature males.

#### *Tyrannosaurus* incertae sedis

Robusts of uncertain or middle TT-zone stratigraphic position that are probably *T. Imperator* or *T. rex*-- BHI 6231, 6232, 6242, USNM 6183; of uncertain proportions and high stratigraphic placement that are probably *T. rex* or *T. regina*-- BHI 6249<u>, DMNS</u> <u>2827,</u> MOR 009<u>, TMP 81.12.1</u>; insufficient proportional and/or stratigraphic information for a species assignment -- AMNH 5027, 30564, CM 1400, RMDRC 2002.MT-001.

#### Tyrannosauridae incertae sedis

BMRP 2002.4.1, 2006.4.4, CMNH 7541, DDM 344.1, LACM 28741, RSM 2990.1, RSM 2347.1, TMM 41436-1, 46028-1, UMNH 11000, NCMNS “BM”, “Jodi”.

### Incorporating Only Two Species of TT-zone *Tyrannosaurus*

These TT-zone *Tyrannosaurus* diagnoses are for just two species arbitrarily assuming that *T. regina* is a junior synonym of *T. rex*, see further discussion below.

#### Tyrannosaurus imperator Paul et al., 2022

*Holotype*: FMNH PR2081.

*Referred specimens*: BHI 4182, 6248/?AMNH 3892, <u>HMN MB.R.91216,</u> MOR 1125, 1128, RGM 792.000, SDSM 12047, TCM 2001.90.1, TMT v2222, MOR 008?, NMNNH P-3698?.

*Age and Stratigraphy*: Late Maastrichtian, lower, lower middle and possibly middle Hell Creek and Lance, Laramie, Arapahoe, McRae?, North Horn?, Javelina?.

*<u>Geographic distribution</u>*: Montana, Dakotas, Wyoming, New Mexico?<u>, Texas?</u>, Utah?.

*Diagnosis*: Gigantic at 6-8 tonnes; usually two slender functional anterior incisiform dentary teeth; postorbital bosses are sometimes large, prominent horizontally extended

*spindles that extend posteriorly to close to or on the anterior squamosal process, do not project much above dorsal rim of the skull presumably among males.

#### Tyrannosaurus rex Osborn, 1905

*Holotype*: CM 9380.

*Referred specimens*: BHI 4100, 6230, 6233, 6249, 6435, 6436, NHMAD “S”, LACM 23484, 23485, DMNS 2827, LL 12823, MOR 009, 980, NHMUK R7994, RSM 2523.8, TMP 81.6.1, 81.12.1, UCMP 118742. USNM 555000, UWBM 99000.

*<u>Age and stratigraphy</u>*: Latest Maastrichtian, upper and possibly middle Hell Creek and Lance, Ferris, Denver, Frenchman, Willow Creek, Scollard.

*<u>Geographic distribution</u>*: Montana, <u>Colorado,</u> Dakotas, Wyoming, Alberta, Saskatchewan.

*Diagnosis*: Gigantic at 6-8 tonnes; usually one slender functional anterior incisiform dentary tooth and no examples with a truly small second dentary tooth yet observed; postorbital bosses are sometimes prominent subcircular, disc-like knobs limited to the frontal process that *project well above the dorsal rim of the skull presumably among males.

#### *Tyrannosaurus* incertae sedis

Robusts of uncertain stratigraphic position that are probably *T. Imperator* or *T. rex*-- BHI 6231, 6232, 6242, USNM 6183; insufficient proportional and/or stratigraphic information for a species assignment -- AMNH 5027, 30564, CM 1400, RMDRC 2002.MT-001.

### Triceratops

Informal draft species diagnoses for large specimens of TT-zone *Triceratops* for evaluation of data quality and consistency within and between species within this genus compared to above results for *Tyrannosaurus*, see further discussion below. Based on characters utilized in Scannella et al., (2014), sample expanded to include some of the specimens outside the limited geographic area of the Hell Creek examined by Scannella et al. (2022) – listed referred specimens are those added in this study, for those referred to the species by Scannella et al. see that study. It is expected that the diagnoses will be further modified with the inclusion of additional specimens beyond those considered in Scannella et al. (2014) and herein as well as new finds, and revisions and expansions to the data base. The noncomprehensive referred specimen lists are only fossils not cited in Scannella et al (2014).

*T. horridus*

*Holotype: YPM 1820*

*Referred specimens*: AMNH 5116, MNHN 1912.20, SDSM 2760, TCM 2001.93.1,

USNM 1201, 2100, 2412, 4928.

*Diagnosi*s: Rostrum sometimes exceptionally elongated, length of nasal short to moderate, all shorter than *T*. sp. and *T. prorsus*, snout usually but not always shallow, angle between nasal process and narial strut of premaxilla usually shallow to sometimes acute, nasal process of premaxilla narrow, all narrower than *T. prorsus* but not *T*. sp., large anteromedial process on nasal; small dorsal boss on nasal, nasal horns very small to moderate size, all smaller than *T. prorsus,* postorbital horns short to very long; frontoparietal fontanelle constricted and closed in late subadults and young adults; variability within species high, and may in part represent dimorphism.

#### *T*. sp

*Diagnosi*s: Rostrum short. Length of nasal moderate, all longer than *T. horridus* but not all *T*. sp. Snout shallow or deep. Angle between nasal process and narial strut of premaxilla shallow to acute. Nasal process of premaxilla width moderate, always broader than *T. prorsus* but not all *T. horridus*. Large anteromedial process on nasal. Small dorsal boss on nasal. Nasal horns small to fairly large. Postorbital horn length moderate to very long. Frontoparietal fontanelle open in late subadults and young adults.

*T. prorsus*

*Holotype*: YPM 1822

*Referred specimens*: CM 1221, LACM 7207.

*Diagnosi*s: Rostrum length short to moderate. Length of nasal moderate to long, all longer than *T. horridus* but not all *T*. sp. Snout deep. Angle between nasal process and narial strut of premaxilla moderate to acute, all more acute than *T.* sp. but not all *T. horridus*. Nasal process of premaxilla broad, all broader than *T.* sp. and especially *T. horridus*. Small anteromedial process on nasal. No dorsal boss on nasal. Nasal horns fairly large to large, all larger than *T. horridus* but not all *T*. sp. Postorbital horn length moderate. Frontoparietal fontanelle constricted and closed in late subadults and young adults. Variability within species low, evidence for dimorphism lacking.

## METHODS AND MATERIALS

### For General Paleospecies Standards Assessment

Being an expansion on Paul et al. (2022, suppl.), this work’s discussion on determining and diagnosing vertebrate intrageneric species examines and cites a large variety of recent examples in the technical literature that have not been considered controversial in the theoretical methods utilized, there being few if any examples of such disputes even when the results are disputed. The soundness of the theoretical foundations behind work on paleospecies is examined and assessed from a practical perspective, and are used to test the validity of the assertions and conclusions regarding the subject in Carr et al. (2022, with the proceeding media comments made by the authors of that paper and others considered when necessary to more fully cover and test the issues under consideration), and for going forward in terms of new research. In order to better illustrate the skeletal similarity of some closely related dinosaur taxa their cranial display ornaments are removed.

The standards and practices assessment is used to ascertain the practical boundaries of what is and is not valid in defining sibling species in dinosaurs, in preparation for applying them to the species level taxonomy of *Tyrannosaurus.* The applicable results and methods are summarized at the end of the general paleospecies discussion.

### For Assessing *Tyrannosaurus* Species

The number of large *Tyrannosaurus* specimens examined for purposes of diagnosing the species by Paul et al. (2022) and herein is now 38, with 32 stratigraphically correlated. The stratigraphy was determined by methods used in P. Larson (2008), Carr (2020) and Paul et al. (1922). Juveniles are not scrutinized for direct diagnostic purposes, although they are assessed regarding their pertinence to the names of the new species.

The variations in anatomy and the stratigraphy techniques that were used in Paul et al. (2022) and within this investigation are directly compared to those regularly applied to other fossil vertebrates in the technical literature, dinosaurs especially.

The proportional data utilized in this study are in part those employed in Paul et al. (2022), which focused on the robustness versus gracility of skull and skeletal elements. The techniques used for measuring the proportions of the maxilla, dentary, ilium, humerus, femur, and metatarsals are presented in Figure 8.9 in P. Larson (2008). None of these measurements were challenged by Carr et al. (2022) and some were utilized by them. In this work new cranial measurements of the 16 large *Tyrannosaurus* skulls are added (Table 1), the techniques for measuring the proportions of the maxilla, the lacrimal, the postorbital process of the jugal, and the posterior ramus of the quadratojugal are illustrated in Figure 8 in direct association with the presentation of those results below. Anteroposterior dimensions of the bases of anterior dentary teeth, largely from P. Larson (2008) and especially Paul et al. (2022), on a given side are from either the teeth themselves, or the alveoli which are usually barely larger. The raw femur measurements are graphically plotted to allow visual comparison of variations in the variability of the proportions between tyrannosaurid taxa; the measurements for all the elements sampled are converted into ratios and stratigraphically correlated to the extent possible, and the results graphically plotted to reveal any resulting evolutionary patterns over time (as per Scannella et al. 2014); related statistical work is explained and presented in Paul et al. (2022), or in preparation.

**Table 1.** Upper skull ratios and related data for large Tyrannosaurus specimens, measurements in mm. Specimens approximately ordered by gross stratigraphic level in the TT-zone, and species assignments, to the degree possible, with robust, gracile or borderline as determined by overall skeletal analysis, largely based on results from Paul et al. (2022). Maxilla length/height rations from Paul et al. (2022) except for additions (UWBM 99000 740/394 mm, RGM 792.000 781/422 mm). All skull lengths are approximate and there are uncertainties about some; more readily measured femoral lengths provide generally more reliable comparative sizes of the individuals. Additional calculations (variation percentage, ratio ranges, ratio averages and medians) for the large specimens of each taxon are at the bottom of each taxon’s data set. Abbreviations -- postorbital boss prominence rankings; not prominent (NP), fairly prominent (FP), prominent (P), and very prominent (VP): nasal ridge rugosity ratings; smooth (S), fairly rough (FR), rough (R), very rough (VR), extra rough (ER): element is too incomplete in at least one dimension or otherwise not measurable or estimable (nm): skull length is not reliably restorable (nr).

|  | Species | Grac<br>or<br>Rob | Level | Skull<br>Length | Femur<br>Length | Max<br>L/D Ratio | Max<br>Fenestra<br>Width | Min<br>Pillar<br>Width | MF/<br>MP<br>Ratio |
| --- | --- | --- | --- | --- | --- | --- | --- | --- | --- |
| NHMAD “Stan” | *T. reg.* | g | h | 1470 | 1350 | 1.96 | 127l | 33l | 3.85 |
| LACM 23844 | *T. reg.?* | g? | h | 1380 | na | 2.17 | 127r | 32r | 3.97 |
| USNM 555000 Wankel | *T. reg.* | g | h | 1360 | 1280 | 2.02 | 97l | 23l | 4.22 |
| MOR 980 Peck's-rex | *T. reg.* | g | h | 1360 | 1232 | 2.26 | 116r | 27r | 4.3 |
| TMP 81.6.1 Black Beauty | *T. reg.* | g | h | 1190 | 1210 | 2.19 | 103l 89r | 27l 24r | 3.76 |
| RSM P2523.8 Scotty | *T. rex* | r | h | nr | 1333 | nm | 118r | 50r | 2.36 |
| CM 9380 | *T. rex* | r | h | 1360 | 1269 | 1.83 | 110l | 33l | 3.33 |
| UWBM 99000 Tufts-Love | *T. rex* | r | h | 1300 | na | 1.88 | 124r | 40r | 3.1 |
| Z-rex/Samson | *T. imp.* | r | l | 1400 | 1343 | 1.87 | 97r | 48r | 2.02 |
| FMNH PR2081 Sue | *T. imp.* | r | l or m | 1410 | 1321 | nm | 112r | ~51r | ~2.2 |
| MOR 008 | *T. imp.?* | ? | na | 1400 | na | 2.06 | 81r | 25r | 3.24 |
| SDSM 12047 | *T. imp.* | r | l | 1400 | na | nm | 103l | 48l | 2.15 |
| HMN MB.R.91216 Tristan | *T. imp.* | r | l | nr | 1220 | nm | nm | nm | nm |
| RGM 792.000 Trix | *T. imp.* | r | l | 1300 | 1170 | 1.85 | 91l 125r | 43l 48r | 2.35 |
| MOR 1125 B-rex | *T. imp.* | r | l or m | 1230 | 1150 | 1.88 | 98l | 49l | 2 |
| AMNH 5027 | ? | ? | Na | 1370 | na | 1.97- | 115l 105r | 38l 36r | 2.98 |
| UCMP 118742 | *T. rex?* | ? | h | na | na | 2.08 | 113r | 44r | 2.57 |
| Range |  |  |  |  |  | 1.83-2.26 |  |  | 2- 4.3 |
| Median |  |  |  |  |  | 2.05 |  |  | 3.15 |
| Average |  |  |  |  |  | 2.17 |  |  | 3.03 |

|  | Lac Height | Min Lac Width | Lac H/B Ratio | Jugal Height | Jugal Width | Jugal H/B Ratio | Quad Height | Min Quad Width | Quad H/B Ratio |
| --- | --- | --- | --- | --- | --- | --- | --- | --- | --- |
| NHMAD "S" | 350l | 51l | 6.86 | 461l | 139l | 3.32 | 304l | 63l | 4.83 |
| LACM 23844 | 325r | 57r | 5.7 | na | na | na | nm | nm | nm |
| USNM 555000 Wankel | 284l | 55l | 5.16 | 398r | 119r | 3.34 | 265r | 66r | 4.02 |
| MOR 980 Peck's-rex | nm | nm | nm | nm | nm | nm | 250r | 52r | 4.81 |
| TMP 81.6.1 B.B. | 229l | 32l | 7.15 | 334r | 100l<br>113r | 3.14 | 199r | 54r | 3.69 |
| RSM P2523.8 Scotty | 372l | 55l 55r | 6.75 | nm | nm | nm | 278r | 71l 70r | 3.94 |
| CM 9380 | 380l | 80l | 4.75 | na | na | na | na | na | na |
| UWBM 99000 T.-L. | 310r | 47r | 6.6 | 397r | 142r | 2.8 | na | na | na |
| Z-rex/Samson | 338r | 69r | 4.9 | 408r | 157r | 2.6 | nm | nm | nm |
| MOR 008 | 378r | 87r | 4.34 | nm | nm | nm | nm | nm | nm |
| FMNH PR2081 Sue | 263l | 70l | 3.75 | 430r | 135r | 3.19 | 267l | 70l | 3.81 |
| SDSM 12047 | nm | nm | nm | nm | nm | nm | 210l | 61l | 3.44 |
| HMN MB.R.91216 Tris | 396l | 81l | 4.89 | nm | nm | nm | nm | nm | nm |
| RGM 792.000 Trix | 280l 346r | 55l 48r | 6.15 | 400l 413r | 130l<br>144r | 2.97 | 208l<br>235r | 55l 58r | 3.92 |
| MOR 1125 B-rex | 286l | 47l | 6.09 | 371l | 142l | 2.61 | 273l | 59l | 4.63 |
| AMNH 5027 | 318l 323r | 56l 65r | 5.25 | 438l | 164l | 2.67 | 274l<br>303r | 58l 56r | 5.06 |
| UCMP 118742 | na | na | na | na | na | na | na | na | na |
| Range |  |  | 3.75-7.2 |  |  | 2.6-3.34 |  |  | 3.44-5.06 |
| Median |  |  | 5.48 |  |  | 2.97 |  |  | 4.25 |
| Average |  |  | 5.6 |  |  | 2.96 |  |  | 4.22 |

|  | Species | Skull Length | Femur Length | Postorbital Boss Score | Postorbital Boss Rank | Postorbital Boss Shape | Nasal Ridge Score | Nasal Ridge Rank | Sex |
| --- | --- | --- | --- | --- | --- | --- | --- | --- | --- |
| NHMAD “S” | *T. reg.* | 1470 | 1350 | 8 | P | hat | 8 | R | M |
| LACM 23844 | *T. reg.*? | 1400 | na | 2 | NP |  | 7 | R | F |
| USNM 555000 Wankel | *T. reg.* | 1360 | 1280 | 9 | P |  | 9 | R | M? |
| MOR 980 Peck’s-rex | *T. reg.* | 1360 | 1232 | 7 | P |  | 4 | FR | M? |
| TMP 81.6.1 Black Beauty | *T. reg.* | 1190 | 1210 | 4 | FP |  | 1 | FS | F |
| RSM P2523.8 Scotty | *T. rex* | nr | 1333 | 13 | VP | knob | 11 | R | M |
| CM 9380 | *T. rex* | 1360 | 1269 | na | na | na | na | na | ? |
| UWBM 99000 Tufts-Love | *T. rex* | 1300 | na | 10 | P | knob | 14 | VR | F? |
| Z-rex/Samson | *T. imp.* | na | 1343 | 5 | P |  | 6 | R | F? |
| FMNH PR2081 Sue | *T. imp.* | 1410 | 1321 | 14 | VP | spindle | 5 | FR | M |
| MOR 008 | *T. imp.*? | 1400 | na | 15 | VP | spindle | 15 | FR | M |
| SDSM 12047 | *T. imp.* | 1400 | na | 12 | P | ~spindle | 10 | R | M? |
| HMN MB.R.91216 Tristan | *T. imp.* | nr | 1220 | 1 | NP |  | 2 | FS | F |
| RGM 792.000 Trix | *T. imp.* | 1250 | 1170 | 11 | P | spindle | 12 | VR | M? |
| MOR 1125 B-rex | *T. imp.* | 1230 | 1150 | 3 | FP |  | 3 | FR | F |
| AMNH 5027 | ? | 1370 | na | 6 | P |  | 13 | VR | ? |
| UCMP 118742 | *T. rex*? | na | na | na | na |  | na | na | ? |

The relative prominence of cranial display structures is comparatively rated by reproducing photographs to a same ornament bearing element anteroposterior length and cross comparing and ordering them until a progressive increase in the degree of prominence was placed in line of order and could then be numerically ranked (Table 1). Postorbital cranial ornaments are illustrated in lateral view by direct tracings of photographs of the structures and the surrounding element. The forms of the bosses are stratigraphically correlated in the text to reveal any patterns of evolution over time.

Skulls are illustrated by direct tracings of photographs and technical illustrations. Procedures for producing high fidelity profile-skeletals and using them to estimate body masses for tyrannosaurids and other tetrapods are detailed in Paul (1988, 1997, 2016, 2019; Larramendi et al. 2021).

The results of the collective analysis are tabulated towards the end of the section on *Tyrannosaurus* species.

## RESULTS AND DISCUSSION

### Current Standard Principles and Practices for Sorting and Diagnosing Paleospecies

#### Monospecificity in a Genus is Not the Automatic Null Hypothesis

Carr et al. (2022) contend that a single paleospecies is the null hypothesis for tetrapod fossil genera. The theory being that the simplest explanation should be presumed unless sufficient evidence indicates otherwise. This is simplistic and problematic for a number of reasons. Biology is normally complex, not simple. Specific to paleotaxa, a genus is always complicated in that it is inherently a collection of a number of species that express varying degrees of gradistic and phylogenetic evolution within the genus (Figs. 2-4), until gradistic differentiation via speciation and phylogenetic divergence add up and result in one or more new genera. In paleozoological taxonomy, the longer a genus is preserved in the known geological record and/or the laterally wider its distribution, the higher the probability that more than one species has been recorded over that time. That is all the more true if other genera preserved in the same sediments show substantial evidence of having undergone speciation. Of course if just one specimen is known from a genus that is just one species, but the more specimens known the higher the likelihood more than one species are on hand. The larger the number of collected specimens contained within a paleogenus both vertically and at a given time, the greater is the probability multiple species have been preserved in the sample.

**Fig. 1.**
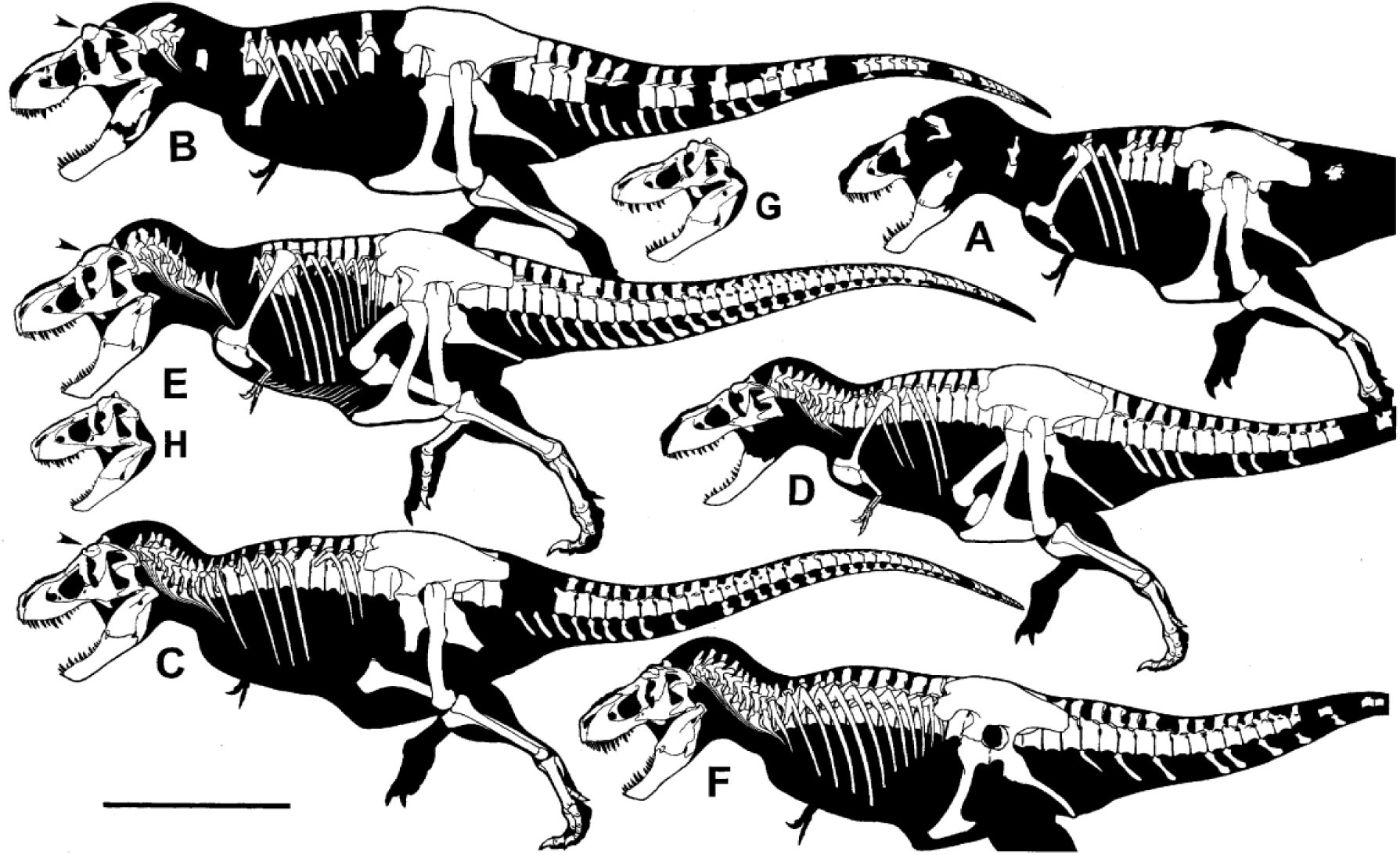
*Tyrannosaurus* known-bone profile-skeletals and skulls to same scale, bar equals 2 m, with skulls revised to varying extents from versions in Paul et al. (2022, Fig. 1) with particular attention directed to accurate postorbital bosses, arrows point to those of large presumed mature males (MM) of the three species. Skeletals: **A** Upper TT-zone *T. rex* holotype CM 9380 (6.5 tonnes); **B** Upper TT-zone *T. rex* RSM 2523.8 (MM 7.8); **C** Upper TT-zone *T. regina* NHMAD “S” (exBHI 3033) (MM, 7.5); **D** Upper TT-zone *T. regina* holotype USNM 555000 (immature? male? 6.1); **E** Lower TT-zone *T. imperator* holotype FMNH PR2081 (MM 7.8); **F** TT-zone unknown *T*. incertae sedis AMNH 5027, preservation of ribs uncertain for this specimen. Skulls: **G** Upper TT-zone *T. rex* UWBM 99000 (immature? female?); **H** lower TT-zone *T. imperator* MOR 1125 (immature? female).

**Fig. 2.**
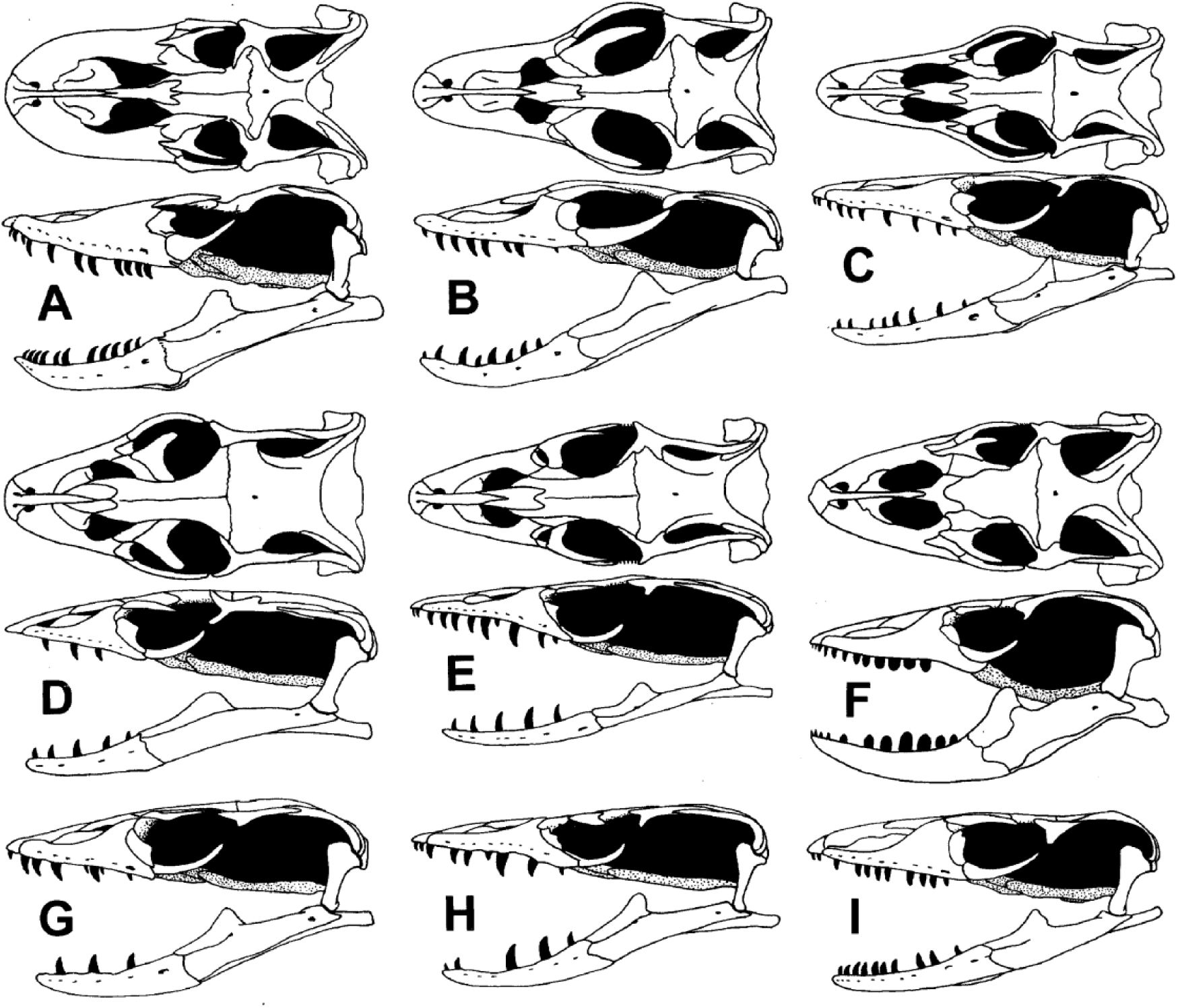
Same length comparisons of predatory reptile *Varanus* skulls in dorsal and/or lateral views. **A** *V. komodoensis*. **B** *V. griseus*, **C** *V. salvator*, **D** *V. gilleni*, **E** *V. prasinus*, **F** *V niloticus*, **G** *V. semiremex*, **H** *V. beccarii*, **I** *V. bengalensis*. Image source Mertens (1942).

In order to establish that a genus known from a substantial sample collected from sediments that were deposited over a span of hundreds of thousands of years or more, is monospecific, it needs to be shown that there is very little or no pattern of variation in anatomy compared to other species, or within the remains over time, and the basal condition is largely or entirely retained throughout the sample. If instead the observed variation is atypically high, and/or than the variation shows a significant pattern when stratigraphically correlated, then the multispecies hypothesis which is the normal condition of genera is superior to the monospecific alternative. All the more so if the shifts tend to be away from the ancestral state.

Taxonomic implications – The monospecific hypothesis is not automatically presumed superior over the multispecific alternative, if anything the opposite is more likely, especially if a considerable span in time and/or geography is on hand in terms of sediments bearing the fossils of the genus under taxonomic examination. Rather than loading the scientific dice with one or another presumption that can cryptically bias the results, the preponderance of cumulative evidence for whatever results best explain the best current data needs to be the predominant means for arriving at systematic conclusions.

#### The Varying Amount of Variation Between Species Within Genera

The amount of morphological skeletal variation between species contained in closely related paleogenera can be very substantial. For example, although the giant, early Maastrichtian Asian tyrannosaurid *T. bataar* was initially considered a species of the markedly later Maastrichtian North American *Tyrannosauru*s (Maleev 1955; as some continue to consider plausible [Paul 1988, 2016; Carr et al. 2005]), it is usually considered to be the one known species of *Tarbosaurus* (Brochu 2003; Hurum and Sabath 2003; Loewan et al. 2013). It is often thought that *Tarbosaurus* and *Tyrannosaurus* are close relatives relative to other tyrannosaurids (Carr et al. 2005; Paul 2016; Loewen et al. 2013), although it is possible that their gigantism caused them to parallel one another (Hurum and Sabath 2003; Paul et al. 2022 Suppl.; this is highly plausible in view of the extensive amount of convergence and parallelism in vertebrates between geographical areas [Oyston et al. 2022]). As documented in the systematics section *Tarbosaurus bataar* is readily distinguished from the collection of *Tyrannosaurus* species by a number of features including relatively less massive and more bladed teeth, less extremely broad temporal region of the skull, lacrimals that do not nearly contact one another along the midline of the skull roof, a smaller pubic boot, longer lower hindlimb elements, and other details. That the two taxa are so distinct is not surprising because, well separated in time, geography, and to a certain extent in phylogeny, they are not very close sibling species in an unambiguous single genus. Much closer in form to one another are *Gorgosaurus libratus* and *Albertosaurus sarcophagus* which differ only in some barely noticeable skull features (Paul 1988, 2008, 2016; Currie 2003a).

Within a genus the relationship between species ranges from very close to more distant, so interspecific skeletal variation tends to conversely range from substantial to very minimal at best. *Varanus* contains a large number of extant and recently extinct species (Fig. 2). Size ranges from small to gigantic (with *Megalania priscus* subsumed into *Varanus* [Molnar 2004; Head et al. 2009]), teeth from serrated cutting blades to bulbous crushing teeth, skulls from fairly narrow to quite broad, skull roof suture patterns differing markedly, as can the robustness of postcrania. On the other hand some *Varanus* species skeletal details are barely distinguishable (Fig. 2C,E,H). *Panthera* contains a number of extant and recently extinct species (Fig. 3E-H). Their skulls and skeletons range from significantly divergent, the snow leopard *P. uncia* sports an extra-long tail, to too difficult to osteologically segregate (Sotnikova and Nikolskiy 2006; Christiansen and Harris 2009; Fig. 3I,J). One of the few features that distinguishes the skulls of lions *P. leo* and tigers *P. tigris* is the convex ventral rim of the mandible of the former compared to the flatter configurations found in the tiger and other *Panthera*. Sotnikova and Nikolskiy (2006) observe that the cave lion *P. spelaea* and its probable descendent the American lion *P. atrox* are barely distinguishable from one another, including flat bottomed mandibles (Fig. 3E,F). While the robust timber wolf *Canis lupus* (Fig. 3B) is not difficult to tell apart from the Ethiopian *C. simensis* and African *C. anthus* wolves (Fig. 3C,D), the latter two and other gracile *Canis* are not easy to tell apart. The overall species situation with the genus and its close relatives is often problematic (Koepfli et al. 2015; Alvares et al. 2019; Perri et al. 2021), with taxa that are morphologically not distinguishable in their skeleton and even soft tissues appear to be different species genetically. It is interesting that while the skeletons of the Eurasian origin timber *C. lupus* and dire wolves *C. dirus* are so alike that they were long considered sibling species (Fig. 3A,B), molecular work indicates that the latter is of quite separate American origin (Perri et al. 2021).

**Fig. 3.**
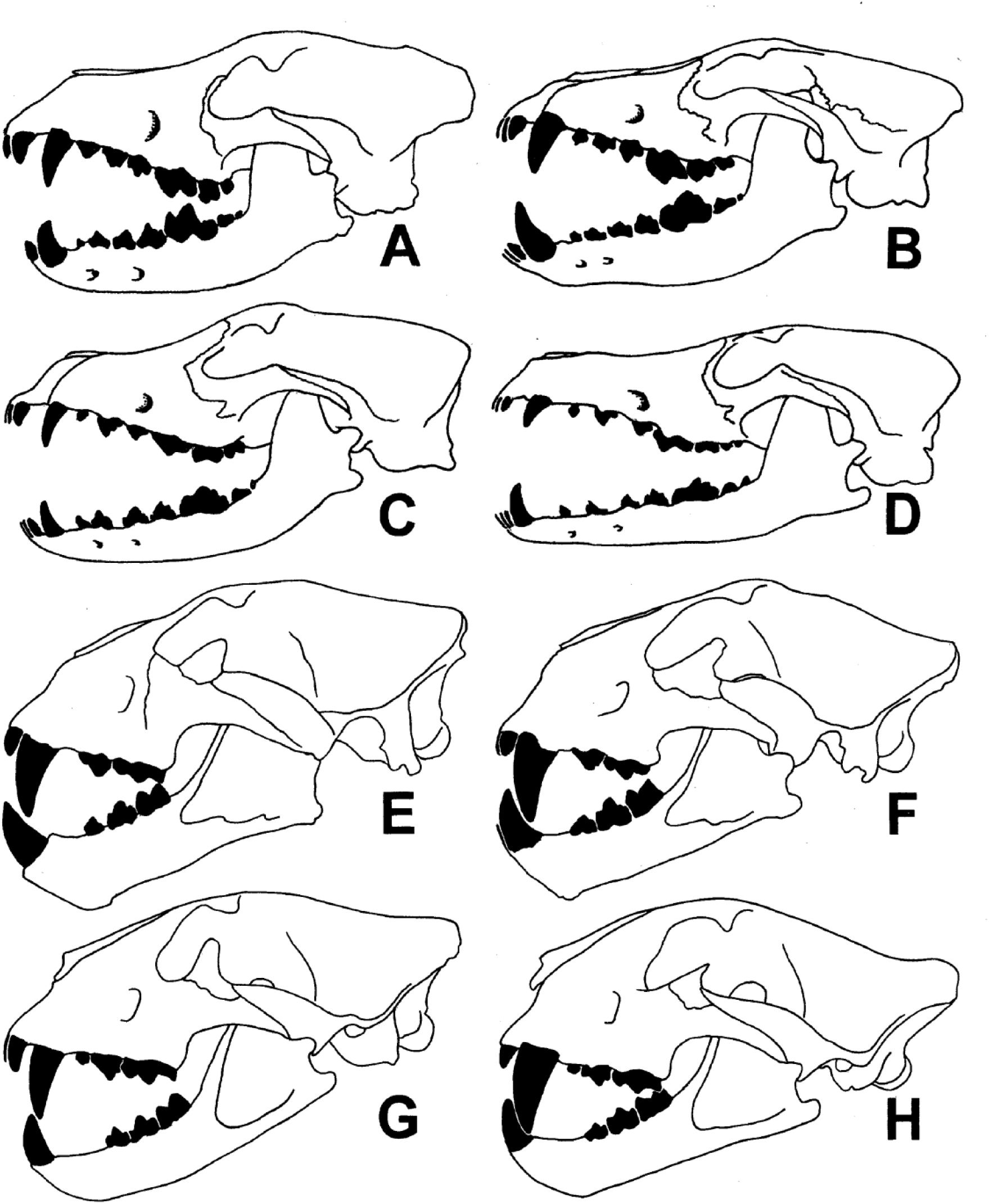
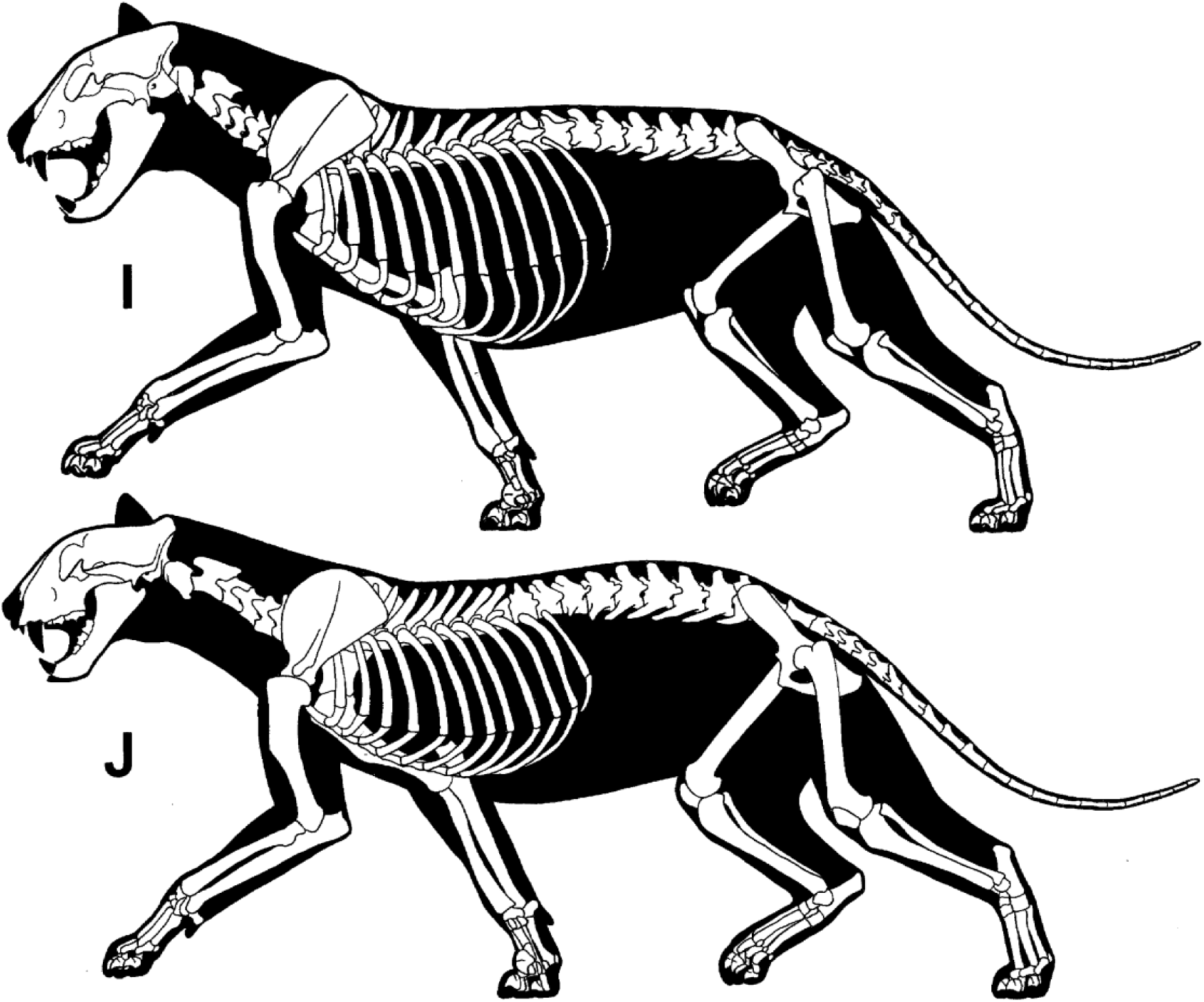
Same length comparisons of predatory mammal skulls. *Canis*: **A** *C. dirus***; B** *C. lupus***; C** *C. anthus*; **D** *C. simensis*. *Panthera*: **E** *P. spelaea*; **F** *P. atrox*; **G** *P. leo*; **H** *P. tigris*. Same length comparisons of *Panthera* skeletons: **I** *P. leo*; **J** *P. tigris*.

Taxonomic implications -- The degree of differentiation between species is to a great extent a function of the degree of phylogenetic and therefore gradistic divergence between species. Because sibling species that have just diverged from one another are barely anatomically distinct from one another, it may be just one reasonably consistent character that distinguishes them. Large character sets are only expected when two species are evolutionarily spaced out and divergently evolved by a substantial number of intervening species.

#### How Many Hard Bone Characters Major and Minor Are Needed to Diagnose Sibling Paleospecies When a Good Fossil Sample is On Hand

It follows that as detailed in Paul et al. (2022 Suppl.) in modern peer reviewed literature as few as one skeletal character, often minor, has been used to separate species extant and fossil known from well documented remains often including complete skulls and skeletons, with many cases involving just two, three or a few features, among continental and marine tetrapods, with proposed species sometimes being contemporary, or at least partly separated stratigraphically (tyrannosaurids -- Currie 2003a; ornithopod *H. multidens* – Barrett et al. 2005; *Panthera spelaea, P. atrox* -- Sotnikova and Nikolskiy 2006; hadrosaurs *Corythosaurus, Lambeosaurus* -- Evans and Reisz 2007; ichthyosaur *Stenopterygius* -- Maisch 2008; Maxwell 2010; brontotheres *Eotitanops, Palaeosyops, Metarhinus, Rhinotitan,* -- Mihlbacher 2008; Mader 2010; *Psittacosaurus* – Sereno 2010; plesiosaur *Pliosaurus* – Knutsen 2012; diplodocids *Apatosaurus, Brontosaurus, Diplodocus* -- Tschopp et al. (2015); chasmosaurines - Campbell et al. 2016; Fowler and Freedman 2020; ornithomimids *Ornithomimus*, *Dromiceiomimus* -- MacDonald & Currie 2018; proboscideans *Mammut, Palaeoloxodon* -- Dooley et al. 2019; Larramendi et al. 2020).

In a reversal of the proposition that species best documented by fossil remains should be distinguished by lots of characters, the *Psittacosaurus* known by the largest number of specimens, *P. mongoliensis*, is distinguished from sister species by the least number of attributes, two, in Sereno (2010). Both characters are minor, consisting of a raised lip on the orbital margin of the prefrontal, and the transverse distal expansion of the ischial blade being about twice the width of the midshaft. For *Mammut pacificus* the diagnostic characters are a molar being unusually narrow, six rather than five fused sacrals, an exceptionally robust femur, no mandibular tusks, and smaller primary tusks (Dooley et al. 2019). The characters diagnosing the species of *Stenopterygius* are variations in the degree of tooth reduction, subtle proportions of the body and fins, skull/body length ratios, and size of a distal tarsal relative to other ankle elements (Maxwell 2012). For *Pliosaurus* the differences are tooth counts, the configuration of the retroarticular process, ventral anatomy of cervicals, and differing limb proportions (Knutsen 2012). None of these examples has excited strong negative reaction including in the news media of the sort towards the splitting of the tyrant lizard king in their immediate wake.

Ironically, the use of very large skeletal character data sets to discern paleospecies involves its own set of issues. For a number of practical reasons including logistical assembling character lists involving many hundreds of cranial and postcranial attributes in a large number of sufficiently complete fossil specimens within a genus is rarely if ever tried much less achieved. One discouragement is that the utility of so much data is problematic. Because sibling species are so similar in most regards, large character sets risk producing a large amount of useless and potentially misleading background noise -- just because 99.9….% of the anatomy of a set of intragenus skeletons is indistinguishable does not mean they are all in the same fossil based species. That is a reason why sibling paleospecies research tends to focus on discerning the limited number of characters that do vary between specimens, a far greater number of details are not tallied they being irrelevant. If a very large number of characters are assessed it becomes difficult to clearly illustrate what was examined and how, making verification and reproducibility increasingly difficult as the character and specimen list piles up. All the more so because the time and effort needed to verify all the character assessments may be very difficult at best to replicate. Using a very large character set to assess species within the only genus that such is available for is provisional, in that the method as not yet been tested in terms of practicality, utility and reliability on a wide basis. To demonstrate the efficacy of the method it would be necessary to employ it on genera the species of which are well established, including their stratigraphy. Examples would be *Panthera* that includes a number of well documented recent and extant species. *Mammuthus* would also be well suited. For dinosaurs *Triceratops* features a large number of specimens whose geology is established or will be, but the genus lacks a large sample of necessary postcrania. Lancian edmontosaurs have the latter, but not yet the stratigraphy (Paul et al. 2022). The dinosaur genus sporting the most species, *Psittacosaurus*, lacks postcrania in some examples, and tight stratigraphy. Tschopp et al. (2015) examined 455 characters in a modest number of usually skullless diplodocid fossils, but in the end the numbers actually distinguishing the widely accepted species came down to just a handful each. Testing the mass character method in dinosaurs may not be possible at this time.

Carr et al (2022) do not agree with earlier media suggestions that intrageneric paleospecies require large character sets.

Taxonomic implications – Just one minor character can diagnose a sibling intrageneric species, and the use of hundreds of characters to assay such may not be as efficacious as may be thought.

#### Strong Consistent Bimodal Character Separation is Not Universal Among Species Within a Genus

Carr et al. (2022) indicate character bimodality is important if not critical to species designation and diagnosis. However, as noted in Paul et al. (2022), characters used to differentiate and diagnose species are often not bimodal and nonoverlapping in distribution (Maisch 2008; Maxwell 2012; Scannella et al. 2014; MacDonald & Currie 2018; Harvati & Ackermann 2022), and statistical, measurements based bimodality is often not even presented in defense of paleospecies (Mihlbacher 2008; Mader 2010; Sereno 2010; Knutsen 2012; Tschopp et al. 2015; Fowler and Freedman 2020; Johnson et al. 2020). Carr et al. (2022) ignore these numerous examples that contradict their view. Bimodality is less likely to occur when more than two species are under examination – while two of the species may exhibit considerable bimodality, a transitional or sibling species may muddy those clear-cut taxonomic waters. So can hybridization between closely related species (Barnosky and Bell 2004; Lister and Sher 2015; Harvati & Ackermann 2022).

The promotion by Carr et al. (2022) of intrataxa consistency and intertaxa nonoverlap of characters used to diagnose taxa is any case inconsistent on their part, because they use maxillary tooth counts to distinguish *Tyrannosaurus* from *T. bataar* even though the numbers vary in the adults of both taxa and sometimes overlap (Osborn 1905, 1917; Maleev 1955, 1974; Brochu 2003, Hurum and Sabath 2003; Carr 2020).

Taxonomic implications – Strong interspecific bimodality and intraspecific consistency in statistical character results is a paleospecies ideal by no means always achieved. Strong trends are often used.

#### Reptile Teeth Can Have Diagnostic Value at the Species Level

The typically simple teeth of reptiles do not contain the intricate taxonomic information common to the complex dentitions of most mammals. But if the tooth characteristics within adults of a reptile genus exhibit of reasonably consistent pattern of differentiation between potential species, then they can be used to help diagnose intragenera paleospecies. Such is common in *Varanus i*n which a number of species have blunt crushing teeth (Fig. 2F) that distinguish them from species with more or entirely flesh slicing serrated blades (Fig. 2A-E,G-I). For that matter teeth help distinguish tyrannosaurid genera, with the more conical teeth of *Tyrannosaurus* marking it from the rest of the more blade toothed members of the family (Russell 1970; Paul 1988; Hurum & Sabbath 2003; Carr 2020). *Stenopterygius* species are diagnosed in part on differing degrees of tooth reduction (Maxwell 2012), those of *Pliosaurus* by tooth counts (Knutsen 2012).

Taxonomic implications – Although not as systematically valuable as is often true of intricate mammal teeth, reptile teeth can help determine the number of species within a genus. This utilization is often limited to large individuals, in order to minimize ontogenetic alterations in tooth forms and counts with ontogeny. Carr et al. (2022) do not object to use of reptilian teeth in the process of assaying theropod species.

#### Variations in Skeletal Strength and Other Proportions Can Have Diagnostic Value at the Species Level

Robustness of the skull and skeleton has important functional implications, and may have additional reproduction associated identification and competition attributes. The latter may be especially true in predators that are less able to use prominent bone based display features. Differences in build that evolve for functional purposes can be secondarily exploited as visual species identification cues and reproductive competition without the development of elaborate display devices. The brontotheres species *Metarhinus abbotti* and *M. fluviatilis* are diagnosed only the robustness of their anterior nasals (Mihlbacher 2008; Fig. 4). *Homo sapiens* and *H. neanderthalensis* are distinguished in part on the greater skeletal strength of the latter. Differing robustness is used to help diagnose mastodon and mammoth species (Dooley et al. 2019; Larramendi et al. 2020). Skeletal robusticity, especially that of the femur, has been used to help distinguish the tyrannosaurids *Daspletosaurus torosus* from *Gorgosaurus libratus* that share the same habitat (Russell 1970; Paul 1988; Currie 2003b; Snively et al. 2006). Currently *Ornithomimus edmontonicus* and *Dromiceiomimus brevitertius* are morphologically differentiated by their different femur tibia ratios (MacDonald & Currie 2018).

Taxonomic implications – Carr et al. (2022) do not a-priori object to the use of bone robustness to help examine and diagnose paleospecies.

#### How Critical a Role Do or Do Not Differences in Prominent Display Characters Play in Distinguishing Species, Particularly in Predators

An important feature of species is a reproductively isolated population, although this is not an absolute (Mayr 1982; Zeinio 2012). It has therefore been argued that visual display structures commonly evolve for purposes of species identification in order to achieve reproductive isolation, including among dinosaurs (Padian and Horner 2011, 2014). Others disagree (Hone and Naish 2013; Knapp et al. 2018), and it is notable that African white and black rhinos living in the same habitats share broadly similar horn arrangements, and are otherwise not strongly divergent in appearance. At the same time, it is true that species are not prone to sport strongly divergent display features regarding their basic shape, particularly between males of a given species, although their degree of development may vary due to dimorphism, maturity, and individual variation. Reasonably consistent differences in bone based cranial displays are therefore widely used to help sort out paleospecies including dinosaurs whether they share a habitat or not (Evans and Reisz 2007; Mihlbacher 2008; Scannella et al. 2014; Campbell et al. 2016; Paul 2016; Fowler and Freedman 2020).

Continental herbivores often evolve elaborate display devices, frequently formed from horns, antlers and tusks that may have evolved at least in part as anti-predator weapons (Nowak 1991), as well as crests that may have no other important function than display, such as the those of the modest set of birds that have evolved bony cranial displays (Hoyo et al. 1992, 1994, 1996, 1997, 2001; Mayr 2018). Some dinosaur taxa, such as the lambeosaurinian species in *Corythosaurus, Lambeosaurus* and others (which should be in the same genus according to Paul 2016), are not significantly distinguishable cranially outside their display structures or postcranially (Fig. 5A-C), it’s all in the crests. A similar situation applies to the centrosaurinian species in *Centrosaurus, Styracosaurus* and others (Fig. 5E-G; Paul 2016), it is a matter of horns, spikes and bosses. Because herbivores commonly evolve large cranial structures in part for defense, these display features can then be modified into very distinctive shapes, even between sibling species. This is true in the primary prey of *Tyrannosaurus*, *Triceratops*, in which the differences in the nasal horns of *T. horridus* and its probable descendent *T. prorsus* are readily visible – however the differentials between those two species and the intermediate taxon, and between the postorbital horns, are less so as discussed below.

**Fig. 4.**
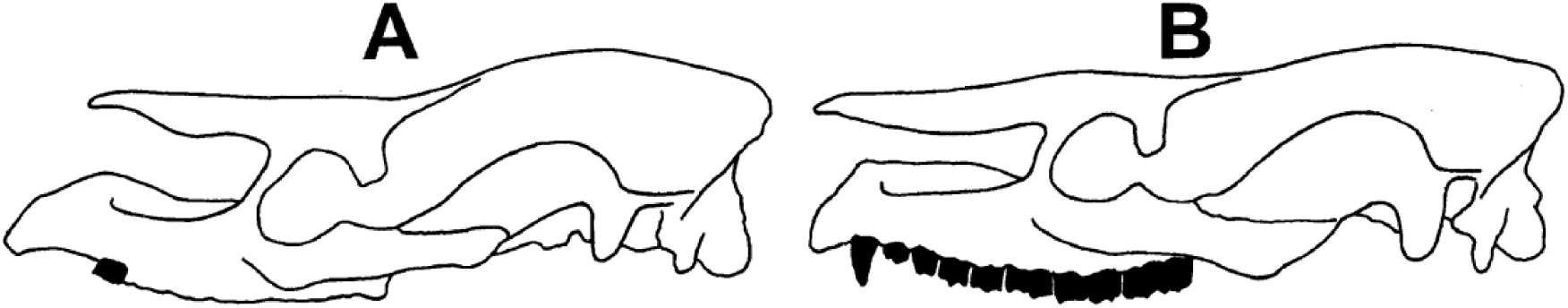
Same length comparisons of brontothere *Metarhinus* skulls. **A** *M. abbotii*; **B** *M. fluviatilis* (posterior skull somewhat dorso-ventrally crushed).

**Fig. 5.**
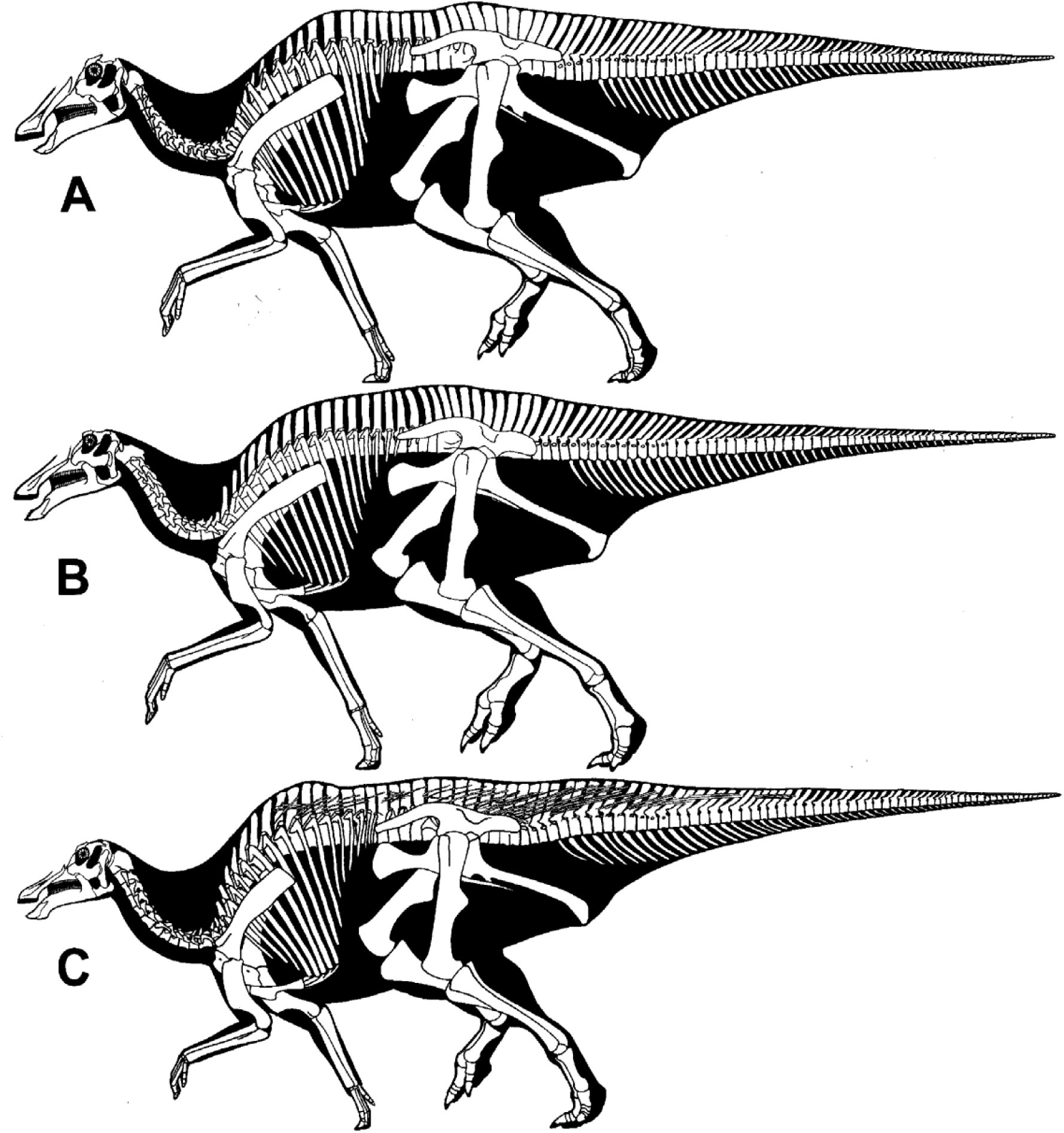
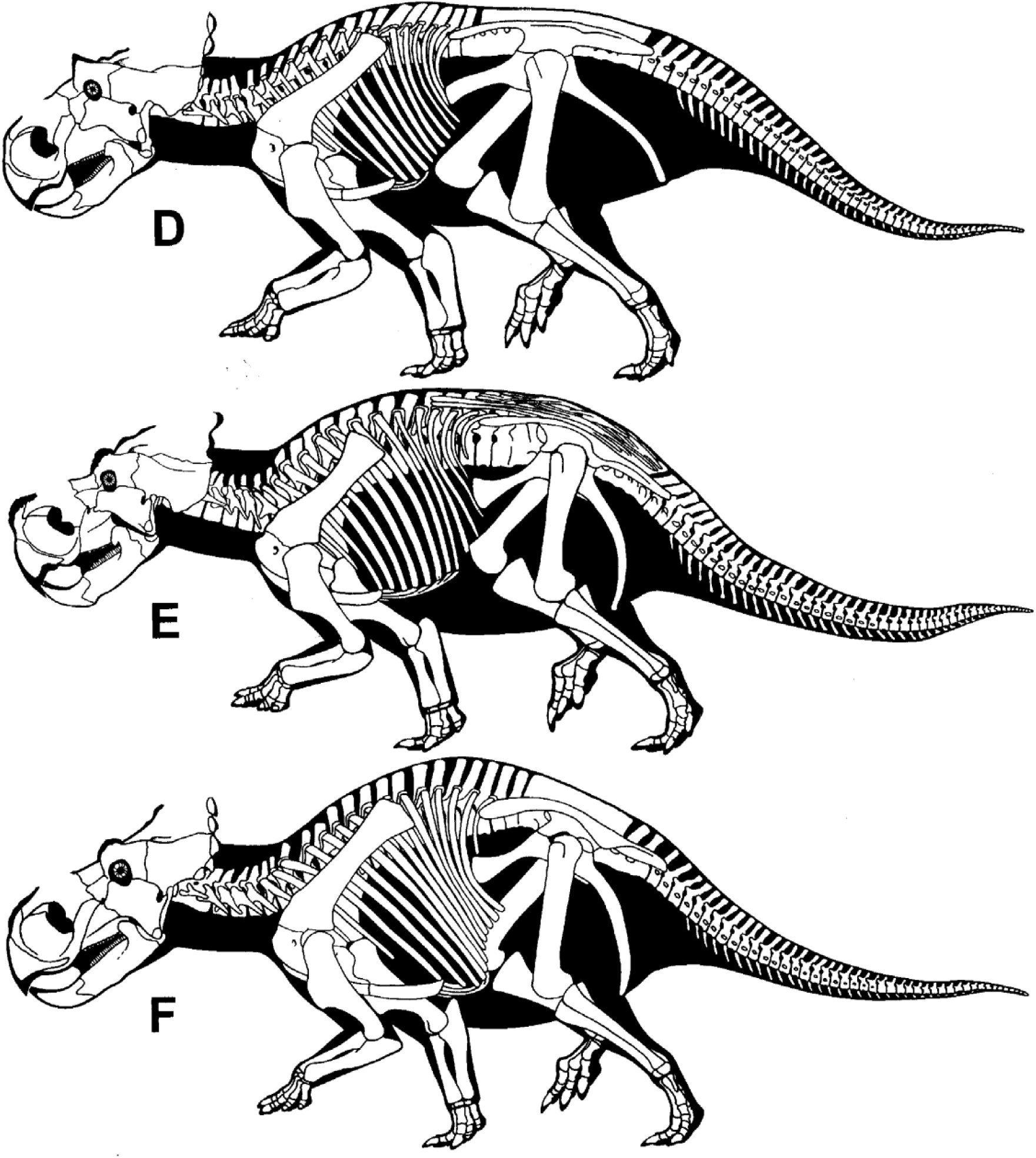
Same length and approximate same scale comparisons of the skeletons of Campanian North American herbivorous dinosaurs sans cranial species specific display structures. Lambeosaurinians: **A** ROM 1218; **B** ROM 845; **C** AMNH 5240; testing ability of viewers to tell which are *Corythosaurus casuarius*, *C. intermedius* and *Lambeosaurus lambei* without their crests. Centrosaurinians: **D** YPM 2015; **E** AMNH 5351: **F** AMNH 5372; testing ability of viewers to tell which are *Styracosaurus albertensis, Centrosaurus nasicornis* and *C. apertus.* Full skeletals in Paul (2016).

Predators are often another matter. Not bearing large defensive weapons that can be readily utilized and differentiated for display purposes, while using other skeletal elements such as cranial crests solely for display risks being a hindrance to predatory combat, bone based display features are often minimal or absent in land predators. There are no notable sexual display characters decorating the crania and postcrania of *Varanus, Panthera, Canis* and a host of other predaceous tetrapods (Figs. 2,3). Living lions and tigers are easy to visually distinguish because of major differences in fur and coloration, but their skeletons are difficult to tell apart (Figs. 3G-J). In many examples skeletal features used to help distinguish species are not sexual display adornments. Among the few predatory groups to have exhibited a fairly frequent propensity towards evolving major cranial display features in the form of ridges, crests, bosses, hornlets and short horns are nonmaniraptor avepods including some basal tyrannosauroids, but tyrannosaurids were limited to very modest structures (Paul 2016).

Reducing the sexual selective requirement for significant alterations in display organs can be the evolution of chronospecies. Because the two species do not meet at the same place and time, there is not a need for reformation of display structures even when they are present. This appears to have been true of *Gorgosaurus libratus* and its possible direct descendent *Albertosaurus sarcophagus*, the dorsal display features adorning the nasals, lacrimals and postorbital are not diagnostically distinguishable between the two closely related tyrannosaurid taxa (Fig. 6A,B). The skeletal adaptations used to diagnose the taxa are largely limited to characteristics of the basal braincase (Currie 2003a).

**Fig. 6.**
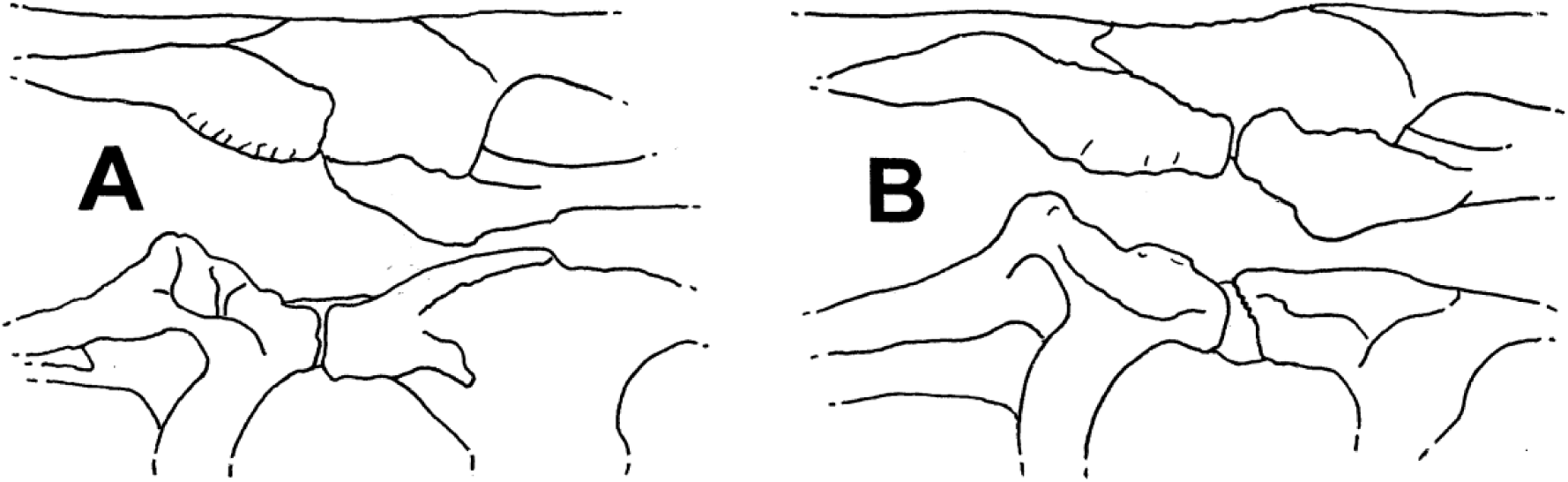
Albertosaur left lacrimal ridges and postorbital bosses in lateral (top) and dorsal (bottom) views to same length. **A** *Gorgosaurus libratus* TMP 91.36.500; **B** *Albertosaurus sarcophagus* TMP 81.10.1.

All the above said, the presence of distinctively divergent optically obvious display characters is strong evidence for the existence of two or more species in a genus. Such is characterized in sibling paleospecies by easily seeable difference in the shape and perhaps the size of readily visible bone display structures, usually cranial. The presence of a single distinctive difference in a display character is sufficient to distinguish and define sibling species from one another even if there are no other differences in the morphology of the specimens.

Complicating matters is how variation dimorphic, individual and ontogenetic can produce background noise (Hone and Naish 2013; Knapp et al. 2018) that the requires appropriate analysis to sort out species specific display characters. In dimorphic species, when females possess species exclusive displays in the form of cranial projections, they tend to be at least basically similar to those of the males in form (as per Nowak 1991, Hoyo et al. 1992, 1994, 1996, 1997, 2001; Mayr 2018) in order to facilitate species identification – having grossly different cranial displays on the two genders in one species risks species ID confusion. In the great majority of cases the more elaborate and/or larger display organs adorn the males, so that is the general null hypothesis. The difference is size and shape between the intraspecific genders range from minimal to substantial. Cassowaries are an exception in that female crests are generally more prominent, which may be related to how males provide most of the parental care (Hoyo et al. 1992). Differentiation of display structures may be dependent on complex qualitative shape variations that are not amenable to quantitative analysis such as simple orientation or size dimensions, but this does not preclude their critical importance.

Also complicating the situation in fossils is that bony cranial ornaments were covered by keratin sheaths that at least have the potential to alter their appearance relative to their skinless appearance. Among lizards cranial prominences are ensheathed in rather thin, shape conforming keratin (Vickaryous et al. 2015; Marghoub et al. 2022). Bird crests small and large are usually covered by thin keratin sheaths according to what data is available (Richardson 1991; Gamble 2007; Naish and Perron 2016). An exception is the crest of the helmeted hornbill *Rhinoplax vigil* in which the anterior portion of the casque is extended by an ivory like keratin about two centimeters thick that can be carved (pixels.com/featured/helmeted-hornbill-skull-natural-history-museum-londonscience- photo-library.html; Mayr pers. comm. pointed this out to me), but the gross form of the crest is still not radically altered. The best preserved direct example of nonavian dinosaur cranial display soft tissues, those on the modest sized hornlets of an ankylosaur, indicate that the keratin significantly but not greatly enlarged the ornamentation, by about a fifth to perhaps a third, and retained the basic shape of the underlying bone (Brown et al. 2017). It is presumed the same was true of the similarly modest sized cranial structures of tyrannosaurids. The soft tissues that sometimes greatly enlarge the transversely flattened midline crests of pterosaurs (Paul 2022a) do not appear applicable to the lower lying, lateral projections of tyrannosaurid crania. Same regarding the entirely soft tissue midline crests atop edmontosaur hadrosaur crania (Paul 2016).

Taxonomic implications – It is common, especially among terrestrial predators, for sibling species to not exhibit differences in bony sexual display structures, either because they are not present, or are not different. Establishing consistent variation in such display organs is not necessary to distinguish and diagnose paleospecies. But if the latter are present, then the species based nature of the divergences is solidly established.

#### Individual Variation Has Very Limited Explanatory Power for Evolutionary Trends

Random individual variation is not a selective force that drives evolution in a direction that is well off the norm for an anatomically uniform group. Ergo, individual variation is a primary causal explanation for variation within a paleogenus only when the observed variation exhibits little or no pattern over time. If a documented pattern of change in one direction or another does exist, then citing individual differences as a cause is at best an idle fallback position of non-scientific opinion that lacks supporting evidence or any cogent evolutionary explanatory power. That is all the more true if the observed variation in a genus exceeds that previously observed in its close relations, including the entire family outside the genus. Also working against individuality is when the pattern over time moves the contents of the fossil genus away from the basal/ancestral condition in one or especially more ways. Nor is genetic drift within a species an optimal causal explanation when the fossils are located in the same core region because there is not a coherent causal case of geographic isolation involved – and if the changes occur over a wide geography then the drift is likely to result in classic drift speciation (Mayr 1982).

Taxonomic implications – Genetic evolution via speciation is the superior, positive, go-to hypothesis when the fossil record reveals a distinct pattern of directional change, especially when it is away from the ancestral condition.

#### Sample Sizes Do Not Need to Be Large, nor is Deep Statistical Analysis Necessary, to Designate Regularly Used Intragenera Paleospecies

*Psittacosaurus i*s widely accepted to contain a large number of species with some consisting of a few or just one specimen (Sereno 2005). The number of specimens placed within two species of *Apatosaurus* is four, within three species of *Brontosaurus* is three, in two species of *Diplodocus* it is five; most of these specimens lack skulls (Tschopp et al. 2015). MacDonald and Currie (2018) used about two dozen ornithomimid specimens to statistically parse out species of *Ornithomimus* and *Dromiceiomimus*, with nine actually pertaining to the two genera. The species of *Lambeosaurus* have been determined by about a dozen large specimens (Dodson 1975; Evans and Riesz 2007). Scannella et al. (2014) state that over 50 specimens were examined, but the number of specimens that are both stratigraphically correlated and the measurements of which are statistically analyzed is about three dozen, some of which are juveniles.

Papers dealing with paleospecies that have not deployed extensive number crunching statistical analysis if any of a large sample include Barrett et al. (2005), Sotnikova and Nikolski (2006), Evans and Reisz (2007), Mihlbacher (2008), Mader (2010), Maxwell (2010), Sereno (2010), Knutsen (2012), Tschopp et al. (2015), Dooley et al. (2019), Fowler and Freedman (2020) and Larramendi et al. (2020).

Taxonomic Implications – While more is always better when it comes to science, a large array of tetrapod sibling paleospecies are named without statistical analysis, and/or based on just one or a few specimens, such being a normal practice. Although in principle such paleospecies are provisional until large samples become available, in practical terms many low specimen based intragenera paleospecies are widely accepted and utilized. Paleozoology regularly works with the data on hand, not what is wished for.

### Stratigraphic Correlations Do Not Need to Be Precise

Carr et al. (2022) assert that stratigraphic positioning needs to be “accurate” and “precise” for the purposes of paleospecies assignment. The first is correct, the latter is not because the first is not dependent on the other in the context of paleospecies determination. And shortly then after Carr et al. (2022) note that taxa can indeed be sorted into broad stratigraphic bins that are taxonomically informative. That is true because unlike many areas of science biology is almost always sloppy and fuzzy. In particular, fine stratigraphic resolution within formations such as that utilized by Scannella et al. (2014) are not required for paleospecies determination because fossil species are prone to last for hundreds of thousands of years (Gould 2002; Burger et al. 2004; Maisch 2008; Scannella et al. 2014; Hunt et al. 2015; Long et al. 2020). Therefore, all that needs to be known with confidence is at what gross level – lower, upper, middle – a given specimen is from. With formation sublevels usually being many tens to hundreds of meters thick, precisely how many meters a specimen is from the top or bottom of the geological unit is usually not vital. Knowing the sublevel of specimens only generally and not necessarily precisely is commonly followed in intrageneric paleospecies studies, Maisch (2008) being an example.

Taxonomic implications – Although the more precision the better is true in science, perfection is the enemy of good enough for what is on hand when that gets the basic job done. Because tetrapod species are prone to exist over significant geological time, the stratigraphic measures of that time do not need to be more precise than overall sublevels.

### Stratigraphic Correlations Do Have Great Utility

Stratigraphic correlations are not always necessary to designate sibling species, sometimes it is not available (as is often the case regarding *Psittacosaurus* species, note imprecision of stage levels cited in Paul 2016), but they are likely to be very useful when the data is present. Works that center on stratigraphic correlations include ichthyosaurs (Maisch 2008; Maxwell 2010); brontotheres (Mihlbacher 2008; Mader 2010), ceratopsids (Scannella et al 2014; Campbell et al. 2016; Fowler and Freedman 2020).

That the probably permanent inability to determine the basic stratigraphic position in the Hell Creek Formation, much less its precise vertical level to within a few meters, of *Tyrannosaurus* AMNH 5027 is an important reason its species status cannot be assigned, is a sterling example of how stratigraphy is very important, but precision not so much.

Taxonomic implications – Paleospecies studies need to incorporate as much geological information as is possible – but no need to go overboard on it.

### Taxonomic Floaters

Carr et al. (2022) emphasize that for a paleospecies diagnosis to have practical value, it must produce consistent results in identifying incomplete, but not necessarily fragmentary, remains. No references in support of this position are cited. The premise is problematic in view of how it is difficult in respect to some extant genera, such as *Canis*, to assign some members of the genus to specific species (Grubb et al. 2000). A substantial number of articulated dinosaur specimens, including some tyrannosaurids, from the Dinosaur Park Formation, have not been assigned down to the species level as per Currie and Russell (2005). Forcing the species of a stratigraphic zone to be one has the advantage of allowing all specimens of that genus to that species, but that may be a false convenience if in doing so results in the evidence for more than one species being played down when more than one was actually present.

Conversely, as long as confidence that only one species is present in a particular level of a formation is well founded, then assigning partial specimens assignable to the parent genus to the known species when the remains lack diagnostic features, but do not possess contrary characters, is the acceptable norm, Galton (1981) being an example.

Taxonomic implications – The goal should not be to diagnose paleospecies in a manner designed to maximize the ability to assign specimens to a species. Diagnoses should be a best effort to characterize the paleospecies that did exist and let the specimens fall where the data on each favors including indeterminate. The more closely species in the sediments are related to one another to more likely it will be difficult to place specimens that are either so incomplete they lack diagnostic structures, and/or have characters that do not sufficiently match diagnosed species.

### Popular Prehistoric Taxa Do Not Require and Must Not Receive Special Scientific Treatment

Statements that fossil taxa that enjoy exceptional levels of popularity deserve and require special levels of scientific analysis at the species level are entirely nonscientific and disturbing. Note that such has not been said regarding extant taxa. Widely liked *Loxodonta* has been split into two species (Grubb et al. 2000) with no attention paid to popular thinking on the matter, and the probability that extant *Giraffa* are multispecific (Coimbra et al. 2021) has not aroused ardent dispute in popular venues. The quality of scientific research should obviously be the same regardless of public opinion regarding the taxon.

Taxonomic implications – Popular feelings must not play any role in scientific procedures.

### The Scannella et al. Paleospecies Standard

The determination of multispecies of *Triceratops* by Scannella et al. (2014) has been considered as setting a new and high standard for the procedure among dinosaurs that other works should aspire to (as per Paul et al. [2022], Carr et al. [2022] also cite the study as a favorable example). The conclusions of the study are widely accepted and have not been challenged. A detailed examination of the *Triceratops* study shows that while its conclusions are sound, its data and analytical methods contents should not be overstated.

The stratigraphic data is more detailed than usual for Mesozoic dinosaurs, yet in the end it still comes down to basically three levels of the Lancian mid latitude Laramidia TT-zone (as per Paul et al. 2022), lower, middle, upper -- in their Figure 2 the intricate original data illustrated in Fig. 1 is condensed down to 6 levels, and further contracting them to 3 as they effectively do does not make a difference in the final results. The sample from the lower level is relatively small compared to those from higher in the column. The entire sample is from a limited geographic area, so a large portion of *Triceratops* remains are not considered (Fig. 7A-G,J-M for example are not in the study).

**Fig. 7.**
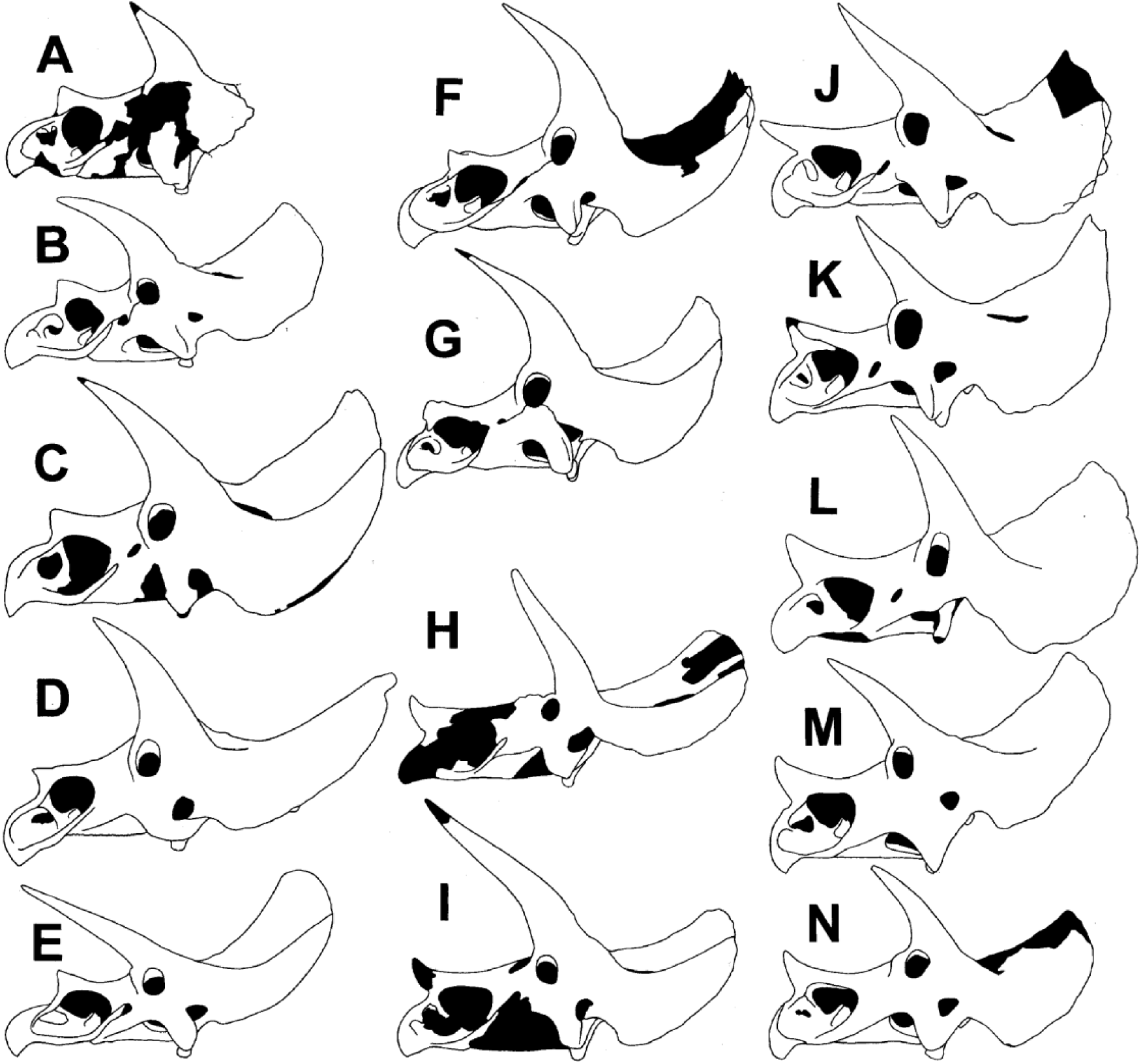
Same main length comparisons of *Triceratops* skulls. *T. horridus* from lower TT- zone: **A** holotype YPM 1820; **B** TCM 2001.93.1; **C** SDSM 2760; **D** MNHN 1912.20; **E** AMNH 5116; **F** USNM 4928; **G** USNM 1201. *T*. sp. from middle TT-zone: **H** UCMP 113697; **I** MOR 3027. *T. prorsus* from high TT-zone: **J** holotype YPM 1822; **K** YPM 1834; **L** SMNH P1163.4; **M** LACM 7207; **N** MOR 1604.

**Fig. 8.**
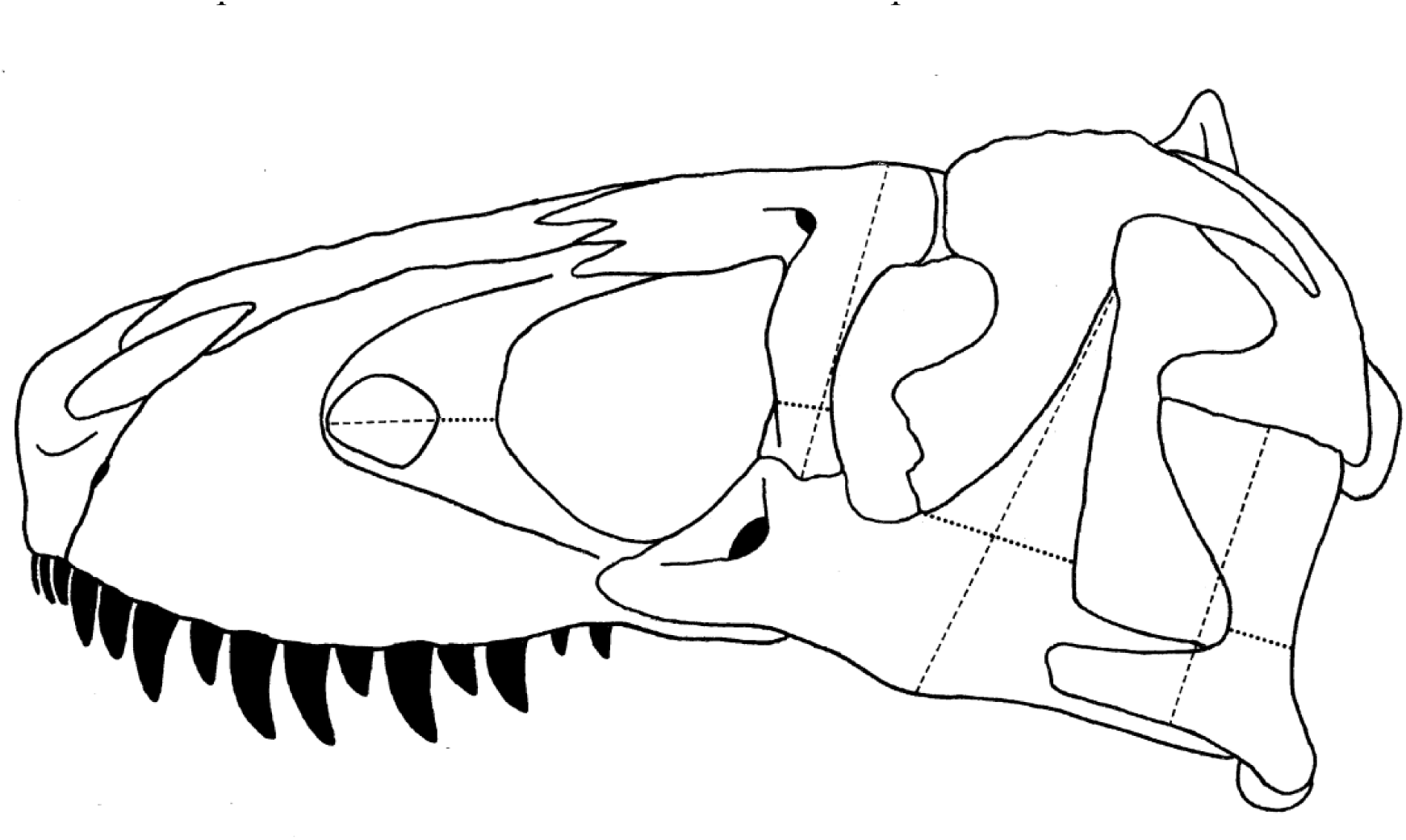
Generalized *Tyrannosaurus* skull showing measurements of skull dimensions presented in Table 1 and Fig. 9K-N.

Scannella et al. incorporate an unusually large sample for dinosaurs, about two and a half dozen specimens of large subadults and adults of varying completeness being stratigraphically correlated (Fig. 1 in Scannella et al. [2014]). All are skulls of which the mandible is not examined, nor are postcrania. In a number of cases dimensional values used to calculate ratios are estimates. 6 character ratios or angles relative to stratigraphic level are plotted, all cranial. A few other nondimensional characters are examined. Many hundreds of characters are not assessed. Of the 6 ratios and degrees it is 5 that exhibit significant trends with time, all having to do with the anatomy of the rostrum. Although there is not overlap in the ratios between the extremes of early *T. horridus* and much later *T. prorsus i*n 5 examples, there is always some degree of overlap one way or another with the unnamed intermediate species, in 5 cases considerable. There are some ratio outliers within species, particularly *T. horridus*. Ergo, clear character separation between the proposed species and bimodality is affirmed as not being the norm. This is all the more true because inclusion of specimens from outside the geographic study zone is certain to further increase the overlaps.

An example is the length of the brow horns relative to that of the main body of the skull. There are no particularly short or long examples in *T. prorsus*, so this is an attribute that can be used to help diagnose the species unless future discoveries indicate otherwise. *T.* sp. have postorbital horns that range from moderate to very long, so the later can be used to help diagnose the taxon at least relative to *T. prorsus*. Overlap with both other species is extensive. *T. horridus* brow horns can be almost as long as those of *T*. sp., the difference not being statistically significant. Within the sample utilized in Scannella et al. (2014) *T. horridus* lacks short postorbital horns. But those of the large *T. horridus* holotype (Fig. 7A) are shorter than any other *Triceratops* including those in the Scannella et al. (2014) sample, so brow horns cannot be used to diagnose that species. How much of the brow horn variation is due to dimorphic, individual or ontogenetic factors has not been fully explored. That the small nasal horn clearly distinguishes *T. horridus* from big nose horned *T. prorsus* makes this and especially clear cut species specific feature between the two taxa, but this factor is highly variable in the intermediate level skulls that extensively overlaps the other two species while not reaching the extremes of either, so *T*. sp. nasal horns at most can be used to define that taxon by not being either especially small nor large. While the size of the nasal horn is very different between *T. horridus* and *T. prorsus*, the orientation is not, that factor ranging from nearly horizontal to much more vertical in both due to unknown levels of dimorphism, individuality, or ontogeny. In Scannella et al. the one plotted rostrum of a *T. horridus* is longer than those of any other *Triceratops*, and such appears true of other members of the species relative to the higher placed species (Fig. 7B,D-F), but the beaks of other specimens of that taxon including the holotype do not appear to be especially long (Fig. 7A,C,G). All *T. horridus* specimens in Scannella et al. sport an acute angle between the nasal process and the narial strut of the premaxilla in contrast to the shallow angle common to *T. prorsus*, but in USNM 2412, 4928, and SDSM 2760 the angle is very shallow, perhaps more so than yet observed in *T. prorsus*. Future expansion of measurements of the narrowness of the nasal process of the premaxilla is likely to increase the degree of overlap between the species, indeed this high probability applies to all the characters.

While the combined anatomy of *T. prorsus* skulls is fairly consistent (Fig. 7J-N), that of *T. horridus* is highly variable (Fig. 7A-G; Fig 2 in Scannella et al. [2014]). all the more so when *N. hatcheri* and *T. latus* are considered to be in that species (Scannella et al. 2014; Paul 2016), so *T. horridus* is harder to define, and the possibility that multiple species are involved is a real possibility. The relative scarcity of *T.* sp. specimens, and that none are highly complete (Fig. 7H,I), hinders assessing and diagnosing that species.

No autapomorphies were noted by Scannella et al. (2014), collective differences being used to sort out the morphotypes as is the frequent practice (contra Carr et al. [2022] stating unique characters being a necessity for taxa). Interestingly Scannella et al. (2014) did not offer a formal systematics diagnoses of the species with assigned specimens even regarding those that they sampled. For purposes of comparative results draft diagnoses are done here, using their characters in the systematics section. Some specimens included are from outside their sample. The results show that the three species are not separated from one another by clear, nonbimodal boundaries with consistent character separations. There is considerable overlapping and some ambiguity, with the occasional character exceptions. This is as explained in Paul et al. (2022) and herein normal in biology, being the result of mosaic evolution – the potential of hybridization is low for this genus because the species are sequential rather than contemporary. The large expansion of the sample will invariably further blur the boundaries between the species. Ergo, the diagnoses are inherently somewhat unstable, and will always be so because of the constant expansion of the fossil sample over time. Because the three morphotypes do not overlap in time sexual dimorphism cannot be used to explain the observed pattern. Nor do individual variation or ontogeny offer explanation for what is an apparent selection driven evolutionary pattern. All lower TT-zone *Triceratops* are currently assignable to *T. horridus,* and all high placed fossils are *T. prorsus*, in part because there is not solid current evidence for contemporary species at those levels the higher especially, in part because specimens appear to sufficiently fit into one or the other taxon. But the much greater variability present in *T. horridus* leaves open the possibility of more than one species in the lower TT-zone. Also possible and more probable is sexual dimorphism, with *T. latus* perhaps being the adult males of *T. horridus* (Paul 2016) – the consistency of *T. prorsus* skulls interestingly leaves no compelling evidence of dimorphism in the species. The situation in the middle level is more ambiguous regarding the number of species and assignments of specimens. Further work may be required to assign some specimens, and it may not be possible to place all examples in the future.

It is notable that the basal *T. horridus* retained some basic characteristics of the earlier relative *Eutriceratops xerinsularis* (Scannella et al. 2014, Paul 2016) of a small nasal horn, a long anterior rostrum, and the often large brow horns (indeed the strong similarity between *E. xerinsularis* and *T. horridus* suggests they are congeneric, as per Paul [2016], all the more so because *T. xerinsular*is is in many regards more similar to *T. horridus* than the latter is to *T. prorsus*). The later derivation of *Triceratops* away from the old morphotype over the span of the hundreds of thousands of years of the TT-zone is additional and strong evidence that it was undergoing selection driven speciation. Scannella et al. could not quantitatively compare the degree of variation in *Triceratops* to still earlier triceratopsines, the data sample for the latter does not exist.

While it incorporates a half of a percent of the characters scored in Carr (2020), Scannella et al. (2014) is a far superior examination of intragenera species because the latter was designed to test the question rather than assuming monospecifity, and stratigraphically correlated over five times as many specimens.

Taxonomic implications – Having helped set the modern standard for assessing dinosaur species within a genus, Scannella et al. (2014) establishes the following. Necessary for best results is a substantial sample size in the dozens but not necessarily many dozens of specimens – if that is considered insufficient then the conclusion that *Triceratops* or any other dinosaur genus encompasses more than one species is to date not substantiated because larger samples of measureable specimens are either not available or have not yet been analyzed in terms of species determination, a point that applies to an enormous number of tetrapods extinct and extant. All or nearly so of the specimens need to be stratigraphically correlated if a large geological time span is involved. The latter ultimately can be assessed at gross levels of lower, middle and upper within a formation. The sample does not necessarily need to incorporate all known major specimens although doing so produces the most complete possible results [contrary to the insistence that all significant remains be placeable into paleospecies by Carr et al. [2022)), and can be limited to just cranial characters. Species diagnoses are not set in stone, being subject to significant alterations as new specimens and analysis comes on line. Quantitative dimensional values need not be precise when such are not fully preserved for exact measurement. Unnecessary are large numbers of characters that show strong trends with time, nor clear, consistent, nonoverlapping and explicit character separation and bimodality between species, autapomorphies, or the ready ability to assign all well preserved skulls to a species [contra such claims in Carr et al. 2022). If the basal species retain ancestral conditions that reinforces the reality of that species relative to latter, more derived taxa. The exercise of formally diagnosing the *Triceratops* species demonstrated the utility of the practice, and indicates it should be the required, systematic norm when assaying and determining paleospecies (this is in line with Paul et al. [2022] and Carr et al. [2022].

### Paleospecies Non/Necessities – Results and Summary

Monospecificity is not the automatic null hypothesis relative to multispecificity within genera, if anything the opposite is true. A given situation is resolved by the preponderance of the currently available collective evidence.

The longer the time over which a genus exists, the more probable it will be the fossils it contains include multiple species, specifically if the existence span exceeds a few hundred thousand years. In that case the null hypothesis shifts somewhat in favor of multiple species.

Between two and three dozen adequate specimens have been used for modern statistical sibling paleospecies work. Far less at the upper end have been used to designate sibling paleospecies, which not do automatically require statistical analysis to be widely accepted.

A study reexamining a prior establishment of paleospecies should not utilize a smaller sample of specimens to do so. All diagnostic type specimens need to be utilized.

Sibling paleospecies are normally diagnosed by a small number of skeletal attributes, as few as one. This is true even if the specimen sample is large.

Degrees of variation observed in one genera need to be compared to those present in other genera, and on up the systematic ladder if necessary, within the clade to help determine the species norms.

Characters used to diagnose sibling paleospecies do not need to include sexual display structures, especially in predators, but because visual species IDs are part and parcel of what a species is, such visually obvious differentiation display features alone are sufficient to determine, diagnose, and name sibling species even when the crania and postcrania are otherwise identical.

Character distribution between sibling species often is not nonoverlapping, bimodal or consistent.

Autapomorphies need not be present to define sibling species. Reptilian teeth can and are used to help diagnose paleospecies. Element robustness can and is used to help diagnose paleospecies.

Because the available data is often improving over time, the diagnoses of paleospecies are often adjusted over time.

When the time span covered by a genus is sufficient for speciation to be a serious possibility or probability, over a few hundred thousand years, it is critical that as many specimens as possible be stratigraphically correlated, in order to try to discern if patterns that indicate the evolutionary trends indicative of speciation exist or not.

Stratigraphic correlations to not need to be very tightly constrained, basic time separations are sufficient to geotemporally sort out paleospecies.

To produce compelling results a follow up reexamination of prior works on a set of intragenera paleospecies intended to test the earlier results, the reexamination needs to be a thorough analysis that addresses all critical aspects of the preceding studies.

Not applying the same high scientific standards to unpopular extinct taxa is unscientific, and popular fossils do not require and should not receive elevated standards.

If a substantial sample of fossils from a genus that existed over a sufficient period of time for speciation to have occurred that also exhibit notable variation between specimens is on hand, then defending and establishing monospecifity is not achieved by preferential opinion. A single species must be shown to provide a more coherent and cogent explanation for the variation in the context of the logic of evolutionary adaptation than the multispecies alternative. This is all the truer when the variation exhibits reasonably consistent shifts over geotime.

The methods and procedures for determining paleospecies are not highly rigorous and exacting, biology being inherently irregular and sloppy, not precise and consistent.

All analyses of paleospecies do need to incorporate and consider the alternative possible explanations for the observed pattern, including dimorphism, individual variation, and ontogeny, and a systematic species diagnosis.

### Sorting and Diagnosing *Tyrannosaurus* Species

#### Prior Work Did Not Show That There is Just the One Species

From the naming of *Tyrannosaurus* (Osborn 1905) until the 1980s the paucity of specimens precluded analysis of the species the contains. Paul et al. (2022) noted that even as the sample began to grow, no prior study had investigated the question of *Tyrannosaurus* species in close to sufficient depth. Carpenter (1990), Larson, (1994), Carpenter and Smith (2001) and Brochu (2003) were working with the very limited sample of specimens available at those times, and did not directly test the species question it being de facto assumed there was one species. Larson (P. 2008) was the one analysis that had taken a serious look at the topic, and is the first to examine the varying robustness versus gracility of a number of elements in a substantial sample. But that sample was still inadequate. A stratigraphic correlation – which at the time would have observed the critical absence of graciles and one incisiform dentary toothed specimens low in the TT-zone – was not conducted. The degree of femoral variation in *Tyrannosaurus* was not compared to other species and groups. Statistical analyses were not executed.

In the wake of the Paul et al. (2020) study, Carr (2020) which utilized 1850 characters was suddenly, insistently and remarkably proclaimed as a definitive prerebuttal of the later work that Paul et al. had errantly failed to take into necessary account, including Carr et al. (2022). The titles and contents of the Carr (2020) refute the claims. Carr (2020) is titled “A high-resolution growth series of *Tyrannosaurus rex* obtained from multiple lines of evidence,” there is no mention of the species issue. Or in the introductory sections of the paper. Nor in the conclusions except for a brief mention that the *Tyrannosaurus* “x” hypothesis is not viable which Paul et al. (2020) agreed with (but see below). The titles for the supplements include “Character list used to resolve the ontogeny of *Tyrannosaurus rex*, sources cited, and list of ordered characters,” and “Character states for each specimen included in the character matrix for recovering the growth series of *Tyrannosaurus rex*,” none of the additional materials were claimed to be pertinent to the species problem. All that makes sense in that the paper focused on the status of the small tyrannosaurid specimens from the TT-zone vis-à-vis the adults, and barely addressed the systematic status of the large specimens that Paul et al. focuses on.

The first mention of intra *Tyrannosaurus* species in Carr (2020) was in a section titled “Assumptions” in which the following is stated -- “For the purposes of this study, it was *assumed* [italics added] that the assemblage of *T. rex*, which spans Laramidia for a duration of less than 1.0 million years (Fowler, 2017 -- note: was more likely 1.5 million or even more (Mallon et al. 2022; Paul et al. 2022 and refs. therein) was a single nonanagenetic population.” Carr basically presumed that there was only *T. rex* from the start, so the paper did not conduct a serious test of the issue in a study mainly looking at the proposed growth of small specimens into the one monospecies. Therefore, because Carr did not take an in-depth look at the species question, that prevented him from finding the evidence for more than one species, so he assumed one species. In contrast, Paul et al. (2022) did not make a-priori assumptions about the number of species in the genus of interest, it being part of a long term exploratory effort to see what would turn up one way or the other, and based any whatever conclusions arose out of the preponderance of evidence.

There was a small effort to look at the stratigraphic issue in Carr (2020), but it included only 7 adult specimens (Table 18), just a quarter of the number geologically correlated in Paul et al. (2022, Table 1 and Fig 6 therein). That the stratigraphy was limited to simple lower, middle and upper was entirely acceptable, but even had the paper been designed to detect speciation patterns, so few correlations were far from sufficient to begin to properly test the competing species hypotheses. Carr did not examine the robustness, as is critical to the later Paul et al. (2022) study, of the maxilla, dentary, ilium, and humerus. This is difficult to understand because these parameters had been examined in Larson (P. 2008) some of whose other characters were reexamined in Carr (2020), and have obvious potentially critical importance in multiple regards. Without this data species analysis is simply not practical. Femur stoutness was considered for only 4 large specimens, of which just 2 were geoplaced (Paul et al. [2022] has a dozen times more stratigraphically placed femurs) which is statistically useless. Same for just two tibias (problematic to use because the strength of the parallel fibula that bore part of the stress load of the middle limb is not taken into account), and there are no adult metatarsals. Nor did Carr look at the fine gradation of robustness as per Paul et al. (2022), it was only scored whether the femur ratio is above or below 2.27 (a value that is too low because BHI specimens are excluded, as result of the inclusion of those specimens the more correct dividing value is 2.4 in Paul et al. [2022]). What data is in Carr (2020) did not find is low lying graciles. In Carr (2020) the anterior dentary teeth were processed in a nonquantitative manner that is statistically inferior, and with a smaller sample both in total numbers and those that can be stratigraphically assessed than in Paul et al. (2022). While the Carr stratigraphic sample is much too small (in part because it excludes all private specimens) to be definitive, it does show all graciles were high in the TT-zone. Not considered in Carr (2020) was the amount of variation in *Tyrannosaurus* compared to other dinosaurs, tyrannosaurids especially. That was entirely logical because Carr’s paper was not devised to examine the species question.

Repeatedly emphasized in Carr et al. (2022), as well as earlier commentaries. were the “1850 variable characters from throughout the skull and skeleton for over 40 [44] specimens” supposedly contained in Carr (2020). The implication, probably unintended, was that the characters were scored 81.400 times. Of the 44 specimens 26 were large specimens critical to species determination. Nor were 81,400 characters actually assayed that not being practical. Hundreds were examined in only two, three or a few specimens. A very large portion were recorded in only one juvenile and one adult, with many being minor attributes of vertebrae caudals especially, and manal and pedal elements, apparently in order to produce an accounting of basic differences between a small individual and an adult in tune with the actual primary subject of the paper. Only a modest fraction of the characters were scored for up to a dozen to a dozen and a half large specimens – and not many more small TT-zone tyrannosaurid fossils they being scarce. So while the Carr character list was laterally broad, its sample depth was too shallow to be statistically highly useful. In a large number of cases it is not entirely clear exactly what was being assessed, it not being feasible to clearly illustrate the item being examined and how. An example is character 601 which concerns a groove on the postorbital of ambiguous nature. As a result, verification and replication are often difficult at best and may not be practical – there is irony in this in that Carr has strongly criticized the problems of data replication of privately held specimens as further discussed below. Some features were assessed in a simplistic manner. For example, the postorbital bosses were described as prominent in all large individuals even though there is considerable variation in their development as detailed below, the irregularity of the nasal ridge was treated in a similarly either or manner; presumably again because the only purpose was to contrast the juvenile and adult conditions. A quite large number of the characters that were observed in a substantial number of specimens appear minor in nature and their taxonomic value is correspondingly problematic, risking being the potentially misleading background noise issue cautioned about above.

It is notable that while Carr et al. (2022) promotes the value of the 1850 character tabulation in Carr (2020), the former does not attempt to use the latter to examine the species question.

While it incorporates over a third of a percent of the characters scored in Carr (2020), Paul et al. (2022) is a far superior examination of intragenera species because the latter was designed to test the question rather than presuming monospecifity by zeroing in on those few characters that have species identification potential, and geologically correlating over five times as many specimens.

The Carr (2022) work abjectly lacked the ability to discern a species level taxonomic signal with just 7 (albeit adequately) stratigraphically correlated specimens, only 2 femora tied to the geology, and robustness measured for just a few elements of a few specimens, in a work that largely assumed one species from the start. That 1850 characters were examined was not decisive because they are off a limited set of specimens too limited in number to be statistically assessed, and they are missing key measures of robustness. Carr (2020) paper did not scientifically test the number of species recorded by large *Tyrannosaurus* remains which it was not designed to do. Paul et al. (2022) correspondingly did not even think to utilize the Carr data set because that was neither set up for the purpose that Paul and company were investigating. Had such been attempted it would have been a waste of effort because the data set is not up to the task at hand. E. g. of the large specimens in Carr (2020) just 11 can be assessed using the Paul et al (2022) stratigraphic data, compared to the much more quantitatively significant 31 in the later study. The only data specific citation on Paul et al. of Carr (2020) is a note that what stratigraphic data in the latter is present is in line with that in the former, and the femoral ratios for the same specimens are also in good accord. The claim of the Carr (2020) title that the study is high resolution is exaggerated to the point of being perplexing, Paul et al. (2022) is more refined and sophisticated in critical aspects, especially regarding assaying paleospecies. Rather than having demonstrated *Tyrannosaurus* monospecificity, the 2020 study confirms that (as observed by Paul et al. 2022) little effort had been conducted to directly challenge the *T. rex* issue due to the long casual assumption there was only one species. That leaves Paul et al. (2022) as the first and until this work the only one to directly take on the issue seriously with a sufficient data set.

Taxonomic implications – Claims in the media and then Carr et al. (2022) that Carr’s 2020 analysis was a resounding prerefutation of the Paul et al. (2022) paper that had not yet been published, and that contains a many times larger sample of stratigraphically placed specimens whose robustness is much more extensively examined, as well as cross comparisons of variation in *Tyrannosaurus* relative to other theropods, should not have occurred, and it cannot be scientifically cited as such in the future. Carr (2020) barely addressed the subject and lacked the data to do so. No work prior to Paul et al. (2022) has significant impact on the species issue – so *T. rex* increasingly was a taxonomic wastebasket as specimens accumulated without rigorous testing of the contents of the genus -- further testing the subject will require ongoing and future work. As for future application of the Carr data set on *Tyrannosaurus*, which Carr et al. (2022) did not attempt, that may not prove as productive as hoped, for the reasons discussed earlier in this analysis. That the nearly all the 1850 characters may prove randomly distributed is not of importance that being normal between species, it is the few differences that count. Until the efficacy of the mass character set method is properly tested with a number of other suitable genera, its utility regarding *Tyrannosaurus* is open to challenge – if Carr in any future work claims to demonstrate one species based on his 2020 data base, then how will that be verified if the same procedure has not been used to affirm or deny the species in *Triceratops*, and *Panthera*?

### Much More Than Just Two Features Separate the Three Species

In order to try to preclude claims that just two characters distinguish the species, as Carr et al. (2022) and prior commentaries did anyway, Paul et al. (2022) explicitly states immediately before the systematic diagnoses that “Note that the species diagnoses incorporate the cumulative proportions of six elements in addition to the femur.” The diagnoses specifically state that is expressed as a matter of general robustness or gracility, which includes the maxilla, dentary, humerus, ilium, femur, and two metatarsals. Only the dentary does not show a plain trend towards gracility with later time, although there is no example of any low geoplaced strongly gracile element. The three holotypes possess all or nearly all of the 7 pertinent elements. Also observe that all those elements show a clear pattern of little variation low in the TT-zone to more variable higher up. This was documented in the data tables and visually in Figure 6A-I. Also note that metatarsals as well as femora were illustrated in Figure 2. The paper uses only one specific robusticity ratio for defining the species, 2.4 for the femur, because that is the only practical way to produce a value that can be readily applied, the individuals all having some internal variation in robustness, and the massive proximal hindlimb element being most commonly preserved intact. In addition to the robustness of important elements, the condition of the anteriormost dentary teeth were utilized, so the total number of elements and character examined was 8. The inaccurate claim that the study works with just two characters should not have been stated, and must never be repeated.

In part to make more clear the number of characters being used to characterize *Tyrannosaurus* species, the systematic diagnoses are more explicitly stated in the systematics section.

Taxonomic implications – The number of characters utilized by Paul et al. (2022) was 8, with 7 showing a cumulatively significant shift from one condition to another proceeding stratigraphically through time, and all 7 robusticity measures further cumulatively showing a very significant increase in variation.

### More Quantitative Cranial Data

Carr et al. (2022) continued to protest the inability of Paul et al. (2022) to place at the species level four largely or entirely complete skulls, although they conceded that the current display status of AMNH 5027 hinders its assessment.

Paul et al. (2022) found that two major nondental cranial features, the length/depth ratios of the largest and primary tooth bearing skull bones the maxilla and dentary, are compatible with an increase towards gracility and/or variation with time in the genus. Four vertical bars the width of which can be measured provide further data on the dorso-ventral strength of 16 *Tyrannosaurus* skulls with which to further test the species question -- the interfenestral pillar of the maxilla, the lacrimal, the postorbital process of the jugal, and the posterior ramus of the quadratojugal. The methods in which they are measured is illustrated in Figure 8, and plotted in Fig. 9K-N (values listed in Table 1). These bars help resist the intense biting force of the giant predator’s massive jaws (Gignac and Erickson 2017). It is often difficult to accurately measure or restore the actual dimensions of skulls because of distortion, and because many are reconstructions assembled from multiple disarticulated elements of varying completeness, so the skull lengths listed in Table 1 are sometimes approximations, and using them for direct statistical comparisons to the dimensions of elements is problematic.

**Fig. 9.**
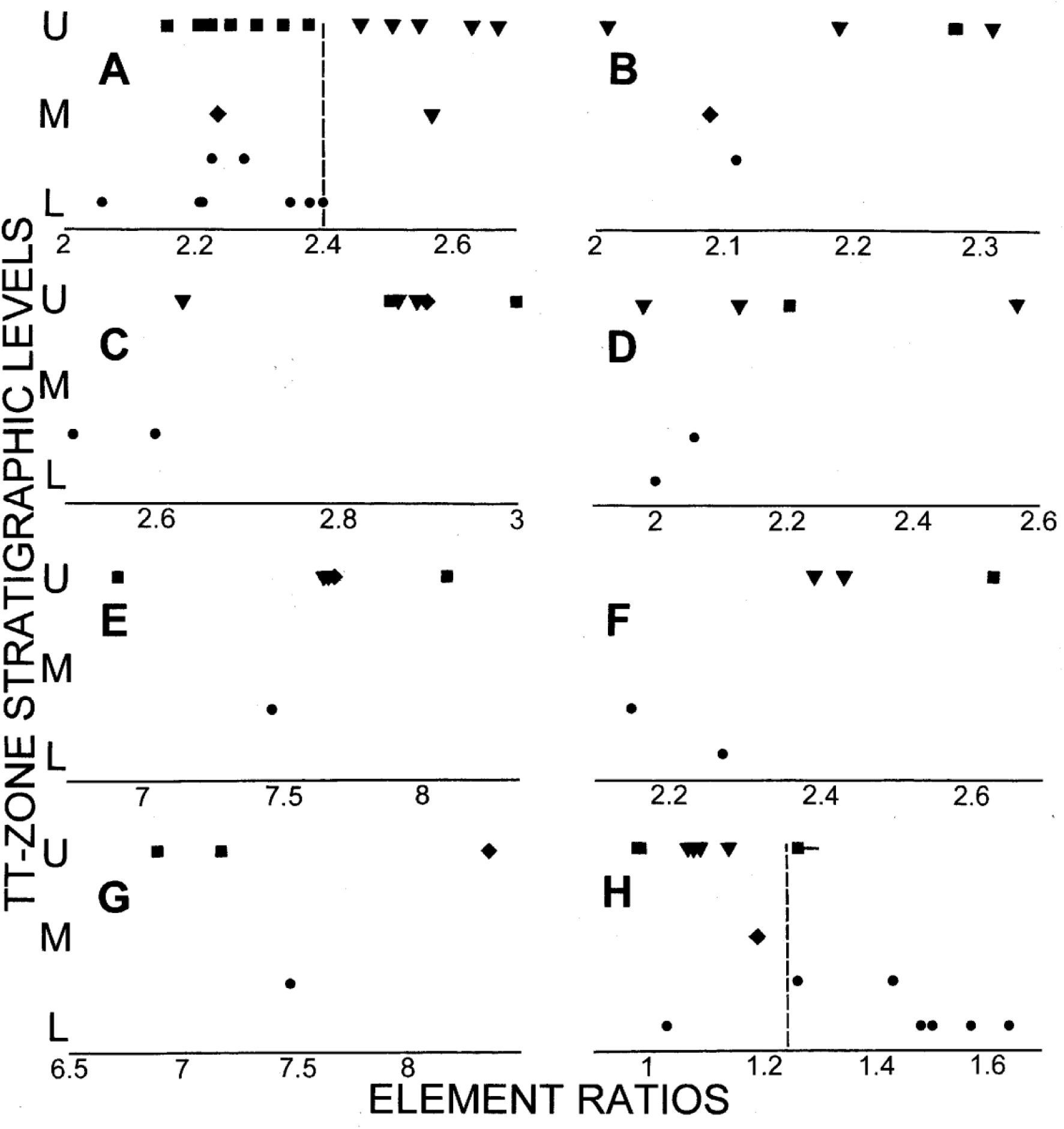
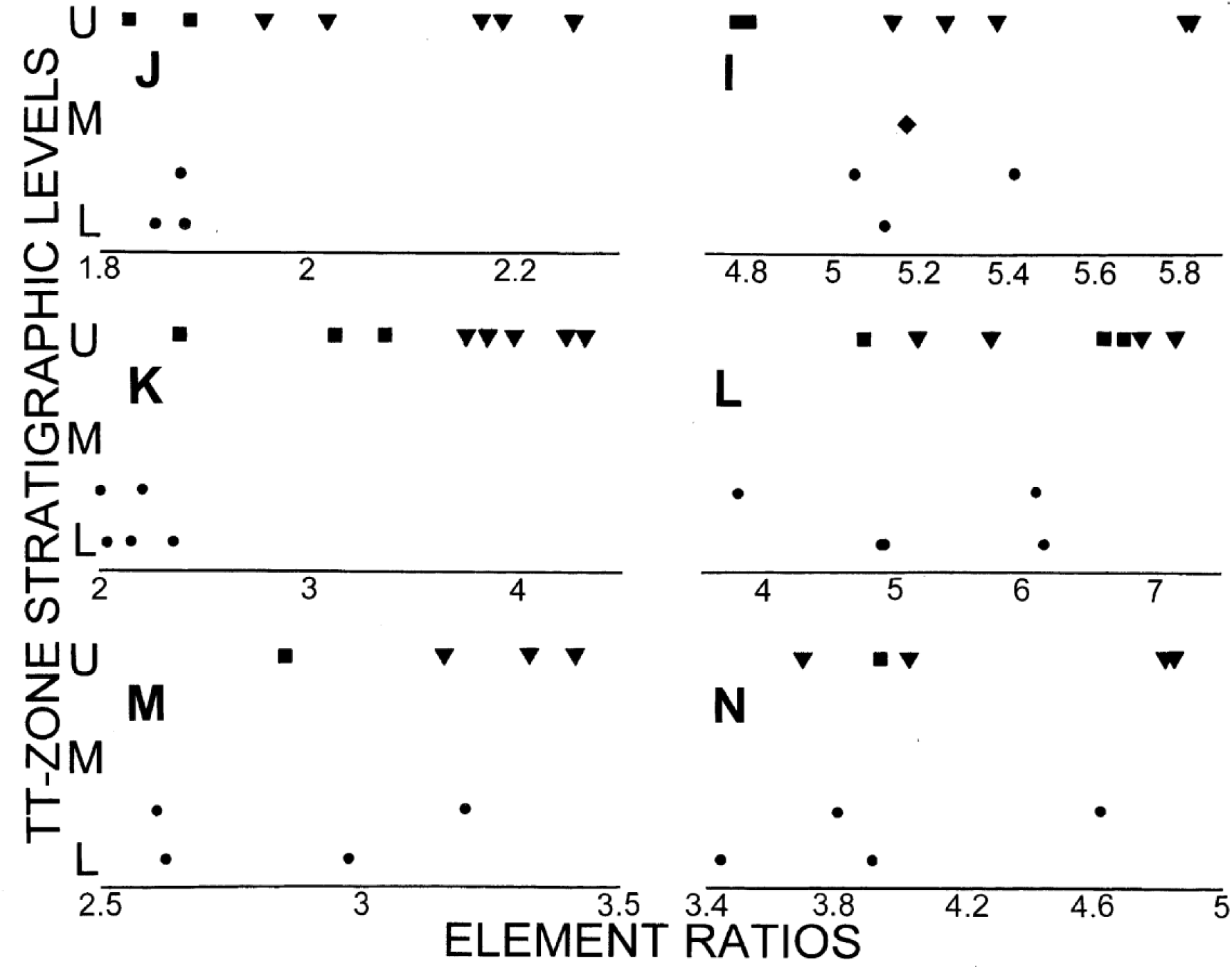
Element ratios for large *Tyrannosaurus rex* (squares), *T. regina* (inverted triangles) *T. imperator* (circles) and *T.* incertae sedis (diamonds) specimen at differing stratigraphic levels (lower L, middle M, upper U) in the TT-zone; specimens that may be from either the upper lower or lower middle T-zone are plotted between the lower and middle levels. For A–G, I-N increasingly bone gracility is to the right, for H increasing 2nd incisor robustness is the left. **A** Femur length/minimum circumference, divisions at 2.4 between robusts and graciles indicated by vertical dashed line. **B** Humerus length/min. circ. **C** ilium length/depth. **D** Metatarsal 2 length/min. circ. **E** Metatarsal 2 length/min. diameter. **F** Metatarsal 4 length/min. circ. **G** Metatarsal 4 length/min. dia.. **H** Dentary teeth/alveoli 2/3 (possibly 3/4 if 1 is no longer functional) anteroposterior base diameters, division at 1.25 between one and two incisors indicated by vertical dashed line, (horizontal line indicate different value of Carr et al. 2022). **I** Dentary length/depth. **J** Maxilla length/depth **K** Maxillary fenestra length/interfenestral pillar min. width. **L** Lacrimal height/min. width. **M** Jugal height/postorbital process width. **N** Quadratojugal height/min. width.

The hourglass shaped interfenestral pillar separates the small maxillary fenestra at the anterior end of the antorbital fossa from the much larger antorbital fenestra that fills most of the fossa. The width of the bar at its anteroposterior narrowest is visually highly and distinctively variable in *Tyrannosaurus*; the width of the bar at its base tends to correlated with that at its midpoint. The stoutness of the pillar is most readily assessed by comparing its width to that of the antorbital fossa it helps contains -- the combined size of this complex is fairly consistent relative to the rest of the skull. The pillar is relatively broad in the lower TT-one *T. imperator*. All of those specimens cluster tightly together -- the value for the holotype is an approximation due to damage to the skull, that it is broad is indicated by its wide base. All pillars in skulls assignable to *T. regina* have slender interfenestral pillars, much more so than yet observed in any *T. imperator*. Those of *T. rex* are intermediate in breadth, all being more robust than *T. regina*, and slenderer than any *T. imperator*, so there is no observed overlap in the sample between the three species, which equals two cases of bimodality in the current sample. The variation in robustness with more recent time is, as often observed in *Tyrannosaurus* elements, substantial, very much so being about five times as great in high placed skulls than it is in the earlier examples. As a result of the above the shift from robustness in the low TT-zone to much more gracility later in time is very strong. The robustness versus gracility of the entire maxilla broadly parallels that of the pillar it contains in that those of *T. regina* always being shallower than those of *T. rex*, as well as *T. imperator*. In both these comparisons of strength the maxillae of *Tyrannosaurus* provide some of the best evidence for three over two or one species (Fig. 9J,K).

The preorbital bar formed by the arced, narrow lacrimal is more difficult to assess in part because it has a complex twisted shape along its long vertical axis, and its minimal anteroposterior diameter must be carefully assessed to not under measure the minimum width. With that caveat, the thickest lacrimal bar is present in *T, imperator*, and thinnest bars belong to *T. regina* always have a thin bar, so while there is extensive overlap there is a trend towards more gracility with stratigraphic height, but not towards more variation – although a decrease is not observed (Fig. 9L).

The strength of the postorbital bar is best measured by the breath of the sharply triangular, plate like ascending process of the jugal at the level of its ventral most articulation with the postorbital, that being compared to the total height of the jugal to generate the ratio – note that the width can look narrower than it is in direct lateral view images of skulls because the lateral surface of the jugal is directed somewhat anteriorly in the genus because the narrow snout flares laterally to the much broader temporal region at this location. Again the most robust examples are early *T. imperators*, the most gracile are late *T. regina*s (Fig. 9M). As with the maxillary pillar, the *T. reg*ina bars are always less robust than those of *T. rex*, although the sample of the latter is one. There is extensive overlap with *T. imperator*, even so an overall trend towards more gracility with time exist, yet no trend towards more variation. Most of the same pattern applies to the strongly embayed quadratojugal, which likewise does not decrease in variation with later time (Fig. 9N). Interestingly, TMP 81.6.1 shows the greatest breadth among the *T. regina* in its aft two bars, while having slender anterior bars, showing a mosaic pattern in the taxon.

A visual survey indicates that the interfenestral pillar of the maxilla is normally robust in earlier large North American and Asian tyrannosaurids (Maleev 1955; Russell 1970; Currie 2003a; Hurum & Sabath 2003; Carr & Williamson 2010; Carr et al. 2011, 2017; Brusatte et al. 2012; Lu et al. 2014), so thick maxillary bars in basal *Tyrannosaurus* is another example of the retention of the ancestral condition in early *T. imperator*, along with its robustness the femur especially, and two incisiform dentary teeth (Paul et al. 2022). And the exceptional gracility of the maxillary pillar and femur of *T. regina* are very atypical adaptations that appear late in the family and genus. It is notable that the basic pattern with all the vertical skull bars is in tune with the overall changes in robustness during the evolution of and within *Tyrannosauru*s, with the advent of exceptional gracility seeing the skull becoming more lightly constructed. This is most clearly seen as the maxilla overall becomes shallower, the interfenestral bar becomes more delicate, but the same applies to a fair extent in the rest of the supporting bars. This reinforces the evidence that a population of late *Tyrannosaurus* were reducing the strength factors of the head and body in the genus characterized by its overall massiveness, and this shift included the snout in an animal know for the power of its bite that apparently exceeded that seen in any other known land predator, a major and consistent trend that is not readily explained by individual variation, dimorphism, or ontogeny.

The observed cranial strength patterns fit that predicted by three rather than two or one species. All known low TT-zone robust *T. imperator* sport interfenestral bars broader than those of latter gracile *T. regina*, and all contemporary robust *T, rex* have bars thicker than those of *T. regina*, with even the fairly stout *T. rex* interfenestral pillars not overlapping with those of the earlier robust *T, imperator*. The situation is not as clear cut with the other measures including the dentary, but in all cases it is *T. regina* that is most lightly constructed, in all but one it is *T. imperator* is the strongest, and in the maxilla *T. regina* is always deeper than that of *T. rex*.

With the expansion of the cranial data set the observed patterns are now sufficient to place all but one of the *Tyrannosaurus* skulls -- none of which are prefect as has been asserted by some -- and skeletons, a few of which had been taxonomic floaters, in one of the three named species at least tentatively. Like the ratio for its pillar, the length/depth ratio of the maxilla of RGM 792.000 of 1.85 is in the robust zone, as is its femur, reinforcing its placement in low TT-zone *T. imperator* (Fig. 9A,J,K; Table 1). A similar maxillary ratio for UWBM 99000 indicates that the high TT-zone specimen is a *T. rex*, as does the broad interfenestral pillar it contains. The thick pillar of the high placed UCMP 118742 maxilla favors placement in *T. rex*. LACM 23844 is borderline regarding its maxilla, dentary and lacrimal. and looks robust in a metatarsal, but the slender interfenestral pillar implies it is a *T. regina*. Species identification of the prior is aided by their known stratigraphic placements. The last item is not yet true of MOR 008 the skull of which is not complete, the proportions of the maxilla are borderline, and the important limb elements are not on hand. The robust interfenestral bar and dentary, and two incisiform teeth are compatible with or suggestive of *T. imperator* status.

The new characters have been added to the revised and expanded diagnoses for the three species in the systematics section.

Taxonomic implications – Repeated variation in the strength of primary vertical strength bars in *Tyrannosaurus* crania records yet more and surprising strength reduction as the genus evolved. Of the four bars the evidence provided by the maxilla’s interfenstral pillar shows the strongest and most clear-cut trends, and appears to move away from the robust ancestral tyrannosaurid condition. The patterns observed in the pillar and the other three bars favors multiple species over one, and three over two taxa.

The evidence provided by the vertical bars is in tune with the overall skew towards slenderer and derived proportions observed in a number of other parts of the skull and skeleton in late *Tyrannosaurus, T. regina* especially, way from the ancestral condition, with *T. rex* retaining the most of the older anatomy. All three species gain new specimen members. The ability to place all but one *Tyrannosaurus s*kull in one of the three species removes the – spurious in its theoretical basis – objection to the multispecies hypothesis by Carr et al. (2022).

### The Quantitative Speciation Pattern

With the addition of the above new four skull characters, the number of ratio measured cranial and postcranial characters used to help track *Tyrannosaurus* speciation and diagnose the species is now a dozen incorporating 11 elements, although the two metatarsals are part of the united tarsometatarsus complex. The results include the following.

In all 6 cranial robusticity plots the most gracile ratio is that of a *T. regina* (Fig 9I-N). None of the *Triceratops s*pecies shows such consistency of proportional cranial extremity.

In only 2 plots (Fig. 9B,D) is a *T. regina* the most robust overall, in two cases the sample is on the small side, especially for *T. imperator*. *T. imperator* is never the most gracile in any element or ratio (Fig. 9A-N). None of the *Triceratops s*pecies shows such consistency of proportional cranial and postcranial extremity.

In 7.5 items the most robust or two incisor tooth condition is seen in *T. imperator* (Fig. 9A,C,F,J,K-N) -- the 0.5 applies to one of the unavoidably divided (see Paul et al. 2022) metatarsal measurements.

In 9 ratios the most gracile condition is observed in derived *T. regina* (Fig. 9A,B,D,J-L).

In 5 elements all *T. regina* are more gracile than any *T. rex* among high level specimens, exhibiting nonoverlapping bimodality between the two taxa despite the possibility of hybridization (Fig. 9A,J-K,M).

In 5 elements all *T. regina* are more gracile than any *T. imperator,* exhibiting nonoverlapping bimodality between the two taxa (Fig. 9A,CE,F,J,K).

In no element is *T. regina* always more robust than is its contemporary taxon.

As a result of the above, in all 11 elements and 13 ratios (Fig. 9A-E,J-N) there is an overall trend, form minor to strong, toward greater gracility progressing geologically upwards. Trends towards increasing robusticity have not been discovered.

There are 10 cases of nonoverlapping bimodal separation between the species (Fig 9A,C,E,F,J,K,M).

11 measurements of robustness in crania and postcrania favor actuality of robust and gracile morphs in generally good accord with femoral robustness.

In 7 characters there is in increase in variation within a given element from modest to many fold (Fig. 9A-E,J,K). In none is there a decrease in variation.

The proportions of at least the femur, the maxillary interfenestral bar, and the anterior dentary teeth shift significantly away from the ancestral tyrannosaurid condition of robust proportions or two incisiform teeth.

### Cranial Sexual Display Characters Support Three Species, Each Dimorphic – And Give *T. rex* a Distinctive New Look

Tyrannosaurid cranial displays were modestly developed, consisting of a long, low, irregular central ridge at the midline confluence of the paired nasals, short ridges of varying shapes on the top bars of the lacrimals, and variable bosses on the dorsal postorbital (Paul 2016).

15 of 16 large *Tyrannosaurus* skulls bear preserved postorbital bosses, with 7 now assigned to *T. imperator* including the holotype, 5 to *T. regina* including the holotype, and 2 to *T. rex* sans the holotype (Figs. 1 B-H; 10A-O). In these specimens there is a very large divergence in the prominence and form of the postorbital bosses, to a degree not seen within and even between some other tyrannosaurid genera (Fig. 6). This conspicuous lack of species specific consistency in the genus is a contributor to the skulls of the tyrant saurian looking unusually variable compared to other tyrannosaurid species (Paul et al. 2022 Suppl.), a point that has never been explained in the context of all the skulls being those of only the tyrant lizard king without royal siblings.

There has yet to be a systematic effort to assess and compare the form and development of the bosses in any effort to tease out their systematic implications. That is because within the simplistic hypothesis of one species the pattern appears chaotic, and therefore due to random, inexplicable individual variation, combined with ontogeny and perhaps dimorphism – Carr (2020) does not describe or score the postorbital bosses in the needed detail, and Carr et al. (2022) make no effort the examine the conspicuous ornaments. Also, the timing of this effort is fortuitous. It was only fairly recently that two new skulls with atypical postorbital bosses became available. So attempts to systematically sort out the taxonomic implications of the bosses coincidently became feasible only during the period when this researcher was working on the problem of *Tyrannosaurus* species starting in 2010, and was made possible by the advent of the data and results in Paul et al. (2022) and immediately above, which allowed the assignment of most skulls to species. As a result, the assignment of 14 of the skulls to stratigraphic levels combined with the above work placing 15 crania in the species serendipitously led to the revelation of a pattern that, in this first detailed examination of the supraorbital bosses of the genus, demonstrates species level differentiation between the taxa. And suggests that the species were probably internally dimorphic. The possession of such display organs is in line with the evidence for intraspecific combat recorded on the skulls of the genus (Brown et al. 2022).

It is presumed that if there is significant variation in the size and form of *Tyrannosaurus* cranial displays in a given species, that the bigger and more ornate expressions are those of males as per the null hypothesis – it being improbable that males exhibited parental care as do smaller crested male cassowaries (Paul 2008).

Unlike in most other tyrannosaurids, the *Tyrannosaurus* lacrimal is consistently little adorned and correspondingly so consistent in form that little if any taxonomic information can be obtained. The size of the lacrimal foramen is often difficult to measure and compare – it is drawn too small for AMNH 5027 in Figure 8.6 in Larson P. (2008) – and does not appear to show a consistent pattern in the genus (Persons pers. comm.).

The varying development of the postorbital boss was approximated by laying out photographs of all sufficiently well preserved left and right examples, which tend to be reasonably consistent in their configuration between the two sides of given individuals (Fig. 10A,B,F-K; this being least true of TMP 81.6.1, Fig. 10O) – that the lefts and rights of the specimens are so uniform on individuals indicates that their shapes and sizes were strongly genetically controlled as expected for display structure intended to visually segregate species in order to inhibit cross species reproduction. The images were replicated to a consistent posterior skull height to facilitate comparisons of degrees of development. The images were then re/positioned relative to one another in a gradistic manner from least prominent to most so, until an order was arrived at, and each specimen was scored from 1 to 15. These results are very approximate because although differing degrees of prominence are very real – the boss of HMN MB.R.91216 (Fig. 10E) is clearly much less enlarged than that of MOR 008 (Fig. 10C) – the fine gradations involve a degree of judgement, which are impacted by differing lighting in the images, as well as extensive differences in the form of the bosses, and other factors. Possibly 3-D scanning of actual specimens can be used to obtain better results in the future. The quantitative results were used to produce the following ratings; (NP), fairly prominent (FP), prominent (P), and very prominent (VP). There are multiple examples of each grade, at different levels of the TT-zone. The degree of rugosity of the nasal ridges, especially the midline profile, is also assessed in a similar manner with similar caveats as fairly smooth (FS), fairly rugose (FR), rugose (R), very rugose (VR), and extra rugose (ER).

**Fig. 10.**
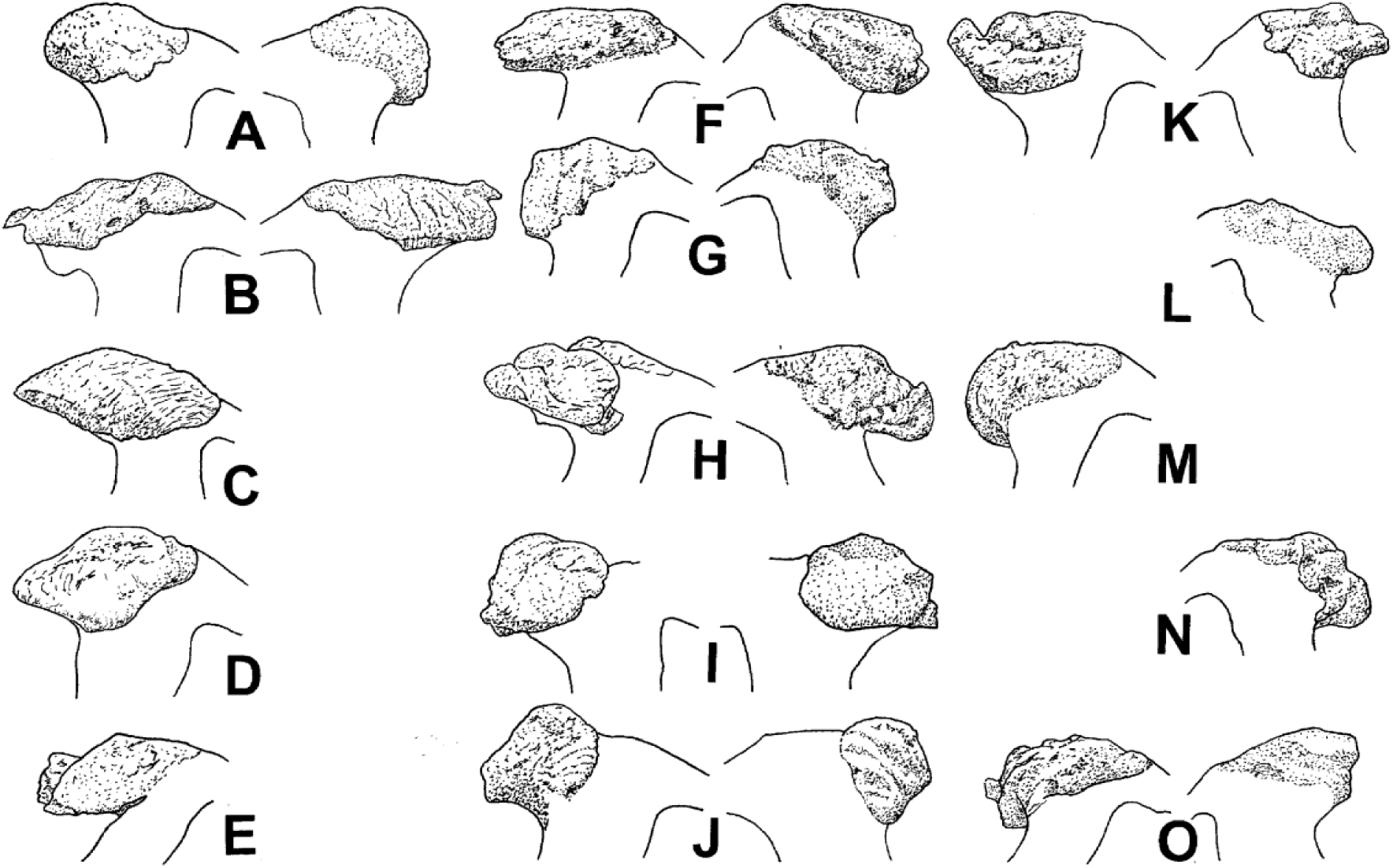
A-O Left and/or right postorbital bosses in lateral view of *Tyrannosaurus*, presented at an approximate constant skull size, in each species the specimens are presented in order of general decreasing size of the individuals. Lower TT-zone *T. imperator*: **A** Samson; **B** FMNH PR2081; **C** MOR 008; **D** SDSM 12047; **E** HMN MB.R.91216; **F** RGM 792.000; **G** MOR 1125. **H** *T.* incertae sedis AMNH 5027. Upper TT-zone *T. rex*: **I** RSM P2523.8; **J** UWBM 99000. Upper TT-zone *T. regina*: **K** NHMAD “S”; **L** LACM 23844; **M** USNM 555000; **N** MOR 980; **O** TMP 81.6.1.

There is little difference in the shape of the nasal ridges in the sample of large specimens. While no nasal is truly smooth, roughness ranges from low to quite high, with MOR 008 alone earning a rating of ER – which is about as maximally rugose as seen in most other tyrannosaurids, but does not match that seen in *Alioramus* (Brusatte et al. 2012; Paul 2016). In the great majority of examples there is very close to good correlation between the roughness of the nasal with the prominence of the postorbital bosses. But in four cases there is a large disparity with the nasals being seriously rugose while the postorbital protuberances are not especially prominent, or the reverse, FMNH PR2081 being especially notable in having VP bosses and a FS nasal. There is some tendency for smaller specimens to have smoother nasals than larger examples and big MOR 008 is exceptionally heavily textured, but there are a number of exceptions such as RGM 792.000 and FMNH PR2081. While VP and EP nasals decorate skulls assigned to robust *T. rex* and *T. imperator*, known gracile *T. regina* appear to lack such, whether that constitutes a taxonomic feature is not clear. There does not appear to be significant taxonomic information contained in large *Tyrannosaurus* nasals.

The most developed orbital bosses are always present on very large specimens, but some large individuals do not have VP or even P/FP bosses, with one having an NP boss. This indicates that although ontogeny of course played a role in the degree of development of the display structures, it was more complicated than just that. *T. imperator* bosses range from NP to VP, *T. regina* from NP to P, and the two *T. rex* are P or VP. All rankings are found at differing levels of the TT-zone.

Among the lower TT-zone *T. imperator* documented female MOR 1125 (Schweitzer et al. 2016) is a medium sized robust that was reproductive but probably not fully mature, and it has an FP protuberance (Figs. 1H, 10G). Having an even less prominent boss, similar sized HMN. MB.R.91216 is another candidate for a female, although immature male is also possible. Although very large, and with a P boss, that of Samson is relatively short, leaving open the possibility it was a fully mature female. The rest of the very large individuals with big spindle bosses are probably mature males. In *T. regina* 3 large specimens have P bosses, but another only a NP, suggesting the first are mature males and the later a mature female. More modest sized HMN MB.R.91216 has an FP boss, so it may have been a female. For *T. rex* that very large RSM P2523.8 has a somewhat more prominent boss than less massive UWBM 99000 favors the former being a fully grown male (Fig. 1B,G), the latter a female with immature male also possible.

If all upper TT-zone *Tyrannosaurus* are the one species *T. rex*, with the robusts and graciles each representing a gender, then it would be expected that as the species evolved from lower TT-zone *Tyrannosaurus*, the specimens with prominent postorbital bosses -- probably males -- would end up limited to one morph or the other. If instead they are two sympatric species, then some skulls of both the robust and the gracile species are predicted to exhibit prominent bosses. Note that this result is not dependent on sample size because what is on hand establishes the lack of correlation between the postorbital bosses and skeletal build predicted by one species in the high TT-zone in favor of two. The apparent preserved pattern in which both robusts and graciles have both poorly and well developed postorbital display structures is instead most accord with the latter circumstances.

Yet stronger and more compelling support for the three species hypothesis is found in the distinctive and varying shapes of the bosses. The topography of the shapes are documented in Figure 10 by direct tracings of photographs. The monospecific hypothesis predicts there should be just one basic type of boss distributed evenly through the TT-zone in the genus, perhaps with some dimorphic variation found between robusts and graciles. Two chronospecies predicts a more complex scheme with at least two differing boss morphologies stratigraphically separated. Three species would be indicated by at least three differing boss forms. The latter is true. A number of low TT-zone *Tyrannosaurus* have very distinctive, anteroposteriorly long and low, spindle shaped postorbital prominences that extend from the lacrimal contact back to nearly or all the way onto the anterior section of the squamosal process of the postorbital, these do not project much above the dorsal edge of the skull (Fig. 10B-D,F). Such bosses are not yet known from the high TT-zone, and all the specimens that have the spindles are assignable to *T. imperator*. On the skull roof small medially oriented anteromedial projections of the postorbital prominences are oriented towards the skull midline, but are far from contacting one another. In sharp visual contrast, two high TT-zone specimens exhibit very different, semi-circular knob/disc shaped bosses that project well above the dorsal rim of the skull, and are limited to the frontal process of the postorbital, with a large space between the posteriormost edge of the protuberance and the squamosal process (Fig. 10I,J; Fig. 12 in Persons et al. 2019; www.rescast.com/case-studies/scotty-the-t-rex; www.trendsmap.com/twitter/tweet/1431126894997614593; www.fossilcrates.com/products/totaltrex; commons.wikimedia.org/wiki/File:Tufts_love_rex.jpg). There are not significant anteromedial projections of the prominences. These similarly exceptional knob bosses of RSM P2523.8 and UWBM 99000 (Fig. 1B,G, 10I,J) reinforce their placement in the same species, that being *T. rex* – that the two skulls especially the latter are fairly new is a factor in their remarkable shaped bosses not being recognized earlier, the absence of postorbitals in the *T. rex* holotype may have hindered appreciation of this anatomical situation to date. No low TT-zone specimens have such orbital projections in shape, elevation, or extreme anterior placement - for that matter no other tyrannosaurid, or theropod, has such idiosyncratic eye catching postorbital displays. *T. regina* bosses are more irregular in configuration, none having either the spindle or disc shape, the bosses always extend more posteriorly than in *T. rex*, sometimes onto the squamosal process (Fig. 10K-N). The largest and presumably most mature and probably male *T. regina* has bosses that have something of a brimmed hat in picture appearance (Figs. 1C, 10K). In at least some cases the anteromedial projections are well developed like those of *T. imperator*.

Differences between the shape ends of *Tyrannosaurus* ornament bosses were far from subtle and would have been readily visible to the living archosaurs. As explained in the section on determining paleospecies, it is very probable that the keratin sheathes covering dinosaur postorbital bosses moderately expanded and enhanced the ornaments’ size while largely replicating their bony cores. This presumption is applied to life restorations of *Tyrannosaurus* heads representing large males of each of the three species (Fig. 11). Lacking examples other than the prior, the possibility that the bosses bore much larger soft coverings that dramatically altered their form is too low to warrant serious consideration much less illustration, and even if such occurred it is very unlikely that the coverings of such very different bone foundations happened to coincide to produce the same final external appearance between the morphotypes and stratigraphic horizons.

**Fig. 11.**
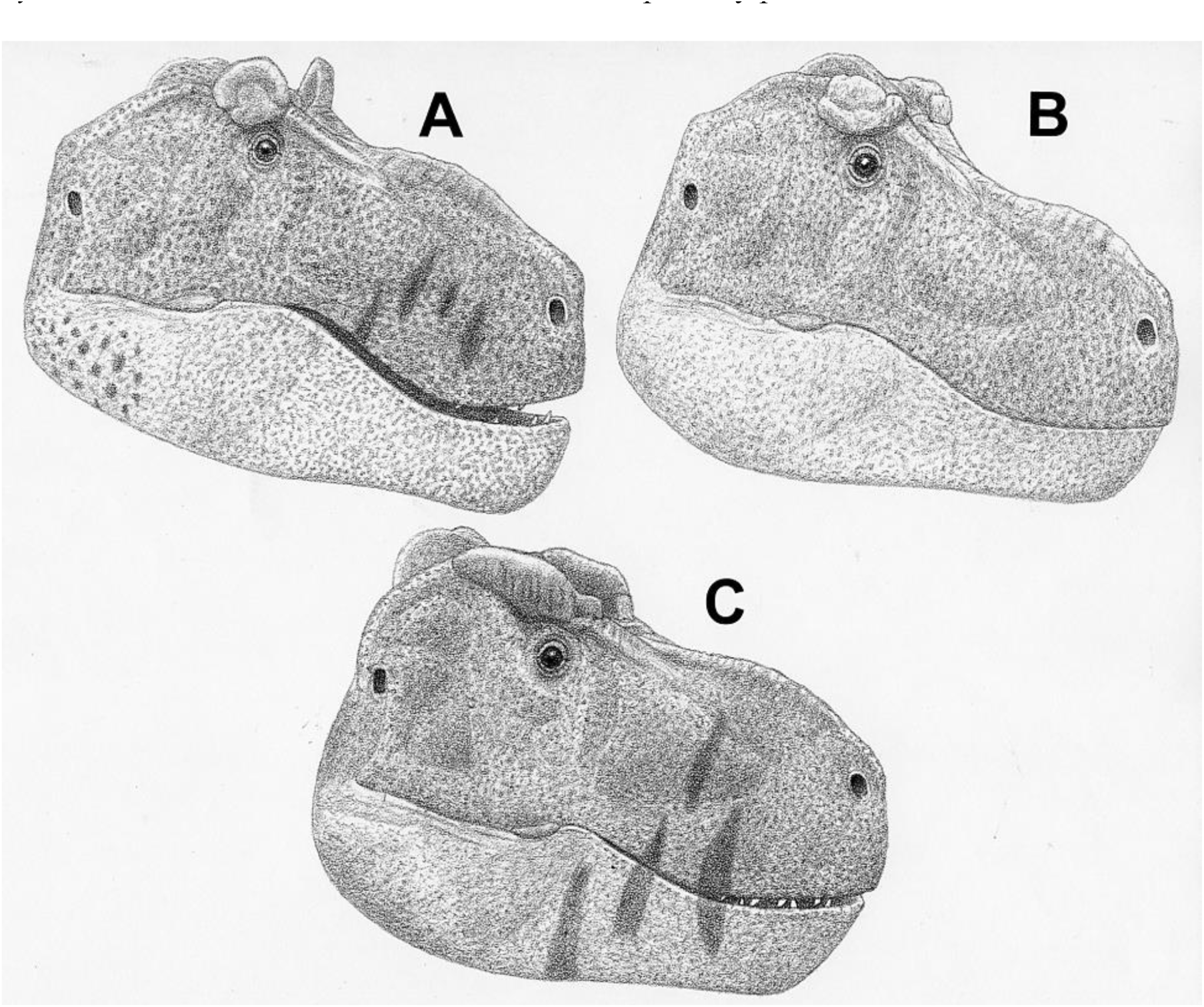
Life restorations of heads to same approximate length and scale of large, presumably male *Tyrannosaurus* showing the highly divergent species specific differences in the postorbital bosses. Bosses carefully proportioned in size relative to skull, and in shape, after being enlarged a modest amount with reconstructed keratin sheaths (as per Fig. 1B-F), scales and color patterns speculative, ornament color patterns kept simple to emphasize surface topographic differences. Upper TT-zone: **A** *T. rex*; **B** *T. regina*. **C** Lower TT-zone *T. imperator*.

**Fig. 12.**
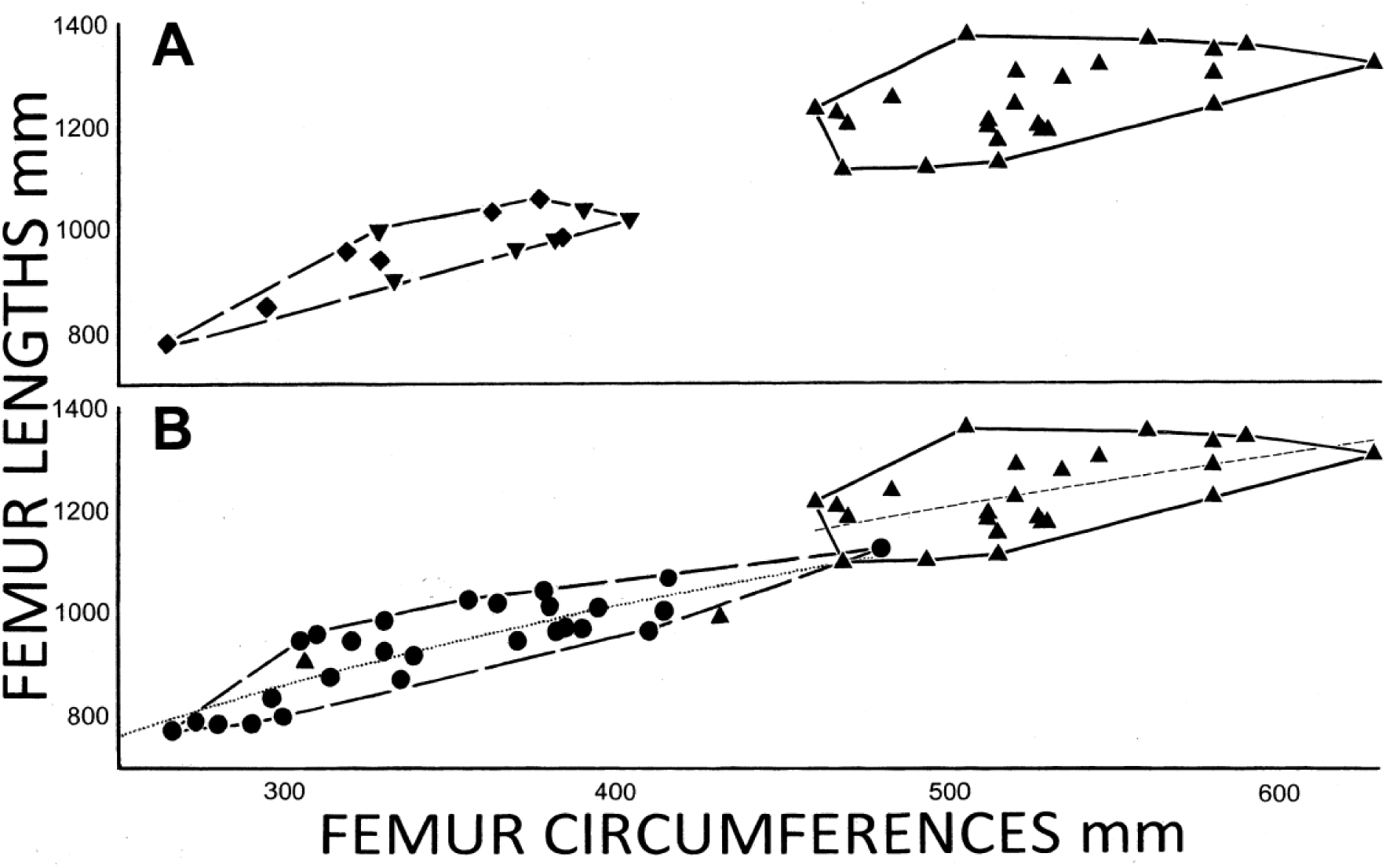
Femoral proportions of all large *Tyrannosaurus* (triangles) compared to **A** *Gorgosaurus libratus* (diamonds) and *Daspletosaurus torosus* (inverted triangles), note that two immature *Tyrannosaurus* specimens are not contained in the least area polygon that incorporates only large specimens of the genus; **B** All tyrannosaurids (circles) aside from those from the TT-zone, modified from Fig. 4C in Paul et al. (2022) including regression lines for two groups. All femora over 700 mm long.

As a result of this analysis, what was orderless in the context of monospecific *Tyrannosaurus* without stratigraphic examination is now a strong pattern of species specific ornamentation distinguishing three species, two of them contemporary. Large, probably male individuals of each species sport bosses distinctive from the other two taxa. The bosses were horizontally elongated spindles in basal *T. imperator*, in latter *T. rex* the bosses are very optically different, anteriorly limited, vertically elevated subcircular discs in the shape of knobs, in coexistent *T. regina* the bosses remain more horizontally elongated and readily distinguishable by late *Tyrannosaurus* eyes (Fig. 11).That there is considerable variation in boss topography within at least two of the species (Fig. 10A-G,K-O) is fully in line with the similar intraspecific divergences seen in species display structures in a host of species, including the tremendous differences in brow horn length and orientation in *T. horridus* (Fig. 7A-G), and the tusks within proboscidean species, and the perpetual search for atypical trophy antlers by hunters. While *Tyrannosaurus* boss shape differences in the best developed examples of larger specimens the three taxa are broadly similar to those that distinguish the males of modern sibling species, they in their numerous considerable differentiations exceed the simpler and lesser variations normal between the genders of extant species, so gender based dimorphism is not as explanatory as is speciation. And just as dimorphism is not a viable simple hypothesis for robust versus gracile proportions because the latter are absent in the low TT-zone, the absence of anteriorly placed knob bosses in any known low TT-zone skulls currently excludes the possibility of dimorphism between the bosses of *T. imperator* and those of later *Tyrannosaurus*. That the differing bosses of upper TT-zone sympatric *Tyrannosaurus* were dimorphic is also improbable because that condition should result in differing sizes of otherwise fairly similar display structures in the genders with those on males being larger, not in the dramatically different knob versus hat shapes that are examples par excellence of the divergent anatomical topographies that evolve as species specific visual cues. Nor is how only a quarter of the high placed skulls bear the prominent knobs in line with an expected 50/50 male/female ratio. To the above can be added the lack of precedence in tyrannosaurids species for differing postorbital display bosses between the sexes. Instead the situation most similar to upper TT-zone robust *T. rex* and gracile *T. regina* is that of robust *D. torosus* and gracile *G. libratus*, with the latter two co-inhabitants of the same ecospace each proffering easily distinguished orbital displays between the taxa.

If well developed, prominent postorbital bosses of distinctive shapes are male display characters then it is expected that females would lack such, at least in terms of size. It is therefore notable that among lower TT-zone *Tyrannosaurus* the known female, MOR 1125, lacks the full developed spindle that appears to have visually distinguished male *T. imperator,* being a robust (Fig. 1E,H). If the upper TT-zone *Tyrannosaurus* females are also robusts then the specimens that bear the most visually attention getting bosses should be graciles, but the two that have such displays are robusts that are probable males. The modest size of the knob boss on the modest sized and potentially female UWBM 99000 (Fig. 1G) is most compatible with *T. rex* being robusts that bore the supraorbital discs whether male or female, leaving them visually distinct from more gracile *T. regina* that lacked the distinct display knobs of its contemporary. If the inferior hypothesis that *T. regina* is a junior synonym of *T. rex* is accepted, then presumably the latter with their tall bosses are robust males and the former gracile females in a peculiar pattern not observed in other tyrannosaurids or dinosaurs.

Other known tyrannosaurids do not bear postorbital bosses that are highly similar in all respects to those of any of the *Tyrannosaurus* species (Maleev 1955; Russell 1970; Currie 2003a; Hurum & Sabath 2003; Carr & Williamson 2010; Carr et al. 2011, 2017; Brusatte et al. 2012; Lu et al. 2014), which are distinctive from one another. Therefore, each of the three types qualify as an autapomorphy among the family and within the genus.

A complex statistical shape analysis of the bosses is beyond the capability of this work if it possible at all, but the distribution of *Tyrannosaurus* boss morphology is bimodal in that no member of the genus from low in the TT-zone has the elevated discs of RSM P2523.8 and UWBM 99000, nor does any documented high placed example have the spindles of RSM P2523.8, MOR 008, or RGM 792.000, and no specimens assigned to *T. rex* or *T. imperator* have a boss like that of NHMAD “S”.

The profile-skeletals in Figure 1 are the first set to show all *Tyrannosaurus* skeletons with preserved postorbitals accurately sporting their varying supraorbital display structures. Aside from necessitating a significant revision in the appearance of *Tyrannosaurus* and its species *T. rex* especially (Appendix 1), the skeletals help show that the species, males in particular, would have been easy to tell apart when living animals with a quick visual glance, as is further indicated by life restorations of the species heads (Fig. 11). *Tyrannosaurus* shows stronger differentiation in bone based specie specific features than are present in within other predatory genera of tyrannosaurids (Fig. 6), theropods, *Varanus, Panthers, Canis,* (Figs. 2-4), *Stenopterygius,* and *Pliosaurus*, predaceous monitors, felids, and canids, and comparable to herbivores such as *Triceratops* in terms of the nasal horn differentiation that is a major critical species marker in that genus (Fig. 7), as well as some mammalian herbivores (Nowak 1991).

The bosses confirm the species identifications of three skulls that had not been species placed in Paul et al. (2022), but were by additional skull proportions above – that further eliminates the objections by Carr et al. (2022) regarding this matter.

The inherently very weak alternatives if only the species *T. rex* is preserved demands an incoherent level of extreme individual variation, which lacks logical evolutionary cogence as it somehow produces a significant pattern over time that is entirely in accord with and evidence for species divergence in three taxa. The degree to which the anatomical/stratigraphic patterns observed with the current specimens do or do not hold up as new specimens come on line will help test the cranial display hypothesis presented in this analysis. Also potentially pertinent is preliminary notice of divergent final growth and size patterns in *Tyrannosaurus* that do not conform with stratigraphy or robustness, and that this may represent cryptic sexual dimorphism (Jevnikar and Zanno 2021). It will be interesting to see to what degree that evidence conforms to the above not at all cryptic results.

Continuing to pose an interesting outlier is AMNH 5027 (Fig. 1E; Fig. 10H). It’s P grade postorbital bosses do not fit readily into the topology of any of the other variants. The main boss is anteriorly placed somewhat like those of *T. rex*, yet are somewhat more horizontally long, and feature a projecting lip along the orbital edge, while the boss narrows down to a ridge atop the dorsal rim of the postorbital. The bosses do not project well above the dorsal rim of the skull as in *T. rex*, the anteromedial projections are present. The bosses are somewhat more similar to those of *T. regina*, but remain distinctive. Rediagnosing the species by forcing *T. regina* into *T. rex* (systematics section) does not aid in the placement of 5027.

The new characters have been added to the revised and expanded diagnoses for the three species in the systematics section, which now incorporate 13 diagnostic characters.

Taxonomic implications – Opposite the simple and fairly consistent supraorbital display arrangement predicted by monospecifity, correlating the highly variable topography of *Tyrannosaurus* preorbital bosses with stratigraphy and species reveals the anatomically and stratigraphically complex pattern that is the hallmark of the identification and diagnosis of sibling species. Of the modest but effective level expected in closely related predators lacking garish display ornamentation. With the variation exceeding that present on the skulls of many predatory genera, and matching those of some herbivore genera. And does so in the manner expected in and best explained by three species, with the differing configurations of the bosses reinforcing the assignments of the skulls to specific species based on other parameters. What once did not make cogent sense now does, and the long noticed but never explained variability of *Tyrannosaurus* skulls is readily explained as due to their representing different species. The criticism of the multispecies hypothesis that differing species exclusive display features are absent in *Tyrannosaurus* is now falsified, that belief being the result of the failure to rigorously test the status of *T. rex* with the substantial data that is available. The already strong preponderance of evidence for three over two over one species is correspondingly greatly enhanced by the identification of species ocular directed devices in each of the taxa based on that data. Which also allows provisional identification of genders within the three species (these being listed in Table 1). With species identification displays being a classic defining attribute of that taxonomic level, the bosses alone establish that *Tyrannosaurus* was not just *T. rex*. The hypothesis of 3 species, with all the graciles being in one of them, has made it possible to tease out these patterns, something the one species proposition is incapable of. Assuming 1 species, or 2 chronospecies, obscures the dimorphism because it fails to explain why both robust and gracile specimens sometimes brandish well developed orbital displays while others less so.

Paul et al. (2022) missed the opportunity to describe the relationship of the supraorbital bosses to the species of *Tyrannosaurus*. Because Carr et al. (2022) focused on criticizing Paul et al. (2022) rather than go beyond to investigate the broader situation, they did too. That Paul et al. (2022) laid the foundations for exploring multispecific *Tyrannosaurus* made these novel results serendipitously possible.

### More *Tyrannosaurus* Sex

Differences in tyrannosaurid robustness may have visual species specific attributes. The depth of the maxilla and dentary relative to their lengths can result in a deeper or shallower head that can have species identification and competitive implications for the living animals. Same can be true for a heftier versus a more lithe body supported by a corresponding difference in the robustness of the skeletal framework. The latter could parallel the difference between the body build of the African black rhino and the somewhat heftier white rhino. These effects may be subtle, which is appropriate for predators than could not repurpose major developments in weaponry evolved for in combat performance for objectives related to sexual selection. The status of the anteriormost dentary teeth was not likely a reproductive indicator because it was probably not especially visible, all the more so if the teeth were normally covered by lips and gums (Fig. 11; as per Paul 2018).

Taxonomic implications – The amount of species ID divergence within *Tyrannosaurus* appears to have ranged from visually readily apparent via the supraorbital bosses to subtle differences in head and body proportions.

### Only One Major Skull and Skeleton Still Systematically Adrift

As noted by Paul (1988) and Paul et al. (2022 Suppl.) the complete AMNH 5027 skull is significantly distorted, being crushed so the top of the skull is pushed subtly but substantially ventroposteriorly and to the right relative to the lower sections. As a result, the left and right sides do not match in configuration, with the right more altered, the right orbit being slanted too posteriorly progressing dorsally, and the right squamosal and quadratojugal are disarticulated. The upper teeth are more procumbent than normal in the genus because the maxillae are dorsoventrally crushed in tune with the rest of the skull, leaving it probably more shallow than it was in life, making it appear more gracile than it actually was. What can be reliably measured scores as borderline to gracile, but the limb material most critical to the proportional portion of species determination is missing, and the anterior dentary teeth are currently not accessible to confirm or refute the presence of two small incisors (Bakker in Larson, P. 2008). As just discussed the postorbital bosses are not in accord with those of other species. The quarry has been flooded by the Lake Peck reservoir, very probably permanently precluding determination of its stratigraphic placement, and any hope of finding the rest of the skeleton. It is possible that 5027 is a distinct species, as per the *T.* “x” hypothesis (Bakker in P. Larson 2008), albeit when applied only to the one specimen (all the other specimens considered potential *T.* “x” in Larson are robusts assignable to *T. imperator*). It cannot be overemphasized that no TT- zone *Tyrannosaurus* that is as anatomically and stratigraphically deficient as AMNH 5027 should be used as a holotype – all current holotypes include an intact femur for instance and that needs to be true for future holotypes any species in the genus -- so testing *T.* “x” requires verification by additional specimens. In any case 5027 may be a transitional form between the other species (Van Raalte pers. comm.), perhaps between *T. imperator* and *T. regina* in that its bosses have features of both, especially if 5027 is from the mid-levels of the TT-zone. Another possibility is that it is a hybrid, perhaps of the just mentioned two species if they overlapped in time. It may prove impossible to ever taxonomically place this specimen.

Taxonomic implications – The *Tyrannosaurus* data base for assessing species is now sufficiently well-developed to assign nearly all major skeleton and/or skull specimens with varying degrees of confidence to one of the three proposed species, as well as a large number of highly incomplete remains. *T.* “x” if it exists consists of only one specimen to date, the stratigraphically and systematically ambiguous AMNH 5027.

### The Comparative Cases of *Tyrannosaur*us and *Triceratops* Multiple Species Studies

Paul et al. (2022) cited Scannella et al. (2014) as setting a standard that could not be approached by the former study. Conducting this analysis found that was an exaggeration in that the studies are actually more comparable to one another. The following considers both the Paul et al. (2022) study, plus that combined with the above new data and analysis.

Both projects center on about two and a half dozen large subadult and adult specimens of varying completeness that are stratigraphically correlated. The last factor is somewhat superior in Scannella et al., but the difference that that creates in the results is not critical. Paul et al. includes nearly all major large *Tyrannosaurus* specimens from the entire geographic TT-zone, the *Triceratops* study is much more limited in that regard, and as a result does not capture as much of the full available variation in the known fossils of the ceratopsid as does the work on the tyrannosaurid which is close to complete as it can be at this time in that regard. In both analyses the number of specimens on hand is higher in upper TT-zone levels than in the lower level. Scannella et al. include estimated dimensions to a greater extent. Paul et al. examine 8 character dimensions to the 6 in Scannella et al., both expressed as fine gradations rather than simplistic over or under the median value rankings. The former analysis is more whole body in that it incorporates cranial including mandibular features, as well as postcranial parameters that tend to produce similar results, not just the upper cranial features of the ceratopsid in which only rostral features have proven determinative. A visual comparison of Fig. 6 in Paul et al. versus Fig. 2 in Scannella et al. shows that of the 8 *Tyrannosaurus* ratios 7 show trends over time, compared to 5 in 6 ratios and angles for *Triceratops* – but that declines to 4 in 6 when the premaxilla angle data from additional specimens is factored. In this paper’s Figure 9 12 in 14 ratios have trends. In both genera there are sometimes considerable variations in the ratios for a given character in a given species, occasionally involving statistical outliers within a species, resulting in considerable ratio overlaps between the proposed taxa, with bimodality correspondingly not being present in those cases. As a result, clear cut character separation between the proposed species and bimodality is not the norm, although it does sometimes occur in both genera, albeit not usually between all three species regarding a given individual character. Cranial display structures suitable for species designation are present in both genera. In *Tyrannosaurus* one of the species, the atypically gracile *T. regina*, exhibits a persistence without exceptions in proportional extremes not seen in the *Triceratops* sample. The exceptional gracility of the femora of *T. regina* is an apparent autapomorphy relative to other *Tyrannosaurus*, and to the Tyrannosauridae. This is further revealed by how the two juveniles that can confidently be assigned to *Tyrannosaurus*, robust USNM 6193 and even the apparently gracile LACM 23845, both plot in or near the general tyrannosaurid pattern, and are well below the adult *Tyrannosaurus* graciles (Fig. 12B). The same is true of the absence of two small anteriormost dentary teeth in in the last *Tyrannosaurus* vis-à-vis the *T. imperator* norm and more basal tyrannosaurids. Species diagnoses for predator and prey are correspondingly often qualified and characterized by trends rather than absolutes, as can be seen when comparing the diagnoses for the species of the two genera in the systematics section. While *Triceratops* species do not show autapomorphies, *Tyrannosaurus* species do regarding the atypical gracility and the distinctive forms of the postorbital bosses. The two studies agree that stratigraphic separation of key characters precludes sexual dimorphism as an explanation for the observed circumstances in certain cases.

With a large sample of more basal tyrannosaurids available, the *Tyrannosauru*s studies can do what Scannella et al. could not, quantitatively compare the variation in earlier species to the TT-zone taxa that exposes the very unusual variability of the last *Tyrannosaurus* during the last few hundred thousand years of the Cretaceous.

That in Paul et al. and Scannella et al. the basal TT-zone taxon retains ancestral features from recent earlier clade members, and then evolves new, derived anatomies, is powerful evidence for classic Darwinian speciation – involving three species according the data on hand in both analyses. The early outlier observed in *Tyrannosaurus*, a sole specimen lacking the two incisiform teeth widely present in earlier tyrannosaurids and *T. imperator* in favor of the only one common to upper TT-zone *Tyrannosaurus*, is not serious evidence against intrageneric speciation because it is paralleled in *Triceratops* species evolution in its wide variation in the angle measurement of the upper posterior medial border of the premaxilla in the basal early *T. horridus*. Interestingly the two genera then evolve in opposite manners – in *Triceratops* variability *declines* progressing towards the end of the Mesozoic, in *Tyrannosaurus* is *rises* dramatically. In the herbivore the mode of speciation appears to have been anagenetic with only one species extant at a given time – although the possibility of more than one contemporary intragenera species cannot be ruled out in the low and middle TT-zone. In its hunter the situation looks more complicated, with the anagenetic versus cladogenetic evolution of two contemporary species from one earlier taxon being less clear in its pathways.

The two sets of work share a level of data availability and sophistication above the norm for dinosaurs. This is logical because both focus on popular and correspondingly extensively searched for dinosaurs from perhaps the most heavily researched dinosaur fossil beds set in two prosperous and peaceful nations. That in part because the TT-zone is the last before the great extinction and therefore of special importance in evolutionary and paleobiological sciences. Being so alike in so many regards, both are similarly greatly superior as intragenera species assessments to the Carr (2020) paper that was not about that particular subject in the first place.

Taxonomic implications – The data content and analytical quality of Scannella et al. (2014) that Carr et al. (2022) correctly cite as a positive example, and Paul et al. (2022) and herein, are relatively close, with both enjoying advantages and disadvantages vis-à-vis one another. The ceratopsid study may have a noncritical edge over the tyrannosaurid because of its more detailed stratigraphy. And an inherent advantage enjoyed by research of *Triceratops* species is that the genus sported the garish display enhancing horns that often facilitate identification of species, that these are subtler in the predator that fed upon it results in a somewhat finer level of speciation that is well marked by the postorbital bosses. In addition, *Triceratops* appears to have been evolving a little more rapidly than its predator, and because the species were not contemporary hybridization would not have muddled differences between taxa as is likely to have occurred between *T. rex* and *T. regina*. Advantages for Paul et al. (2022) include the inclusion of postcranial as well as cranial anatomy from all accessible major specimens, the consistent extremities of the proportions of one of the latter species and its strong divergence from the basal species of the genus, and possibly some autapomorphies. The many parallels between the two works means that evidence presented in both that the two genera evolved away from their ancestral states into new species over the span of the TT- zone is compelling for both. And that the results of Paul et al. need to be taken with more serious equanimity than they have been. As the lateral geographic stratigraphic correlations of the TT-zone improve and the percentage of *Triceratops* specimens being stratigraphically correlated rises to all specimens with adequate quarry data, then the horned dinosaur will begin to enjoy a permanent larger sample size lead over its apex predator.

### There Are Enough Characters in Paul et al. to Diagnose *Tyrannosaurus* Species

A comparison of the diagnoses for the species of *Tyrannosaurus* and *Triceratops* presented in the systematics section demonstrates how the number of characters is somewhat higher than that for the latter examination, which is widely accepted as valid. The same is shown comparing Fig. 2 in Scannella et al. (2014) to Figure 9 herein. It is not clear if a larger set of characters will be markedly more informative, such not being the norm when sorting out sibling species.

Taxonomic implications -- The number of characters utilized in this examination and Paul et al. (2022) is typical for studies on dinosaur species, and well within the range of those used to diagnose many fossil tetrapods.

### The Characters Are by No Means All Minor

The differentiation in femoral robustness in the single genus *Tyrannosaurus* from the last few hundred thousand years of the Mesozoic being half again as great as all other Tyrannosauridae taxa spanning a dozen or more times longer span of time put together (when smaller juveniles are excluded as per Paul et al. 2022) is a distinctive evolutionary development that is far from taxonomically minor. All the more so because it is a major divergence from the ancestral tyrannosaurid condition both in the sudden unprecedented onset, and in being driven by an atypical autapomorphic burst of gracility (Fig. 12B; see Fig. 6C in Paul et al. [2022] for full curves). And that when the shift to slenderer proportions is expressed in much of the skull and skeleton in features minor and major, and represents a reduction of bone strength in a genus known for the opposite. To that can be added the visual species level display organs provided by the distinctive autapomorphic orbital bosses.

Taxonomic implications – The characters range from minor to major -- the latter including two features of the strength of the maxilla and the robustness of the femur, and the orbital bosses – as they do for *Triceratops* and many other intrageneric sibling paleospecies. It is when combined that they consist major evidence of significant evolutionary species level developments.

### The Sample Size is Better than Normal for Dinosaur Species, and Replication Problems Have Been Exaggerated

The sample of large specimens in this examination and Paul et al. (2022) is larger than normal for multispecific dinosaur genera, and matches or exceeds that for many fossil tetrapods. The size of the sample is dependent upon inclusion of all specimens for which data is available. Omitting a large set of remains as per Carr (2020) advocates and does, and Carr et al. (2022) advocate and do not do – after criticizing Paul et al. (2022) for not practicing due scientific and ethical diligence for using BHI specimens and X-rex they then used our entire femoral data set - is a major evidence exclusion. A game of pretend that when actually practiced severely reduces the ability to test the monospecies versus multispecies hypothesis, to the degree that the results will be too impaired to overturn the multispecies *Tyrannosaurus* hypothesis that is not the automatic inferior alternative – for example the most robust and gracile known femora are/were both BHI specimens, 6248 and NHMAD “S” (x BHI 3033). Data replication can if necessary usually be achieved via casts and photographs. That Carr et al. (2022) used those and other private specimens because they had to maximize their *Tyrannosaurus* data set indicates that they do not actually consider the replication issue to be seriously critical. That some of the data in Carr (2020) has replication difficulties as described above further indicates that criticism of use of private specimens on that basis has been inconsistent and exaggerated. Further note that a number of technical papers have utilized private *Tyrannosaurus* specimens (Bates et al. 2009; Hutchinson et al. 2011; Sellers et al. 2017). That said private possession of major fossils does create problems. NHMAD “S” is the most optimal holotype for *T. regina*, but its private status during the process of producing Paul et al. (2022; future status uncertain) precluded that possibility. It is important that the total *Tyrannosaurus* sample includes very large specimens from the lower and upper TT-zone, and from all three proposed species (Fig. 1B,C,E), doing so effectively precludes differences in ontogeny and size providing explanations for the variations in proportions and teeth.

Whether a dramatically larger data set will produce a major alteration in taxonomic conclusions is possible but by no means certain. Consider whether a much larger sample is most likely to overturn the basic results of Scannella et al. (2014), or tweak them? A dinosaur genus that does have a far larger sample of fossils is *Coelophysis* via the Ghost Ranch Whitaker quarry that contains many hundreds of usually articulated skeletons (Rinehart et al. 2009). But the thin walled bones including the femora are often too crushed to provide diameter or circumference data, and are still imbedded in the matrix in any case.

Taxonomic implications – Critics of multiple tyrant lizard species as per Carr et al. (2022) cannot have it both ways – assert the hypothesis lacks a sufficient sample size, and then try to test the hypothesis with a smaller sample. Either incorporate the full sample, or get out of the taxonomic research that depends upon it. The results of a sub sample will not be valid.

### The Stratigraphic Data Base is Adequate

In tune with prior media commentaries, Carr et al. (2022) take Paul et al. (2022) to lengthy task for the latter’s simple stratigraphic positioning of *Tyrannosaurus* specimens into low, middle and high bins in the geographically laterally expansive TT-zone, in comparison to the tighter metric specific placements for *Triceratop*s specimens in the geographically limited section of the Hell Creek in Scannella et al. (2014), The extensive criticism is perplexing in a number of regards. Including how Carr (2020) has been widely praised as superior to Paul et al. (2022), even though it too only uses generalized low, middle and high stratigraphic categories for cross data analysis for far fewer specimens. Carr et al. (2022) accept broad stratigraphic bins for the actual correlative taxonomic work. That Paul et al. (2022) cite and use the data in Carr (2020), and as a result the data sources he utilized, was not noted in Carr et al. (2022).

To be specific. The Paul et al. (2022) multispecific *Tyrannosaurus* hypothesis makes the gross level proposition that the species of the genus in the lower portion of the TT-zone that lasted a very substantial geological time did not survive into the upper zone where there were new species, with a probable but uncertain mix in between -- this is illustrated in Figure 1b in Carr et al. (2022) in which each species spans about two thirds of the TT-zone/ Paul et al. (2022) do not offer a more time precise theory in which there was one species in the lowest fifth of the formation, a new sequential one in the next fifth, yet another in the fifth after that, and two more anagenetic species in the final two fifths. It terms of geotime it is a very simple either/or stratigraphic issue of lower and upper, with an apparent -- but concerning the validity of the hypothesis not critical -- species overlap. As per perfection being the enemy of sufficient to get the core paleotaxonomic job done, placing specimens in such time broad zones simply does not require metric precision.

Regarding the Canadian sample, there is no significant vertical placement issue relative to the Paul et al. (2022) lower and upper species hypothesis. As explained by Paul et al. (2022 and refs therein, also Eberth and Kamo 2019; Mallon et al. 2022), the technical literature demonstrates that all the Canadian TT-zone formations are from the upper half of TT-zone. It follows that, according to the data published in all pertinent academic studies, no *Tyrannosaurus* found in Canada is from the early portions of the TT-zone, so all Canadian *Tyrannosaurus* have to be assigned to the upper sections. This widely accepted reality is not directly mentioned by Carr et al. (2022), although it is in accord with the data cited by Carr (2020). and references cited by the latter and in Carr et al. (2022).

Whatever geostrata issues if any may pertain to the subjects under consideration apply to the United States sample, that extending from the bottom to the top of the TT- zone. While Carr et al. (2022) negatively critique without positive evidence the stratigraphy of Paul et al. (2022), they do not actually challenge the placement of a single specimen American or otherwise in the latter with hard data. That is not surprising because the Paul et al. (2022) data set is, despite the imprecise complaints, actually well founded as to the basic positioning of the specimens. The probability of any known adult gracile proving to come from the lower TT-zone is low. Same for a known spindle boss from the upper layers, or a knob boss from low in the zone.

The situation with the spindle bossed, two small incisor toothed, robust *T. imperator* holotype well illustrates the American TT-zone situation. Carr et al. (2020) sharply criticize Paul et al. (2022) for relying on N. Larson (2008) and pers. communications for much of their stratigraphy. FMNH PR2081 is stated as coming from just 5 meters above the base of the Hell Creek in N. Larson (2008). This is in basic accord with the stratigraphy that Carr (2020) and Carr et al. (2022) cite, as do Paul et al. (2022 because that cites Carr (2020). However, P. Larson personally informed Paul that being from the shallow eastern portion of the Hell Creek Formation probably precludes the specimen from being from being very low in the TT-zone, upper lower or lower middle being more plausible. So Paul et al. (2022) discussed the issue and took care to provisionally position FMNH PR2081 between the two levels as is done in this analysis. In contrast Carr (2020, Table 18) simply assigned the specimen to the lower TT-zone with no discussion of the geological complexities, perhaps being unaware of them because of a lack of pers. comm. with the person who excavated the specimen – this helps show why pers. communications are a frequently useful and oft cited norm in science, expert personal knowledge sometimes not being in the literature. That aside. it is not of critical importance exactly how many meters above the bottom of the TT-zone FMNH PR2081 was preserved. It is whether the fossil is from the lower portion or the upper, and that the possibility that the holotype of *T. imperator* dwelled in the upper third of the TT-zone is so improbable that it does not warrant consideration. The paleozoological problem is not that Carr (2020) did what he did in vertically positioning FMNH PR2081, it being sufficient for paleospecies determination. The problem is that Carr et al. (2022) then criticize Paul et al. (2022) for their geological methods when Carr (2020) was not superior and was if anything inferior, and the results in Paul et al. (2022) are sufficient for the task at hand. It follows that while raising some concerns is legitimate, the excessive criticism contained in Carr et al. (2022) was not fair and objective as well as perplexing, especially because it did not actually discredit any of the vertical placements.

Carr (2020) observed that resolving the time transgression issue for the TT-zone resulting from the eastwardly regressing interior seaway was “beyond the scope of this work, the results of which are offered here as a hypothesis for further, more rigorous testing of stratigraphic correlation.” Exactly. So why were sufficient but not exacting time correlations issues acceptable in the study that some claim shows there was only *T. rex* in the TT-zone, while doing the same was not in an analysis that stated it was based on the best available preponderance of evidence – including a much larger geological anatomical correlating data set – that discovered more than one species? A major methodological inconsistency that Carr et al. (2022) do not acknowledge or address?

Only about one out of nine of the large specimens being examined in Paul et al. (2020) and herein cannot be reasonably confidently grossly stratigraphically placed at this time; that compares to how only about one in four large specimens were stratigraphically correlated in Carr (2020). To seriously contradict the stratigraphy of Paul et al. (2022) it is necessary to actively refute an acute set of the geological positions that that study presented for specific specimens. Which Carr et al. (2022) do not do because such is apparently not possible at this time if ever. If specimens seriously contradicting the Paul et al. (2022) hypothesis from the stratigraphic end of things do exist, it is likely that they have yet to be excavated. If such happens then that will be time for a major reconsideration of the species numbers.

The critical stratigraphic items regarding the tyrant king are that graciles and knob postorbital bosses so far have not been documented to be preserved in the lower TT-zone, and going into the future are likely to prove to be at best rare compared to robusts that retain the tyrannosaurid basal stout femur condition in the lower TT-zone. Where the tyrannosaurid basal condition of two incisiform dentary teeth is nearly exclusive, and spindles bosses are the norm, while such are so far absent in younger sediments, where variation on robustness by the sampled elements is usually much higher.

Taxonomic implications – As Paul et al. (2022) explain and Carr et al. (2022) note, improving the *Tyrannosaurus* stratigraphic correlations to the Scannella et al. (2014) precision will require a large scale organizational effort over many decades. If such is possible across the TT-zone in view of the scarcity of radiometrically datable deposits, and the possibility that laterally varying habitat conditions may complicate other means of lateral cross dating. Even if markedly finer stratigraphic placement of nearly all specimens becomes possible, unless in the improbable event that the results dramatically differ from the basic information in Paul et al. (2022) and herein, the multiple species hypothesis is likely to survive. Finer stratigraphic data is detail work that may help parse out intricacies of how and why *Tyrannosaurus* species evolved, but is not necessary to establish the initial outlines as being constructed with the currently available information. Because it has not been possible to actually refute or even cast critical doubt on the geodata in Paul et al. (2022) relative to the stratigraphic accuracy needs of the not exact time critical hypothesis it contains, the criticism of it has been very excessive and not refutative.

### Tyrannosaur Teeth Have Diagnostic Value at the Paleospecies Level

Paul et al. (2022) measured the base dimensions of the anteroposterior anterior dentary teeth on a given side from either the teeth themselves, or the alveoli which are usually barely larger. Measurements were not combined from different sides (contra Carr et al. 2022). Paul et al. (2022) set the boundary between one and two incisiforms at a ratio of 1.2. Because a specimen with one incisiform has a ratio of 1.19, the ratio divide is reset at 1.25 herein (Fig. 9H), with the proviso that specimens close to this value on both sides are intermediates. Note the criticism by Carr et al. (2022) that the assignment of a ratio boundary regarding these teeth is arbitrary and therefore lacking utility because of the lack of a bimodal gap is not in accord with how Carr (2020) used a femur ratio boundary despite the absence of a bimodal separation in the sample in that study.

Carr et al. (2022) disputed a few of the measurements of Paul et al. (2022) on varying grounds. Their results for *T. rex* RSM 2523.8 differ very little from ours (Fig. 9H) – it is noted that as the upper TT-zone with the highest ratio this specimen is too marginal in to be considered to have two small incisors, and has a much lower ratio than most lower TT-zone specimens. Both 2^nd^ dentary teeth of marginally robust, possible *T. rex*, or *T. imperator*, NHMUK R7994 are about the same base diameter at about 40 mm. That is about 15-20% less than that of the right 3^rd^ tooth based on large format, high resolution photographs on both sides of that dentary (opposite tooth absent), and the 2^nd^ teeth are also somewhat shorter than the next few more posterior dentary teeth. So the Paul et al. (2022) ratio for the specimen stands, and as per that study the taxonomically marginal NHMUK R7994 has the intermediate condition in accord with a specimen from the middle TT-zone. The Paul et al. (2022) measurement for *T. regina* MOR 980 is from P. Larson (2008), the left 2^nd^ tooth is about the same height as the left 3^rd^ so it is not a smaller incisiform tooth, and even the Carr et al. (2022) results do not put the specimen into the ratio range of specimens with two properly small incisors. The right second tooth of probable *T. regina* LACM 23844 has a base about as large as is the diameter of both the alveoli behind it and the left 3^rd^ tooth, so this is another high TT-zone specimens without two small anterior dentary teeth.

Carr et al. (2022) make an interesting suggestion concerning a possible ontogenetic factor behind the configuration of the anterior dentary teeth, contending that the first tooth position is essentially lost with growth in some adults. In that case some of the measurements in Paul et al. (2022) seemingly of positions 2 and 3 are actually of 3 and 4. The Carr et al. (2022) hypothesis is plausible but not verified especially for all specimens. Because of the current uncertainty in anterior tooth counts it is not possible to entirely settle the disagreements in ratio values. But, even if the Carr et al, (2022) ontogenetic tooth thesis is correct, then their results do not show that there are high TT- zone *Tyrannosaurus* with two functional small incisiform teeth, there to date not being a single documented case of a specimen with such near the K/Pg boundary, the T. rex holotype included. The basic Paul et al. (2022) results therefore stand with only minor modification. Nor does the Carr et al. (2022) ontogenetic hypothesis explain why the reduction – which in practical terms is a functional adaptation, tooth count homology is not the critical issue (contra Carr et al. 2022) -- occurs almost exclusively in the last *Tyrannosaurus,* almost all earlier *T. imperator,* as well as early Maastrichtian and earlier Campanian tyrannosaurids dating back 10 million years and on two continents, having the two small incisors. Whether few if any of the last *Tyrannosaurus* came to have two small anterior lower teeth because tooth 2 became large (perhaps during ontogeny), or tooth 1 largely disappeared (during ontogeny), what is evolutionarily and therefore taxonomically important is that one way or another a significant shift occurred in this expressed, functional feature on a highly consistent basis, and that was probably a DNA driven bioevolutionary event that can be used to help track speciation patterns and diagnose species like any other such characters regardless of the ontogeny factor. This conclusion is not impacted by related ontogenetic factors because only similarly large specimens from differing levels of the TT-zone are compared. The Carr et al. (2022) remeasurements have not significantly altered the basic statistical results of Paul et al. (2022), there still being a very strong skew from nearly always two small incisiform dentary teeth in basal *Tyrannosauru*s, towards one functioning incisiform on the derived species. Some data tweaking may be required as more information becomes available.

Taxonomic implications – The apparent sudden and very late evolution of *Tyrannosaurus* frontmost lower teeth from the longstanding tyrannosaurid ancestral to a new derived functional condition is fully and most compatible with the speciation expected over the long evolutionary time span of the genus in the TT-zone.

### Variations in Tyrannosaur Proportional Skeletal Strength Has Diagnostic Value at the Paleospecies Level

Carr et al. (2022) compare femoral lengths to circumferences in extant birds with those of extinct tyrannosaurids. They do not cite prior examples of such being done in paleotaxonomy. This procedure was not followed by Paul et al. (2022) because it risks the possibility of comparing statistical fossil apples to modern oranges, in that the samples ancient and modern may not be comparable. It not clear whether the avian sample excludes captive birds that may exhibit atypical variation, bird femora are highly pneumatic and thin walled, the birds are much smaller, having varying lifestyles, and are fliers in which the hindlimbs provide secondary locomotion, sometimes semiaquatic. Paul et al. (2022) follow the more common paleontological practice of comparing fossil elements dimensions to those of other fossils among animals of similar form and function as discussed in the section on standards for determining paleospecies. It was of course not intended in Paul et al. (2022) nor herein to try capture and compare the full range of element variations in the actual populations of the taxa, that not being possible or necessary in paleozoology. The crucial taxonomic need is to compare the relative differences between tax and changes over time in the theropod proportions, which having tried to dismiss by the problematic comparisons to modern taxa Carr et al. (2022) do not attempt. That is best done by limiting sample comparisons to once wild living fossils of comparably arch predatory megatheropods that shared thick walled femora in hindlimbs that were the only locomotary organs and were used almost solely on land. Comparing element ratios to stratigraphic positions is a normal paleotaxonomic practice as per Scannella et al. (2014).

Carr et al. (2022) note that because the Paul et al.’s (2022) *Allosaurus* femoral sample does not include many adults it may be missing the variation that the population actually had. That paper neglects to note that the small sample of *Tyrannosaurus* large juveniles detailed by Paul et al. (2022) shows that substantial variability was present in the genus well before maturity was reached, so the absence of the same in the subadult allosaurs gives some support to lesser variability in that species vis-a-via the genus *Tyrannosauru*s.

The available fossil data finds that the distribution of the strength proportions as measured by the intraelement comparative dimensions of a number of major elements is not random in tyrannosaurids and *Tyrannosaurus* specifically (Paul et al. 2022). The normal basal condition of Asian and American Campanian and early Maastrichtian tyrannosaurids, retained in nearly all early *T. imperator*, is robust femora. Late Maastrichtian *T. rex* and *T. regina* fossils show a much wider variation of robustness in the femur and a number of other cranial and postcranial elements, driven by the sudden appearance – after 10 million years of tyrannosaurid evolution – of the much more gracile condition in *T. regina*, in opposition to the expectation of enhanced robustness in the dinosaurian giant. The latest was a major and evolutionarily systematically telling finding of Paul et al. (2022) that Carr et al. (2022) did not directly address. Concerning the Carr et al. (2022) criticism of comparing *Tyrannosaurus* femurs to those of other family species of lesser numbers, combined, observe that femoral variation in TT-zone *Tyrannosaurus* (n=24) is about twice that present in the Dinosaur Park Formation’s *D. torosus* (N=6) and *G. libratus* (n=7) femora over 700 mm, whether the latter two are considered independently or combined (n=13) (Fig. 12A) – that doubling the sample size via the combination barely changes the results suggests that the difference between the latest Maastrichtian dinosaur and its late Campanian precedents is not just the result more specimens producing more variation. The same results persist when the data for the two albertosaurs *A. sarcophagus* (n=7) and *G. libratus* are compared individually or combined (n=14) (Fig. 6B in Paul et al. 2022), and when the femoral proportions of all albertosaurs and daspletosaurs are united (n=20) which is approaching the *Tyrannosaurus* sample. That proposition is confirmed by how the divergence remains so much larger in the single genus *Tyrannosaurus* than all tyrannosaurids from two continents over 10 million years combined, being three quarters greater when all the latter with femora at least 700 mm (n=27 for nonTT-zone, non*Tyrannosaurus*) are compared (Paul et al. 2022; Fig. 12B). Ergo, expansion of the sample size of non*Tyrannosaurus* to equal to that of the one genus does not come close to eliminating the gap. While Carr et al. (2022) dubiously compared *Tyrannosaurus* bones to those of today’s birds, they dismissed the more taxonomically informative fossil patterns. That such extreme variations in skeletal strength have not been documented in any one dinosaur species is the opposite of supportive with the single species hypothesis. The last is true whether the single species is applied to the TT-zone as a whole, or the upper section specifically.

Taxonomic implications – Divergences in fossil tyrannosaurid skeletal proportions within the clade and other megatheropods, including those of the femur – they not being overturned by less taxonomically pertinent comparisons of femoral robustness variability in ancient tyrannosaurids to today’s birds -- constitute standard anatomical differences and changes that can be used to help discern evolutionary patterns and diagnose paleotaxa. The apparent evolution of *Tyrannosaurus* bone proportions from the ancestral to a derived condition is fully and most compatible with the speciation expected over the long evolutionary time span of the TT-zone.

### Individual, Ontogenetic and Dimorphic Causes Do Not Explain the Variations in *Tyrannosaurus*

Paul et al. (2022) carefully examined the alternative explanations for the condition of *Tyrannosaurus* and found that they all failed to explain the degree and pattern of the variation in proportions and teeth as well as speciation. The study agreed with Carr (2020) that growth was not the cause for reasons detailed in both papers, and it is not likely that histological analysis will change that conclusion when some *Tyrannosaurus* femora that are less than three quarters the length of the longest femur as more robust than the latter (Fig. 12B). Another reason differences in ontogeny and size cannot be a solution is because the dimensions of the largest specimens from the three basic levels and three species are very similar. All are large individuals with femurs of 1100 to 1350 mm, and the largest specimens in all three levels and species sport femora of 1320 to 1350 mm, with estimated masses that vary by only a few hundred kilograms (Fig. 1B,C,E). Sexual dimorphism was not the cause because the ratio between robusts and graciles is well off 50/50, and because only robusts have been documented to be present early in the TT-zone – Carr et al. (2022) did not integrate these important factors into their paper. The same issues pertain to the stratigraphic separation between morphotypes of the postorbital bosses. Also contrary to the dimorphism hypothesis is how divergent growth and maturation patterns observed by Jevnikar and Zanno (2021) do not line up with the morphs. If the changes over time resulted from genetic drift, then that is what often creates new sibling species. The variation in the genus is not random over time as it would have to be to be plausibly attributable to individuality, so the latter does not provide a compelling, positive, evolutionary explanation for the shifts. Why would individual variance result in a difference in femur robustness in one tyrannosaurid species that exceeds that seen in all previous members of the family combined? All the more when variation in basal *Tyrannosaurus* is in the tyrannosaurid norm, and the variation according to the data on hand only appears in the upper TT-zone, and is skewed away from the ancestral condition? Likewise, how do intraspecific wandering explain the differences in postorbital bosses that are exactly the type that evolve to minimize interspecific reproduction? Attempts to use individual variation to explain the highly peculiar observed pattern will be ad-hoc opinion without scientific value.

A version of the individuality hypothesis for *Tyrannosaurus* variability proposes that the long lifespan of the dinosaur combined with its exceptional final size is responsible for inconsistent anatomy in a genus that underwent lots of transformation as it matured (Witton 2022). The three decades that *Tyrannosaurus* could achieve (Erickson et al. 2004, Hutchinson et al. 2011; Carr 2020; Cullen 2020) is short by the standards of mammals of similar size (Nowak 1991). Other giant theropods including tyrannosaurids had similarly long lives (Erickson et al. 2011; Cullen 2020). Growing from a few kilograms to elephantine mass in a taxon that may have exhibited little or no parental care can explain the tremendous variation in form with ontogeny – not unknown in other species – and may or may not show that some or all small specimens are juveniles of the large. But it is not clear how this offers a compelling evolutionary explanation for the variation observed in the nearly and entirely mature members of a species known from a limited geographic area over which the variations are laterally uniform – that is a reason Paul et al. (2022) and herein limited species determination to nonjuveniles -- the changes having occurred over a considerable span of time in a pattern that smells of Darwinian speciation. Long lives offer nothing to explain the postorbital boss diversity pattern.

Taxonomic implications. The evolution of variation in *Tyrannosaurus* dimensions in the crania teeth included and the postcrania away from the long standing tyrannosaurid ancestral conditions quickly to a derived status unique among tyrannosaurids is fully compatible with, and can only readily be logically explained by, selective genetics driven speciation.

### The Variation Factor

The taxonomic story of *Tyrannosaurus* that had not been fully appreciated and deeply examined is the exceptional degree of variation in the genus, and all the more its change over time. In terms display bosses, skull and skeletal robustness, and incisiform teeth. Such extensive variation has not been observed in other theropod species, and is not coherently explicable as the result of individual, ontogenetic, or gender difference within a species, all the more so because there appears to be a strong stratigraphic segregation between important aspects of the variability. The exceptional variation in *Tyrannosaurus* via-a-vis earlier tyrannosaurids is probably not the result of a large sample size because it is smaller than for the other tyrannosaurids, and because the outliers in gracility on the one hand (NHMAD “S”) and robustness (BHI 6248) were discovered fairly early (Larson, N. 2008) when the sample size of the genus was markedly smaller than it is now.

Taxonomic Implications -- No attempt to render the placement of all specimens in one species the superior hypothesis can succeed unless it is convincingly explained how two incisiforms and spindle bosses are known from the low TT-zone only, and graciles and knob or hat bosses only from high in the zone, and why so much quantitative inconsistency in so many regards is not observed in other tyrannosaurid species.

### *Tyrannosaurus* Floaters

Carr et al. (2022) are highly and for reasons discussed earlier problematically critical of the inability of the *Tyrannosaurus* species diagnoses of Paul et al. (2022) to place a substantial number of specimens in a specific species. That the species of the tyrannosaurid genus of concern are siblings very similar in most attributes heightens the probability that diagnostically insufficient specimens will exist. The placement herein of all but one major specimen is a named species undermines the Carr et al (2022) criticisms. The remaining unidentified specimens lack critical diagnostic features, and those in the upper layers are further conflicted by the apparent existence of two species. Lack of ability to vertically position a number of specimens is a hindrance that future stratigraphic work can be expected to alleviate but probably not eliminate. Small specimens are also often particularly difficult to place due to lack of adult characters, or potential status outside the genus.

Taxonomic implications – This and the proceeding Paul et al. (2022) are best efforts to determine and diagnose the *Tyrannosaurus* species that the evidence indicates existed, not to make one taxon so broadly defined that it conveniently but probably inaccurately accommodates all specimens on hand.

### Derived Tyrannosaurine Diagnoses

The statement by Carr et al. (2022) that “*Tyrannosaurus rex* can be distinguished from its *sister* [italics added] species *T. bataar*” is incorrect in that they are not that closely related. Being substantially separated by time, geographic distance, and anatomy, there had to have been a number of anatomically gradistic intervening species between them even if the two genera shared a common direct ancestor distinct from earlier large western hemisphere tyrannosaurids, and yet more interceding species if the American *Tyrannosaurus* descended from earlier tyrannosaurids on that continent rather than from Asian examples. Because they do not consider the situation at the genuine species level Carr et al. (2022) are actually diagnosing genera, or at least subgenera. In the process of doing so they fail to capture the complexities of the taxonomic data. In particular, the greater anatomical diversity contained exhibited by *Tyrannosaurus* vis-à-vis its more uniform Asian relation. The Carr et al. (2022) effort is also obsolete in lacking any consideration of the postorbital bosses, which their forcing of all *Tyrannosaurus* specimens into one species renders it impossible to define the visually catching divergences in the species display structures. Also out dated is their failure to take the stratigraphy into account.

The diagnoses herein are for the two most derived tyrannosaurine genera – although it remains possible that *Tarbosaurus* is a subgenus – and all the named species of both. Doing so better reflects the complexities of the anatomy and taxonomy. The characterizations are based on large specimens, and are collected from Paul et al. (2022), Carr et al. (2022), and those produced above. All pertinent characters are detailed, rather than some being diagnosed collectively as in Paul et al. (2022), partly in order to more explicitly describe the differences while avoiding the character undercount alleged by Carr et al. (2022). Some of the characters are tweaked vis-à-vis Paul et al. (2022), but there are not major alterations. These diagnoses include the postorbital bosses, Paul et al. (2022) being obsolete in that regard. Not utilized at the genus level are size differences because the contents of genera often vary greatly in dimensions, as per *Varanus, Canis, Panthera, Homo, Balaenoptera*. There is no observed size difference in the species of *Tyrannosaurus* (Paul et al 2022). Character overlaps and caveats are allowed as per *Triceratops* species diagnoses based on the Scannella et al. (2014) results, and for that matter in Carr et al. (2022) when it comes to tooth counts, as well as studies cited in the paleospecies determinations section. The stratigraphic factor is fully utilized.

Being exemplars of species determination, the postorbital bosses play in major role in this first broad diagnosis of the greatest tyrannosaurid genera and species. Large *Tarbosaurus* possess a rather subtle subcircular knob that does not project significantly above the top rim of the skull (Maleev 1955, 1974; Hurum and Sabath, 2003; assorted online photographs). It is most similar to those of *Daspletosauru*s, both *D. torosus* and *D. horneri* (Russell [1970]; Carr et al. [2017]; assorted online photographs). Due to the lack of diversity in these structures all specimens are readily accommodated in one species on this basis. The bosses of *Tyrannosaurus* are so radically divergent that it is not possible to produce a single, short description that accommodates all the variants, they range from horizontally elongated, low spindles to short but tall knob discs to less vertically prominent hat shapes. A situation entirely different to the uniformity in *Daspletosaurus* with its at least two species and *Tarbosaurus* with its current one, the only way to generate species pertinent diagnostic descriptions for the *Tyrannosaurus* variants in display devices is to place them in distinct species diagnoses, specifically three. Those who disagree with the multispecies diagnoses for *Tyrannosaurus* are challenged to produce a monospecific diagnosis for *T. rex* that incorporates all the differences in supraorbital bosses, as well as those in element robustness versus gracility.

Taxonomic implications – Carr et al. (2022) are correct that diagnoses are an important part of determining and defining paleospecies, but doing so for *Tyrannosaurus* using the full anatomical and stratigraphic data set produces different results from their more limited comparison of genera.

### What *Tyrannosaurus* Species Bosses and Diagnoses Reveal About Its Ancestry

The diagnosis of basal *Tyrannosaurus* in the form of *T. imperator* with a spindle boss is of phylogenetic importance because it is very unlike those of earlier and more basal tyrannosaurines *T. bataar* and *Daspletosaurus*. That indicates early *Tyrannosaurus* were not retaining an ancestral condition at least vis-a-via those genera in this particular regard. It is therefore possible that the ancestry of the TT-zone genus lies elsewhere. The *T. imperator* boss is more like that of the albertosaurines (Fig. 6), but is far from identical. Also of interest is that the *Tyrannosaurus* boss most like the earlier tyrannosaurines is the late *T. rex*, although its extra prominent boss is more derived and may have evolved independently. Or it represents the last of an unknown population of *Tyrannosaurus* with the old boss that moved into the TT-zone shortly before the K/Pg boundary, and its boss became more prominent to better distinguish itself from the more gracile *T. regina*.

Taxonomic implications – The *T. rex* only hypothesis has failed to notice the absence of the ancestral condition of the primary bony species specific ornamentation in basal American *Tyrannosaurus* because it inherently pays little attention to such variations in the genus. The multiple species thesis does pay very close attention to those critical items, and therefore advances the analysis of its phylogeny relative to the rest of the tyrannosaurs.

### Multiple Species is the Null Hypothesis for the Tyrant King

If *Tyrannosaurus* was known only from a shallow set of sediments spanning just a couple hundred thousand years, with little in the way of evidence of speciation of other dinosaurs in the same deposits, and if multiple taxa of large predators dinosaurian and otherwise dwelling in the same ecospace were rare or absent, then monospecificity would be the null hypothesis. As it is, the titanic predator is known from sediments that span at least half a million and more likely up to or over 1.5 million years (refs. in Paul et al. 2022, Mallon et al. 2022), with strong evidence for speciation in contemporary ceratopsids, some evidence for in pachycephalosaurs, and perhaps in hadrosaurs (Scannella et al. 2014, Fowler 2017, Paul et al. 2022; Carr et al. 2022). And two or a host of big theropods living in the same habitat is frequent in the Mesozoic. In this situation convincingly demonstrating monospecificity would require showing that the amount of cranial and postcranial variation in the giant TT-zone tyrannosaurid genus is low and randomly distributed. Instead, the scale of the variation and its distribution in a pattern that indicates evolution away from the ancestral condition refutes individual or dimorphic variation within a species, and is in much better accord with speciation toward more derived conditions that recap the contemporary robust and gracile species observed in earlier tyrannosaurids.

Carr et al. (2022) argue that the geological longevity of more basal tyrannosaurid species indicates the same could have been true of *T. rex*. This postulate is weak because it is not certain those species really were just one taxon each (as noted by Paul et al. 2022: Carr et al. 2022) in part because of limited sample sizes. Particularly pertinent is the possibility that *D. torosus* contains a cryptic sympatric species (Carabajal et al. 2021), a pertinent item Carr et al. (2022) do not mention.

And the geographic and evolutionary situation in the middle and late Maastrichtian of Laramidia was not the same as it had been in the period of relative stasis over the ten million years of rhino sized tyrannosaurids in the Campanian and early Maastrichtian, when the interior seaway isolated the western dinosaur populations from the bulk of the continent, limiting their size. The reunification of North America into one continent produced a general size increase to elephantine size in the region’s dinosaurs that dramatically increased their rate of evolution. Which appears to have been continuing in the late Maastrichtian tyrannosaurids as *Tyrannosaurus* experienced a split into the older robust and a new gracile form, replicating earlier patterns of two such tyrannosaurid morphotypes sharing the same habitat (Paul et al. 2022). Carr et al. (2022) neglect to discuss the significance of this exceptional evolutionary potential of this pre K/Pg speciation factor.

Taxonomic implications -- Multiple species is the null hypothesis unless the fundamental data concerning *Tyrannosaurus* is seriously challenged by examination of the status of the specimens utilized in Paul et al. (2022) and this analysis. The simplistic argument by Carr et al. (2022) that a monospecific *Tyrannosaurus* is the null hypothesis is correspondingly biased towards that theory.

### One, Two, Three or More Species

In view of the long time span over which *Tyrannosaurus* lived during which some other dinosaur genera underwent speciation, and the observed changes in anatomy, the question is less likely to be whether *Tyrannosaurus* was multispecific, but how many species are represented by the TT-zone fossils. The shift in incisiform tooth count and the expansion in proportional variation with the advent of gracility strongly indicates at least two species. As explained in Paul et al. (2022) if the upper TT-zone specimens are one taxon then the onset of the expanded variation relative to early tyrannosaurids *T. imperator* included is the evidence of the novel reproductive shift that would mark a new chronospecies. But that hypothesis is inferior to two late species because such strong dimorphism had not been seen in prior tyrannosaurid species, *T. imperator* included, by a factor of two. If instead the upper TT-zone *Tyrannosaurus* remained all robust that too would favor chronospecies, as would all high placed specimens being gracile, and a lack of major variation in postorbital boss form in the very last *Tyrannosaurus* would indicate chronospecies. That the evidence instead indicates there is atypically high variation in high TT-zone *Tyrannosaurus* is most compatible with separation into two taxa of robust and gracile form as had been observed in earlier tyrannosaurids inhabiting the same ecospace. That is even more probable because that variation is entirely due to the swift shift to gracility away from what had been the long lasting tyrannosaurid norm, impacting even the bar separating the preorbital fenestrae, which best fits the adaptative speciation model. Also consider that if heavily built *T. imperator* is valid, then the separation from that is greater in distance vis-a-vis gracile *T. regina* than it is compared to stouter *T. rex*, so *T. regina* is an anatomical divergence driven species. Aside from the femur the maxilla (in both overall dimensions and the pillar dimensions) produces the strongest bone results in support of 3 species. To that add that the varying configurations of the postorbital bosses is most in line with sexual dimorphism within three species.

In order to test two chronospecies versus three species *the T. imperator* and *T. rex* were diagnosed, with *T. regina* arbitrarily subsumed into the contemporary *T. rex* in the systematics section. The result was the dramatic reduction of the diagnostic characters down to the anterior dentary teeth and the orbital bosses. The separation between the two species remains fully valid, just one character being sufficient, and the species grade display structures being especially definitive. But all the many differentials between the species regarding robustness and display features, and the exceptional anatomical and statistical variation in the high TT-zone fossils compared to the much more uniform *T. imperator* including the unprecedented shift to gracility, is disappeared without logical scientific justification. And the sharp reduction of the character list is contrary to those who favor large numbers of characters separating species. Statistically awkward is that only two of the many *T*. “*rex*” skulls have the knob supraorbital bosses that help diagnose the species, when many of the *T. imperators* have the spindles that characterize that taxon. So two chronospecies, although well superior to *T. rex* alone, is markedly inferior to three species characterized by a host of features. This exercise reinforces the need for all studies that designate paleospecies to incorporate species systematic diagnoses to help test the favored hypothesis.

Taxonomic implications – With ontogeny, random individuality and dimorphism falling short in explaining the changing circumstances of the giant theropod progressing from the lower TT-zone to high in the formations, a single species is scientifically inferior to two, and two is inferior to three. AMNH 5027 may hint at yet more.

### Famed *Tyrannosaurus rex* Does Not Deserve Special Scientific Deference, Protection and Effort

In the submitted version of Paul et al. (2022) it was repeatedly noted that *Tyrannosaurus* is just another dinosaur and it should be treated like such, and our results not be subjected to greater or lesser critical analysis. Reviewers required that those comments be cut back, which proved to be a mistake.

Taxonomic implications -- To repeat, *Tyrannosaurus* is just another dinosaur, and the status of the species it contains must be dealt with like those of other extinct tetrapods with no special consideration. The notion that any fossils deserve special scientific attention should never have been proposed.

### Anagenesis or Cladogenesis?

If the only upper TT-zone species is *T. rex*, then it is a candidate for direct descent from *T. imperator*, but anagenesis is not certain, all the more so because the postorbital display bosses of the two species were so divergent. If there are two upper TT-zone species, then at least some degree of cladogenesis must have occurred. That neither *T. rex* nor *T. regina* bore the distinctive spindle postorbital bosses of *T. imperator* further complicates the difficulty of determining which if either of the first two species directly descended from the latter (Paul et al. 2022), and again increases the possibility that anagenesis was not involved. So does how while the orbital boss of *T. regina* was somewhat more similar to that of *T. imperator* than was that of *T. rex*, the latter better retained the robust construction of the tyrant lizard emperor. Yet another complication is that none of the TT-zone species retains postorbital bosses like those of *Tarbosaurus* and *Daspletosauru*s.

That increases the possibility of cladogenesis of the later species relative to *T. imperator*. Figure 1b in Carr et al. (2022) is therefore simplistic in that is represents only the anagenesis from *T. imperator* to *T. rex* with cladogenesis for *T. imperator* hypothesis, the alternatives are not included. And *T. imperator* probably evolved before the advent of the zone, while substantial populations of *T. rex* and *T. regina* were probably liquidated by the K/Pg crisis. Also possible are intermediate populations, perhaps in the mid TT-zone, of which AMNH 5027 may be a representative.

Taxonomic implications – Determining the mode of evolution within TT-zone *Tyrannosaurus* is not workable with the current data, and this impediment may never change.

### *T. imperator* and *T. regina* are not junior synonyms of juveniles of the species *N. lancensis* and *G. megagracilis*

This is an item that does not challenge the existence of the two new species, but of their names.

Paul et al. (2022) note that the holotype of the species from the lower TT-zone is not, as has been suggested, the juvenile *Nanotyrannus lancensis* (Bakker et al. 1988). It is it a nomen dubium – only a distorted skull too immature to possess the diagnostic supraorbital boss or skull proportions, sans the diagnostic appendicular material, or that or other postcrania to examine and compare its growth pattern, and too juvenile to assess whether it would grow up to be a robust or gracile in any case. Adding to the problem is the strong possibility that at least some small TT-zone tyrannosaurids did not grow up to be giant *Tyrannosaurus* – the sharing of a habitat by a gigantic and a much smaller, far more gracile taxa has the precedent of *Tarbosaurus* and *Alioramus* (Brusatte et al. 2012). The arm of NCMNS “Bloody Mary” from low in the TT-zone is longer than the femur which is not observed in any other Campanian/Maastrichtian tyrannosaur/ids except *Dryptosaurus* (Paul 1988; Brusatte et al. 2011) from earlier east coast sediments (condition not known in *Appalachiosaurus*), the forelimb is always being shorter including in juveniles of Asian and western North American tyrannosaurids. And the NCMNS “BM” hand is literally as long of that as its supposed grownup *T. imperator* holotype, while the similar hand of another small tyrannosaurid is even longer (Fig. 13; Larson 2013). That does not happen in ontogeny. It is potentially pertinent that the growth curve of BMRP 2006.4.4 does not appear to be in accord with that of *Tyrannosaurus* (Jevnikar and Zanno 2021), it will be interesting to see the results for NCMNS “BM”. *N. lancensis* is not even a good holotype for its own taxon if it is not a young *T. rex* because it is nondiagnostic, much less for any adult *Tyrannosaurus* species.

**Fig. 13.**
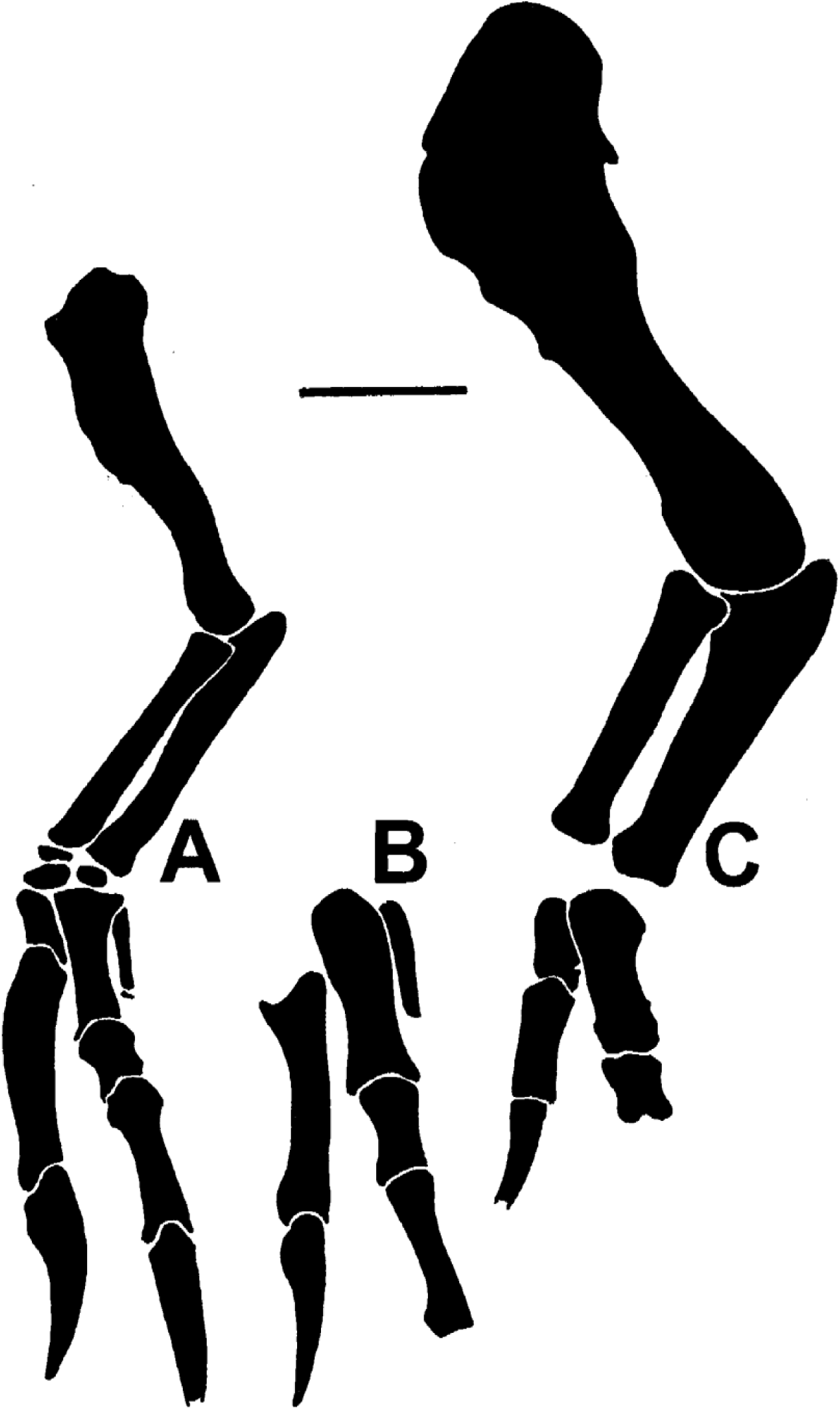
Preserved elements of tyrannosaurid forelimbs from lower TT-zone to same scale, bar equals 100 mm. **A** NCMNS “BM”. **B** “Jodi”. **C** *T. imperator* holotype FMNH PR208, placement of distal elements not certain for this specimen.

The holotype of *A. megagracilis* (Paul 1988) is from high in the TT-zone, very probably is a *Tyrannosaurus*, and may belong to *T. regina* as suggested in Paul et al. (2022). But like the *N. lancensis* holotype this fossil also is too juvenile and incomplete to be used as the holotype of the gracile species. The individual is too small to apply to it the critical 2.4 femur dimensional ratio. Although the incomplete femur looks like it was gracile by juvenile standards, that is an estimate not a measurement, so the actual ratio is not known, and it is possible that some *T. rex* juveniles of this size had the same femur ratio. Not available for assessment is a mature postorbital boss. Placement of this holotype in *T. regina* is therefore automatically tentative -- to the point is may be a *T.* incertae sedis -- too much so to be taxonomically significant. *A. megagracilis* is another nomen dubium.

A cautionary taxonomic tale. the first described remains that unambiguously belong to *Tyrannosaurus* and – if the genus is considered monospecific – to *T. rex*, is the very fragmentary AMNH 3982 that is the holotype of *Manospondylus gigas* (Cope 1892). The fossil may be from the lower TT-zone, possibly being part of BHI 6248 (Larson, N. 2008) which scores as a robust *T. imperator* (Paul et al. 2022). The specimen was logically dismissed as a nomen dubium by Osborn (2017). Because it can be referred to *Tyrannosaurus, Manospondylus* would be the proper name for the genus if being referable to a taxon alone means the name of the holotype of the earlier named genus takes precedence regardless of its diagnostic nonvalue. Also overturned is *T. rex* if the genus is monospecific, or *T. imperator*/*T. lancensis* if the later low placed species is valid. Ergo, if *T. lancensis* must be the name that takes precedence over *T. imperator*, and *T. megagracilis* over *T. regina*, then the same taxonomic il/logic forces *Tyrannosaurus* to be subsumed into *Manospondylus* – note that *M. rex* and *M. regina/megagracilis* survive if the multispecies hypothesis holds. The moral of this systematic tale is that a paleogenus and its species need to be founded on sufficiently diagnostic types.

Taxonomic implications -- Had Paul et al. (2022) used the *N. lancensis* skull as the foundation for the species of the lower TT-zone *Tyrannosaurus* (*T. imperator*) instead of big and highly complete FMNH PR2081 it would have been severely and correctly criticized. Same if Paul et al. had it tried to use the immature and fragmentary *A. megagracilis* holotype in place of the far better adult USNM 555000 for the higher gracile TT-zone *Tyrannosaurus* (*T. regina*). Opposition would have approached that had Paul et al. (20220 tried to sink *Tyrannosaurus* and *T. rex* into *Manospondylus* and *M. gigas*.

### Evidence for Multiple North American *Tyrannosaurus* Sibling Species – the Summary

**For at least two species --**

Studies prior to Paul et al. 2022 had not thoroughly tested much less strongly verified the monospecific status of the wastebasket taxon *T. rex*.

Multiple species are the norm within a genus.

Sufficient geotime was available for sibling level speciation and may favor such, in parallel to speciation observed in *Triceratops* over exactly the same stratigraphic span.

Radical alterations in regional geography in Maastrichtian as North American continent reunited and dramatically expanded resource base, probably favored a burst of rapid evolution in tyrannosaurids that could favor more rapid speciation than observed in earlier tyrannosaurids.

Paleospecies can be and are regularly designated based on the sample that is available, it does not require an ideal large statistical data set which if it becomes available can then be used to test the hypothesis – perfection is the enemy of good enough.

The cumulative data in support of multiple species of *Tyrannosaurus* is broadly comparable for *Triceratops*, in some regards better (the tyrannosaurid work is more holistic in that significant patterns are observed in both the crania and postcrania, not just the rostrum as in the ceratopsid), and superior to other intrageneric dinosaur species in terms of sample size, stratigraphic changes over time, and other factors.

Specimen sample size analysis is based upon is larger than usual for nonavian dinosaur genera which can be as few as two, and is comparable to that for *Triceratop*s.

Number of characters utilized to diagnose species, 13, is well within norms for identifying fossil sibling species which can be as few as one.

Stratigraphic data is adequate for purposes, and comparable or superior to that for most paleospecies level taxonomic work.

There is stratigraphic separation between distinctive postorbital boss visual displays – horizontally elongated spindles only low in the TT-zone, anteriorly placed, elevated knobs only high in the formations -- that are in full accord with, and a leading identifier of species. The monospecies hypothesis lacks a viable explanation via ontogenetic, individual or dimorphic inconsistency for the variable and sometimes time separated bosses, which alone firmly establish that the genus was multispecific.

The variation in bone based species grade features as represented by the postorbital bosses is higher in *Tyrannosaurus* than is usual for predators, including other tyrannosaurid species, and is comparable to that present in some herbivores; cranial ornamentation.

The general consistency between right and left bosses – there are no specimens with a spindle on one side and the disc knob on the other – confirms that they were genetically programmed, adaptative emergent structures that evolved among differing species.

Much more variation in fossil femoral robustness than observed in any other theropod or dinosaur species – the variation being significantly statistically greater than a sample of femora of an allosaur species from a single quarry -- including any tyrannosaurid species including two that dwelled in the same formation, and statistically well above that yet observed in all other tyrannosaurids combined consisting of up to 7 genera and 8+ species from two continents spanning 10 million years compared to 0.7-1.5+ million years for a smaller *Tyrannosaurus* sample from a small region, strongly favoring speciation over sexual dimorphism, ontogeny, or individual variation.

That total number of robust femora are over twice that of all gracile femora strongly contradicts both sexual dimorphism and ontogeny as causes.

That growth and maturation curves do not match up with robustness contradicts dimorphism as a cause.

Some femora that are only two thirds adult size are robust, in some cases more so than some of the longest femora, while that some of the largest are gracile with the longest known femur being the slender-most among adults, directly contradicting ontogeny as the cause of high robustness.

That reproduction has not been shown to have been occurring as early in ontogeny as the onset of large variations in the robustness of juveniles means that early reproduction does not currently offer an explanation for the observed pattern.

That the solely robust femora of early *Tyrannosaurus* followed by the much greater variation in proportions higher up include substantial gracility is due to a relatively smaller earlier sample is not the most likely scenario, because while the variation in a smaller sample may be less than in a larger sample, is not likely to be skewed one way or the other relative to the latter

The most robust femora from the upper TT-zone not being as stout as the most robust examples from low in the zone further supports the pattern being real, and is in accord with a proportional shift in the genus, rather than the stasis most compatible with no speciation.

Because low variability limited to robustness in early *Tyrannosaurus* appears to be a retention of the ancestral condition observed in other earlier tyrannosaurids (both individual species or in total) additionally supports the limitation to only robustness among basal *Tyrannosaurus* as probably being real.

That gracile femora are found only in upper TT-zone, while robusts are present in all levels, contradicts the consistent strong proportional variation necessary for dimorphism to be persistently present.

That proportional variation in low TT-zone *Tyrannosaurus* is not higher than observed in other tyrannosaurid species is compatible with and indicates that only one species was extant at that level.

Proportional variation being low in the lower TT-zone and remarkably high in the higher TT-zone strongly indicates speciation either because the sudden onset of major dimorphism indicates the kind of dramatic shift in reproductive behavior that is the epitome of species separation and designation, or two new contemporary species with each retaining the limited dimorphism apparently typical of dinosaurs.

The same basics as immediately above apply if the doubling of proportional variation in the upper TT-zone was primarily due to a new ontogenetic pattern or individual variation both of which are improbable, but in any case indicate a change radical enough to require recognition of species differentiation.

The visually dramatic shift in supraorbital display structures in parallel to the great increase in proportional variation combines to produce overwhelming evidence of the shift in reproductive behavior indicative of species differentiation.

In 7 characters there is in increase in variation within a given element from modest to many fold. In none is there a decrease in variation.

The persistently robust *Tyrannosaurus* sample from the lower TT-zone is smaller than the more gracile set from higher levels, but the sample size difference should not result in the strong skew. So as the lower sample increases in abundance it is not likely that gracile specimens will prove to be as proportionally numerous as they are higher up if they appear at low levels at all.

If the smaller sample of lower TT-zone femora greatly expands to include much more gracility than in other tyrannosaurids with future finds, then the great proportional variation compared to other theropods is most in accord with the presence of two species early in the evolution of the genus. If a future lower set shows that graciles are a present but rare compared robusts then the case for two species at that level will be at least as strong, or more so.

That gracile *Tyrannosaurus* femora are unusually slender by normal tyrannosaurid standards when allometry is accounted for, and represent a highly atypical shift over a short period of time, directly contradicts ontogeny while strongly favoring subtle adaptative evolution via speciation.

10 other measurements of robustness in crania and postcrania favor actuality of robust and gracile morphs in generally good accord with femoral robustness.

In 7.5 items the most robust or two incisor tooth condition is observed in basal *T. imperator*.

1. *T. imperator* is never the most gracile in any element or ratio – such proportional extremity has not been observed in *Triceratops*.

In only 2 ratios is a *T. regina* the most robust overall, in two cases the sample is on the small side, especially for *T. imperator*.

In 9 ratios the most gracile condition is observed in derived *T. regina*.

In 5 elements all *T. regina* are more gracile than any *T. imperator*.

In all 11 elements and 12 ratios there is an overall trend, form minor to strong, toward greater gracility progressing geologically upwards. Trends towards increasing robusticity have not been discovered.

There are 10 cases of nonoverlapping bimodal separation between the species – the extent of non/bimodality is broadly similar to that observed in many other tetrapod intragenera paleospecies including *Triceratops*.

General shift from the probable ancestral condition of 2 small functional anteriormost dentary incisiform teeth to just one progressing or less upwards in TT-zone, is an adaptative trend not actively explained by dimorphism, individual variation, or ontogeny.

“Long” life span and related issues do not provide strong explanation for the observed patterns.

Progressive change in functional dentary incisor number over time correlates with changing robustness femoral and otherwise, plus the dramatic alteration in supraorbital boss shapes, all occurring in synch, strongly accords with evolutionary speciation rather than dimorphism, ontogeny, or mere individual variation.

The unique shapes of *Tyrannosaurus* postorbital bosses, and the exceptional gracility of *T. regina*, are species level autapomorphies.

The preponderance of the cumulative evidence including sample size and other parameters that is the norm in modern paleospecies work now overwhelmingly favors speciation over all alternatives, and is stronger than average for other multiple sibling species in dinosaur genera, being close to that documented for contemporary *Triceratops*, including the species definitive diversity in species category display features. Multiple species of *Tyrannosaurus* is at this time easily the best documented hypothesis over the alternatives.

### For three species rather than only two chronospecies

Three species are readily diagnosable with each being about as distinguishable from the other two as are the others, diagnosing just two chronospecies greatly reduces the number of characters distinguishing the remaining taxa without justification, fails to take into account the tremendous level of variation within and between the upper TT-zone specimens, and that is the opposite of taxonomic logic.

Much more observed variation in fossil femoral robustness high in the TT-zone than observed in any other theropod or dinosaur species, and in all other tyrannosaurids combined, and twice that observed in earlier *T. imperator*, strongly favors lateral speciation in addition to and over just vertical chronospeciation.

No prior tyrannosaurid or dinosaur species has been shown to have been sexually dimorphic to the degree seen in late TT-zone *Tyrannosaurus*.

Two taxa of earlier western North American giant tyrannosaurids, with one more robust than the other, are present in the same levels of the same formation.

Strong divergence in postorbital boss visual displays, including the apparently male knob versus hat shapes, in upper TT-zone is a leading indicator of species. The chronospecies hypothesis lacks a similarly viable explanation for the very different bosses of *T. rex* and *T. regina*.

The number of upper TT-zone skulls with a prominent knob shaped postorbital boss is half that expected if they represent one gender of one species.

Two high TT-zone species is in best accord with the hypothesis that expansion of the resource base was a driving factor in the combination of both vertical and lateral speciation of elephant sized giant predators as the latest Maastrichtian progressed.

In 5 elements all *T. regina* are more gracile than any *T. rex* among high level specimens.

In all 6 cranial robusticity plots the most gracile ratio is that of a *T. regina* – such proportional cranial extremity has not been observed in *Triceratops*.

In no element is *T. regina* always more robust than is its contemporary taxon.

That *T. regina* cranial elements always include the most gracile or all gracile examples as well as the femur, indicates it underwent an extensive evolutionary overhaul involving *reducing* the strength of major and minor skull and skeletal bones in a genus known for its immense size and massive construction, a species level development that was highly divergent from more basal and traditionally robust *T. imperator*, rather than just a side variant of *T. rex*.

The three sibling species hypothesis offers the ability to provisionally determine the sexual dimorphism patterns in the species, while adding to the cumulative evidence that 1. *T. rex* and *T. regina* are not conspecific.

### Assessment of Carr et al. (2022)

Sets higher standards for determining, diagnosing, and naming sibling paleospecies than is the norm in vertebrate paleozoology.

Assumption that monospecifity of *Tyrannosaurus* is null hypothesis problematic because genus existed over a sufficient time span for speciation to have occurred. As a result quickly produced Carr et al. (2022) is a narrow and negative examination of Paul et al. (2022), rather than a positive expansive examination and testing of number of species in genus.

Does not cite evidence that other tyrannosaurid species that appear to be long lasting may be taxonomic chimeras.

Claims that Carr (2020) demonstrated one species when that work assumed such, was not designed to test the specie question, and used inferior anatomical and stratigraphic data bases for determination of paleospecies.

Does not document that any stratigraphic data in Paul et al. (2022) is errant, or note that some of it is in accord with Carr (2020).

Does not demonstrate that the quality of the stratigraphic data in Paul et al. (2022) is not adequate for determination of paleospecies over a span of many hundreds of thousands of years.

Criticism of the Paul et al. (2022) sample size is excessive because it is larger than for most other noncontroversial dinosaur paleospecies, is markedly larger than used by Carr (2020), and approaches that of Scannella et al. (2014) *Triceratops* sample that Carr et al. (2022) cite positively.

Strong criticism of Paul et al. (2022) for using private specimens on practical grounds including data replication, as well as ethical issues, is itself ethically and scientifically problematic. Excluding the data severely hinders testing the species problem by seriously reducing the sample size and eliminating some of the extreme data ends. And Carr et al. (2022) themselves use the Paul et al. data without alteration or deletion of the privately held remains to test Paul et al.’s results.

Does not consider how dramatic geographic changes underway in late Maastrichtian North America accelerated evolutionary rates in the dinosaurs of that time and place.

Does not refute any Paul et al. (2022) robustness measurements, utilizes some of them as per immediate above.

Comparison of proportions of ground striding, free living, fossil megapredator’s thick walled femurs to small, flying, sometimes semiaquatic extant archosaur’s thin walled femurs is problematic. In part because Paul et al. (2022) conduct the paleozoological research norm of comparing and tracking differences and changes among fossils in variations and directions in robustness, especially between last *Tyrannosaurus* to more basal examples of genus, and to all earlier tyrannosaurids combined, which Carr et al. (2022) do not consider.

Does not consider the sudden exceptional increase in proportional variation and the sudden trend towards gracility in the last *Tyrannosaurus* vis-a-via early members of the fossil genus and earlier tyrannosaurids, including the variation if femoral proportions being much greater than all tyrannosaurids combined.

Does not consider that *Tyrannosaurus* robusts are about twice as common as graciles.

Does not refute major shift from two small incisiform anterior dentary teeth in nearly all lower TT-zone specimens to no examples in later specimens, does suggest possible ontogenetic development behind the adaptative evolutionary functional trend.

Criticism of lack of nonoverlapping bimodality in data sets is excessive because such is not critically necessary is assaying sibling paleospecies, and because not all the data used by Carr et al. (2022) to diagnose *T. rex* is intraspecies consistent and interspecies entirely distinctive.

Diagnosis of *T. rex* is simplistic actually being of genus *Tyrannosaurus* vis-à-vis *T. bataar*, does not attempt to address complexities of stratigraphy or directly text species diagnoses.

Does not examine cranial display ornamentation.

The contents of Carr et al. (2022) does not overcome the strong preponderance of evidence for a multispecific *Tyrannosaurus*.

### What Needs to Be Done to Firmly Refute More Than One *Tyrannosaurus* Species

Do not cite papers prior to Paul et al. (2022) as having had established the monospecificity of *T. rex*, unless it can be shown that any of them included a large sample of dozens of stratigraphically correlated specimens that did not show a pattern of change over time.

Because monospecificity is not the null hypothesis either in general, or for *Tyrannosaurus* because the evidence for multispecies meets current paleozoological standards, and the popular status of *T. rex* must be ignored in scientific analysis, it needs to be convincingly shown that the preponderance of evidence favors one species over more than one sibling species.

That requires a large sample of dozens of stratigraphically correlated specimens that do not show a pattern of change over time.

It follows that the maximum available data sample must be used as per Paul et al. (2022). If the sample is smaller, then any results are automatically less or none definitive, and may be misleading.

The second need also requires showing that there is not an exceptional level of variation in *Tyrannosaurus* compared to other tyrannosaurids.

The above in turn further requires at least in part demonstrating that there are significant errors in the proportional measurements in Paul et al. (2022) and herein that when altered favor monospecificity.

And/or positively demonstrate that there are significant errors in the stratigraphic placement in a number of specimens in Paul et al. (2022).

In particular, firmly establish that a number of gracile specimens were located in the lower TT-zone. If that is not possible then the multispecies hypothesis is strongly supported over the alternatives.

Firmly establish that a number of specimens with two small incisiform dentary teeth are present high in the TT-zone. If that is not possible then the multispecies hypothesis is strongly supported over the alternatives.

Firmly establish that a number of specimens with knob shaped postorbital bosses were located in the lower TT-zone and spindle bosses in the upper TT-zone. If that is not possible then the multispecies hypothesis is strongly supported over the alternatives.

If the above cannot be done, present a plausible, natural selection based hypothesis that logically explains the selection of visually distinctive display organs of the type well suited for species determination and diagnosis, without the evolution of new species.

In the case that the variation over time patterns observed by Paul et al. largely hold, then if dimorphism, ontogeny or individual variation are proposed to explain the observed pattern it must be explained in detail how any or all are positively superior to the speciation hypothesis in the context of adaptative natural selection as per evolutionary theory; the “long” lifespan of gigantic *Tyrannosaurus* is a weak ad-hoc opinion for the reasons noted above. Studies that show that such patterns of variation within paleospecies need to be specifically cited and discussed.

In other words, actually show that there was just one species, not merely try to negate the evidence for more species, the latter hypothesis being at least as plausible as the former if not more so. For example, the current evidence currently favors the existence of only one species in the early TT-zone because of the overall low variability of the characters. It is the late TT-zone that contradicts one species both because of significant changes in characters from the basal tyrannosaurid and *Tyrannosaurus* conditions, and the uniquely wide variations both among between the latest specimens of the genus, away from the basal condition of the genus and its family. So show why the condition of the upper TT- zone *Tyrannosaurus i*s not indicative of speciation, when the circumstances are so radically different form the earlier *Tyrannosaurus* that more clearly fit into one species at that level. How for instance is the extreme variability in femoral proportions in large *Tyrannosaurus* compared to the rest of the genera and species in Tyrannosauridae clearly explainable without speciation in the one former genus?

A mass data base in number of characters has not been tested for efficacy, may not be replicable by other researchers, and a resulting small number of distinguishing characters meets norms for designating sibling paleospecies.

Do not treat Paul et al. and the *Tyrannosaurus* situation in isolation from the that seen in other modern works of dinosaur and other tetrapod paleospecies. Doing that leaves open the possibility of claiming the data does not support multiple species when it has not been shown that is true relative to the prevailing norms, and it fact the standards have been met. It is therefore necessary to directly demonstrate that Paul et al. (2022) and herein do not meet and been consistent with the methodological and procedural criteria widely used to establish and diagnose fossil sibling species of dinosaurs and other tetrapods based on skeletal remains, with citations and comparative analysis of a variety of studies including those cited by Paul et al (see the 2022 paper’s supplement). That includes citing studies that found similar patterns of variation over time while explicitly rejecting speciation as the cause. Or, demonstrate that the current standards are insufficient, need extensive reform, and detail how such should be done in the context of the current literature on the subject – and in view of the inability to even rigorously define what a species is whether living much less extinct -- while noting what other paleogeneric multispecies do not stand up to the elevated standards. Do not fail to do one or the other; treating *Tyrannosaurus* species determination outside the context of current norms is inconsistent and not scientific as it evades critical comparative analysis – i. e. it is giving the genus the special status that no genus should receive.

In particular, treat *Tyrannosaurus* as just another fossil predator that as usual for the trophic type possesses little in the way of species specific display structures, do not primarily compare the variation the genus contains to herbivores with prominent horns, antlers, crests and the like.

Assess whether the *Tyrannosaurus* data supports siblings species that have minimally diverged from one another, not species in large genera that have widely diverged from one another over many millions of years, or are in different genera.

If multiple species are accepted, but just two chronospecies (*T. rex* and *T. imperator*), then it needs to be directly explained how this hypothesis is superior to two high TT-zone species when such a large variation in robustness is not observed in other tyrannosaurid species or the rest of the family as a whole, while two taxa one robust and the other gracile are observed in other tyrannosaurids from the same paleohabitat, and the graciles are so extreme in their proportions.

Remember that *Tyrannosaurus,* and *T. rex*, is just another tetrapod, and that the nondivisibility of the type specifies requires no more or less defense than does *Brontosaurus excelsus, Triceratops horridus, Stenopterygius quadriscissus, Ornithomimus edmontonicus, Psittacosaurus mongoliensis, Metarhinus abotii*…..

The above list is not necessarily all that is required.

## CONCLUSIONS

Paul et al. (2022) and this examination are the first studies that found the following. That -- as demanded by some multispecies skeptics -- the three named species bore the diversity of cranial display features that are hallmarks of species determination, while in the process allowing provisional assignment of specimens to genders. That basal *Tyrannosaurus* retained some of the general ancestral tyrannosaurid conditions that had previously been in force for ∼10 million years. Starting with limited proportional variation, in particular with femoral proportions continuing to be persistently robust when allometry is accounted for. The anterior lower incisiforms numbering two. That with exceptional evolutionary rapidity – apparently in association of the general elevation of dinosaur evolution driven by the reunification of North America -- the proportional differentiation suddenly expanded in major elements of the crania and postcrania for the first time, manyfold in most cases, exceeding all previous tyrannosaurids combined in femoral variation. Dentary incisiforms dropped to an unprecedented just one as the new norm. Those very nonrandom events left the latest portion of the TT-zone inhabited by robust and gracile examples of giant tyrannosaurids, replicating the presence of a similar situation in earlier habitats in the same region. The sudden gracility of half of the most recent specimens was the opposite of the massiveness expected in the most gigantic tyrannosaurid, and included strength reduction of the most important tooth bearing elements and the vertical struts in a skull otherwise notable for its exceptional biting power, unmatched in any other terrestrial predators. The remarkable changes in robustness regarding both variation and more gracility, in anterior dentary tooth counts, and in display boss shapes all occurred in evolutionary synch with one another. Because these developments fly in the face of longstanding tyrannosaurid patterns, and/or are the opposite of the attributes predicted to characterize the most titanic genus of the clade, and are the changes expected between divergent species, they are dramatic evolutionary shifts that cannot without convincing warrant be dismissed as minor and intraspecific. Ergo *Tyrannosaurus* was multispecific, thrice so, based on current evidence.

Instead of noting that ground breaking nature of Paul et al. (2022) and its utility for study of the genus and its species, the general reaction by researchers most especially Carr et al. (2022) was to make a series of sometimes severe criticisms of the paper not seen before in paleozoology regarding other dinosaur species. This examination documents that the criticisms were often factually problematic at best both in regards to the issue of *Tyrannosaurus* taxonomy, and the determination of tetrapod paleospecies.

If the basic science supporting the evolutionary patterns observed by Paul et al. in 2022 and herein bear out over time, is it viable to attribute such a notable set of nonrandom events to individual variability which is inherently random, or to ontogenetic or dimorphic factors that do not fit the nature of the changes? The answer is a solid negative. Attempts to dismiss such a complex of atypical and nonrandom events by attributing them to either random individuality or ontogeny or dimorphic factors that do not fit the nature of the changes, are likely to prove uncompelling and arbitrary because they do not meet the basic criteria of a constructive cogent scientific hypothesis. Such intraspecific risk being ad-hoc, casual opinion that do not via a positive explanatory hypothesis actually explain the outstanding pattern, and consequently have a low possibility of being correct and informative. Such is not rigorous, evolution based coherent explanations for a pattern that is readily and fully explained by genetics driven speciation in response to selective factors. To synonymize *T. regina* and even more so *T. imperator* with *T. rex* will require a major set of positively affirming data, incorporating all the specimens.

Paul et al. (2022) proposed that the evolution of sibling species within *Tyrannosaurus* was an example of a level of evolution more subtle than seen in other dinosaurs including contemporary *Triceratops*. With the addition of the data of the visually vivid differences in the supraorbital bosses, the evolution of *Tyrannosauru*s species was not so subtle after all.

In addition to future examinations of possible sibling paleospecies being required to examine all potential causal explanations for variation including dimorphism, ontogeny and individuality as proposed by Paul et al. (2022 suppl.), another necessity should be a diagnoses of the species at that basic taxonomic level.

By utilizing data taken from privately owned specimens for their statistical advantage and necessity, Carr et al. (2022) ironically negated their criticism of the practice, and de facto have left others free to do the same in future work on *Tyrannosaurus* and other fossil vertebrates.

Taxonomic implications - In order for individual, ontogenetic, and dimorphic factors to explain the observed remarkable nonrandom events regarding *Tyrannosaurus*, they must offer logical reasons that one of them are behind the exceptional pattern. But they do not do so. Only Darwinian speciation presents a viable evolutionary explanation, indeed that hypothesis is forced by the evidence that is on hand. To put it another way, the data indicates that a population of the last of *Tyrannosaurus* consisting of half the specimens from the upper TT-zone, had adapted characteristics that suddenly verged well off from what had been the tyrannosaurid norm for the last quarter of the Late Cretaceous. The variance was so extensive that it even effected details such as strength of a maxilla bar. As two new species appeared their cranial ornamentation evolved to produce distinctive new and autapomorphic cranial display features. That this represents nothing more than minor random variation or drift within a species that remained just one over an extended time span that records speciation in a period of unusual evolutionary rapidity in other dinosaurs is not logical in scientific and evolutionary terms. It is a patent, classic example of speciation into new, anatomically and visually distinctive adaptative forms. The *Tyrannosaurus* multispecies hypothesis meets the current standards and norms for assessing and naming paleospecies. The evidence indicates that rather than being stuck in *T. rex* stasis, the last tyrannosaurid genus of the region was evolving new species as were its herbivorous dinosaur prey before events came to a very rapid termination.

## ACKNOWLEDGMENTS

Thanks go to Scott Persons, Jay Van Raalte, Donald Henderson, John Scannella, Gerald Mayr, Thomas Holtz, Niclas Olsson, Tanner Frank, Eugen Zudin, and those listed in Paul et al. (2022).

## APPENDIX 1

### Restoring the Differing Appearances of *Tyrannosaurus* Species

Until now there has been little effort to use scientific techniques to restore *Tyrannosaurus* in the context of multiple species, it in particular being assumed that the widely differing postorbital bosses were merely intraspecies variants including in Paul et al. (2022). That the exceptional bosses of *T. rex* as exemplified by UWBM 99000 and RSM 2523.8 (Fig 1B,G) were not available until recently has contributed to this failure to pay closer paleobiological and paleoartistic attention to the bosses. After the publication of Paul et al. (2022), Paul (2022b) indicated that there are no clear visual attributes that distinguished the species. In that article (and accompanying some cited news stories) *Tyrannosaurus* with orbital bosses shapes not yet found in lower TT-zone *T. imperator* were illustrated attacking *Triceratops horridus* from that level, the illustration of upper TT-zone *T. regina* in combat with a *T. prorsus* is accurate because the former was based on NHMAD “S”.

With the data and improved analysis now on hand, restorations of *Tyrannosaurus* need to be executed bearing the appropriate species specific structures (Fig. 11). Whether the illustration is new, or revised in accord with the new data. If an illustration is intended to represent *T. imperator*, then the supraorbital bosses should be spindles at least if they are intended to represent mature males, and *T. rex* and *T. regina* should never be shown with such (nor should *T. imperator* ever be shown with *Triceratops prorsus*), there are a few restorations of *Tyrannosaurus* that show the spindle (but such should never be used to represent the genus near or at the K/Pg boundary). If *T. rex* mature males are being illustrated, then supraorbital bosses need to be the vertically prominent knob discs limited to the anterior section of the postorbital, which should never be shown adorning *T. regina* or *T. imperator* (and neither *T. rex* nor *T. regina* should never be shown in the same scene with *T. horridus*), I have not seen these newly realized knob structures correctly illustrated.

## INSTITUTIONAL ABBREVIATIONS

AMNH: American Museum of Natural History, New York
BHI: Black Hills Institute, Hill City.
CM: Carnegie Museum of Natural History, Pittsburgh
CMNH: Cleveland Museum of Natural History, Cleveland
DMNS: Denver Museum of Nature and Science, Denver.
FMNH: Field Museum of Natural History, Chicago
HMN: Humboldt Museum fur Naturkunde, Berlin
LACM: Los Angeles County Museum, Los Angeles
LL: The Manchester Museum, Manchester.
MNHN: Museum national d’Histoire naturalle of Paris, Paris.
MOR: Museum of the Rockies, Bozeman.
NCMNS: North Carolina Museum of Natural Sciences, Raleigh
NHMAD: Natural History Museum Abu Dhabi, Abu Dhabi.
NHMUK: Natural History Museum, London
PIN: Paleontological Institute, Moscow.
RGM: Rijksmuseum van Geologie en Mineralogie, Leiden.
RMDRC: Rocky Mountain Dinosaur Resource Center, Colorado Springs
ROM: Royal Ontario Museum, Toronto.
RSM: Royal Saskatchewan Museum, Regina.
SDSM: Museum of Geology, South Dakota School of Mines and Technology, Rapid City.
SMNH: Saskatchewan Museum of Natural History, Regina
TCM: The Children’s Museum, Indianapolis.
TMP: Royal Tyrrell Museum of Palaeontology, Drumheller
TMT: Tate Museum, Casper.
UCMP: University of California Museum of Paleontology, Berkeley
UMNH: Utah Museum of Natural History, Salt Lake City.
USNM: National Museum of Natural History, Smithsonian, Washington DC
UWBM: University of Washington Burke Museum of Nature and Culture, Seattle
YPM: Yale Peabody Museum, New Haven.

## Corrections to Paul et al. (2022)

The number of specimens examined was 37.

In the systematics section AMNH 9340 is a typo, it is 9380.

In Table 1 USNM 555000 in referred to by its MOR number 555 in one location.

In Table 1 the BHI 3033 metatarsal 3 diameter has been remeasured at 77 mm.

In Table 1 the BHI 6230 metatarsal 2 diameter of 64 mm is a typo, it is 74 mm.

In Table 1 in the section containing tooth dimension data the column heading Hum Ratio is a typo, it should be Inc Rat

## REFERENCES

1. Alvares, F. et al. (2019). Old World spp. with taxonomic ambiguity: Workshop conclusions and recommendations. IUCN/SSC Canid Specialist Group.

2. Ashworth, J. (2022). Controversial paper suggests there are three *Tyrannosaurus* species. NHM, https://www.nhm.ac.uk/discover/news/2022/march/controversial-paper-suggests-there-are-three-tyrannosaurus-species.html.

3. Bakker, R. T., Williams, M. & Currie, P. J. (1988). *Nanotyrannus*, and new genus of pygmy tyrannosaur. Hunteria, 1, 1–30.

4. Barnosky, A. D. & Bell, C. (2004). Evolution, climatic change and species boundaries: Perspectives from tracing *Lemmiscus curtatus* populations through time and space. *Proceedings of the Royal Society B*. Biological Sciences, 270, 2585–2590.

5. Barras, C. (2022). *Tyrannosaurus rex* may actually be three separate species; New Scientist, https://www.newscientist.com/article/2308160-tyrannosaurus-rex-may-actually-be-three-separate-species.

6. Barrett, P. M., Butler, R. J. & Knoll, F. (2005). Small-bodied ornithischian dinosaurs from the Middle Jurassic of Sichuan, China. Journal of Vertebrate Paleontology, 25, 823–834.

7. Bates, K. T. et al. (2009). Estimating mass properties of dinosuars using laser imaging and 3D computer modeling. PLoS ONE, 4, e4532.

8. Brochu, C. A. (2003). Osteology of *Tyrannosaurus rex*: insights from a nearly complete skeleton and high-resolution computed tomographic analysis of the skull. Journal of Vertebrate Paleontology, 22, *Memoir* *7*, 1-138.

9. Brown, C. M., Currie, P. J. & Therrien, F. (2022). Intraspecific facial bite marks in tyrannosaurids provide insight into sexual maturity and evolution of bird-like intersexual display. Paleobiology, 48, 2–43.

10. Brown, C. M. et al. (2017). An exceptionally preserved three-dimensional armored dinosaur reveals insights into colorations and Cretaceous predator-prey dynamics. Current Biology, 27, 1–8.

11. Brusatte, S. L., Benson, R. B. J. & Norell, M. A. (2011). The anatomy of *Dryptosaurus aquilunguis* and a review of its tyrannosauroid affinities. American Museum Novitates, 3717, 1–53.

12. Brusatte, S., Carr, T. D. & Norell, M. A. (2012). The osteology of *Alioramus*, a gracile and long-snouted tyrannosaurid from the Late Cretaceous of Mongolia. Bulletin of the American Museum of Natural History, 366, 1–197.

13. Campbell, J. A., Ryan, M. J, Holme, R. B., & Schroder-Adams, C. .J. (2016). A re-evaluation of the chasmosaurine ceratopsid genus *Chasmosaurus* from the upper Cretaceous Dinosaur Park Formation of western Canada. PLoS ONE, 11, e0145805.

14. Carabajal, A. P., et al. (2021). Two braincases of *Daspletosaurus*: anatomy and comparison. Canadian Journal of Earth Sciencs, 58, 885–910.

15. Carpenter, K. (1990). Variation in *Tyrannosaurus rex*. In K. Carpenter & P. J. Currie (Eds.), Dinosaur Systematics: Perspectives and Approaches (pp. 141–145). Cambridge: Cambridge University Press.

16. Carpenter, K. (2010). Species concept in North American stegosaurs. Swiss Journal of Geosciences, 103, 155–162

17. Carpenter, K., & Smith, M. (2001). Forelimb osteology and biomechanics of *Tyrannosaurus rex*. In D. Tanke & K. Carpenter (Eds.) Mesozoic Vertebrate Life (pp. 90–116), Bloomington: Indiana University Press.

18. Carr, T. D. (2020). A high-resolution growth series of *Tyrannosaurus rex* obtained from multiple lines of evidence. PeerJ, 8, e9192.

19. Carr, T. D. (2022). All About *T. rex* in the 21^st^ Century, https://urldefense.com/v3/ https://www.youtube.com/watch?v=FBMZkDsAczE;!!LIr3w8kk_Xxm!rkdUHp2_OBeLKQKUmPIDiNITX8iZpmiDkhK9gQOWaRPZPFPEFvBcRcPYy2WtOs7MbC52DFo0jVAXmPGva74$.

20. Carr, T. D., Williamson, T. E. & Schwimmer. D. R. (2005). A new genus and species of tyrannosauroid from the Late Cretaceous Demopolic Formation of Alabaama. Journal of Vertebrate Paleontology, 25, 119–143.

21. Carr, T. D. & Williamson, T. E. (2010). *Bistahieversor sealeyi*, gen. et sp. nov., a new tyrannosaurid from New Mexico and the origin of deep snouts in Tyrannosauroidea. Journal of Vertebrate Paleontology, 30, 1–16.

22. Carr, T.D., Williamson, T.E., Britt, B.B. & Stadtman, K. (2011). Evidence for high taxonomic and morphologic tyrannosaurid diversity in the Late Cretaceous of the American Southwest and a new short-skulled tyrannosaurid from the Kaiparowits formation of Utah. Naturwissenschaften, 98, 241–246.

23. Carr, T. D., et al. (2017). A new tyrannosaur with evidence for anagenesis and crocodile-like facial sensory system. Scientific Reports,7, 44942.

24. Carr, T. D. et al. (2022). Insufficient evidence for multiple species of *Tyrannosauru*s in the latest Cretaceous of North America: A comment on “The tyrant lizard king, queen and emperor: Multiple lines of evidence support subtle evolution and probable speciation within the North American genus *Tyrannosaurus*.” Evolutionary Biology, preprint.

25. Coimbra, R. T. F. et al. (2021). Whole-genome analysis of giraffe supports four distinct species. Current Biology, 32, 2929–2938.

26. Cope, E. D. (1892). Fourth note on the dinosauria of the Laramie. American Naturalist, 26, 756–758.

27. Cullen, T. M. et al. (2020). Osteohistological analyses reveal diverse strategies of theropod dinosaur body-size evolution. Proceedings of the Royal Society B, 287, 2020.2258.

28. Currie, P. J. (2003a). Cranial anatomy of tyrannosaurids form the Late Cretaceous of Alberta. Acta Palaeontologica Polonica, 48, 191–226.

29. Currie, P. J. (2003b). Allometric growth in tyrannosaurids from the Upper Cretaceous of North America and Asia. Canadian Journal of Earth Sciences, 40, 651–665.

30. Currie, P. J. & Russell, D. A. (2005). The geographic and stratigraphic distribution of articulated and associated dinosaur remains. In P. J. Currie & E. B. Koppelhus (Eds.), Dinosaur Provincial Park: A Spectacular Ancient Ecosystem Revealed (pp. 537–569). Bloomington: Indiana University Press.

31. Dalman, S. G. (2013). New Examples of *Tyrannosaurus rex* from the Lance Formations of Wyoming, United States. Bulletin of the Peabody Museum of Natural History, 54, 241–254.

32. Davis, N. (2022). *Tyrannosaurus rex* may have been three species, scientists say. The Guardian, https://www.theguardian.com/science/2022/mar/01/tyrannosaurus-rex-may-have-been-three-species-scientists-say.

33. Dodson, P. (1975). Taxonomic implications of relative growth in lambeosaurine hadrosaurs. Systematic Zoology, 24, 37–54.

34. Dooley, A. C., et al. (2019). *Mammut pacificus* sp. nov., a newly recognized species of mastodon from the Pleistocene of western North America. PeerJ, 7, e6614.

35. Dunham, W. (2022). Scientists propose *Tyrannosaurus* had three species, not just ‘*rex*.’ Reuters, https://www.reuters.com/business/environment/scientists-propose-tyrannosaurus-had-three-species-not-just-rex-2022-03-01.

36. Eberth, D. A. & Kamo, S. L. (2019). First high-precision U-Pb CA-ID-TIMS age for the Battle Formation, Red Deer River Valley, Alberta, Canada: Implications for ages, correlations, and dinosaur biostratigraphy of the Scollard, Frenchman, and Hell Creek Formations. Canadian Journal of Earth Sciences, 56, 1041–1051.

37. Elbein, A. (2022). *T. rex* might have had close cousins. The New York Times, 171(59,349), D1,4, https://www.nytimes.com/2022/02/28/science/tyrannosaurus-rex-species.html.

38. Erickson, G. M., et al. (2004). Gigantism and comparative life-history parameters of tyrannosaurid dinosaurs. Nature, 430, 772–775.

39. Evans, D. C. & Reisz, R. R. (2007). Anatomy and relationships of *Lambeosaurus magnicristatus*, a crested hadrosaurid dinosaur from the Dinosaur Park Formation, Alberta. Journal of Vertebrate Paleontology, 27, 373–393.

40. Fowler, D. W. (2017). Revised geochronology, correlation, and dinosaur stratigraphic ranges, correlation, and dinosaur stratigraphic ranges of the Santonian-Maastrichtian formations of the Western Interior of North America. PLoS ONE, 12, e0188426.

41. Fowler, D. W. & Freedman, E. A. F. (2020). Transitional evolutionary forms in chasmosaurine ceratopsid dinosaurs: evidence from the Campanian of New Mexico. PeerJ, 8, e9251.

42. Galton, P. M. (1981). *Dryosaurus*, a hypsilophodontid dinosaur from the Upper Jurassic of North America and Africa postcranial skeleton. Palaontologische Zeitschrift, 55, 271–312

43. Gamble, K. C. (2007). Internal anatomy of the hornbill casque described by radiography, contrast radiography, and computed tomography. Journal of Avian Medicine and Surgery, 21, 38–49.

44. Gignac, P. & Erickson, G. M. (2017). The biomechanics behind extreme osteophagu in *Tyrannosaurus rex*. Scientific Reports, 7, 2012.

45. Gould, S. J. (2002). The Structure of Evolutionary Theory. Cambridge: Harvard University Press.

46. Greshko, M. (2020). Should *T. rex* be 3 species? New study sparks fierce debate. National Geographic, https://www.nationalgeographic.com/science/article/call-to-split-tyrannosaurus-rex-into-3-species-sparks-fierce-debate.

47. Grubb, P. et al. (2000). Living African elephants belong to two species: *Loxodonta africana* and *Loxodonta cyclotis*. Elephant, 2, 1–4.

48. Harvati, K. & Ackermann, R. R. (2022). Hybridization in the Late Pleistocene: Merging morphological and genetic evidence. bioRxiv, 2022.04.20.488874.

49. Head, J. J., Barrett, P. M. & Rayfield, E. J. (2009). Neurocranial osteology and systematic relationships of *Varanus* (*Megalania) 85prisca* Owen, 1859. Zoological Journal of the Linnean Society, 155, 445-457.

50. Hernandez, D. (2022). The mighty T-Rex could actually be three separate dinosaur species. The Wall Street Journal, https://www.youtube.com/watch?v=HwFV35rBoEc.

51. Hone, D. W. E. & Naish, D. (2013) The ‘species recognition hypothesis’ does not explain the presence and evolution of exaggerated structures in non-avialan dinosaurs. Journal of Zoology, 290, 172–180.

52. Hoyo, J. et al. (1992). *Handbook of the Birds of the World. Barcelona*, V 1. Lynx Edicions.

53. Hoyo, J. et al. (1994). *Handbook of the Birds of the World. Barcelona*, V 2. Lynx Edicions.

54. Hoyo, J. et al. (1996). *Handbook of the Birds of the World. Barcelona*, V 3. Lynx Edicions.

55. Hoyo, J. et al. (1997). *Handbook of the Birds of the World. Barcelona*, V 4. Lynx Edicions.

56. Hoyo, J. et al. (2001). *Handbook of the Birds of the World. Barcelona*, V 6. Lynx Edicions.

57. Hunt, G., Hopkins, M. J., & Lidguard, S. (2015). Simple versus complex models of trait evolution. Proceedings of the National Academy of Sciences, 112, 4885–4890.

58. Hunt, K. (2022) *Tyrannosaurus rex* may have been misunderstood. CNN, https://www.cnn.com/2022/02/28/world/t-rex-three-different-dinosaurs-scn/index.html.

59. Hurum, J. H. and Sabath, K. (2003). Giant theropod dinosaurs from Asia and North America: Skulls of *Tarbosaurus bataar* and *Tyrannosaurus rex* compared. Acta Paleontologica Polonica, 48, 161–190.

60. Hutchinson, J. R, et al. (2011). A computational analysis of limb and body dimensions in *Tyrannosaurus rex* with implications for locomotion, ontegeny, and growth. PLoS ONE, 6, e26037.

61. Jevnikar, E. M. & Zanno, L. E. (2021). Bimodal trajectories and unresolved early growth stages in *Tyrannosaurus rex* growth. Society of Vertebrate Paleontology 2021 Annual Meeting Abstracts, 150-151.

62. Johnson, M. M., Young, M. T. & Brusatte, S. L. (2020). The phylogenetics of Teleosauridea and implications for the ecology and evolution. PeerJ, 8, e9808.

63. Kim, S. E. (2022). Is *T. rex* really three royal species” Paleontologists cast doubt over new claims. Popular Science, https://www.popsci.com/animals/t-rex-different-species-debate.

64. Knapp, A. et al. (2018). Patterns of divergence in the morphology of ceratopsian dinosaurs: sympatry is not a driver of ornament evolution. Proceedings of the Royal Society B, 285, 20180312.

65. Knutsen, E. M. 2012. A taxonomic revision of the genus *Pliosaurus*. Norwegian Journal of Geology, 92, 259–276.

66. Koepfli, K.-P. et al. (2015). Genome-wide evidence reveals that African and Eurasian golden jackals are distinct species. Current Biology, 25, 2185–2165.

67. Kruger, G. & Ricci, S. (2022). *Tyrannosaurus rex* has two new sister species? *I Know Dino*, https://podcasts.apple.com/us/podcast/tyrannosaurus-rex-has-two-new-sister-species/id960976813?i=1000553526816.

68. Larramendi, A. Zhang, H., Palombo, M. R., & Ferretti, M. P. (2020). The evolution of *Palaeoloxodon* skull structure: Disentangling phylogenetic, sexually dimorphic, ontogenetic, and allometric morphological signals. Quaternary Science Reviews, 229, 106090.

69. Larramendi, A., Paul, G. S. & Hsu, S. (2021). Review and reappraisal of the specific gravities of present and past multicellular organisms, with an emphasis on vertebrates, particularly pterosaurs and dinosaurs. Anatomical Record, 304, 1833–1888.

70. Larson, N. (2008). One hundred years of *Tyrannosaurus rex*: The skeletons. In P. Larson and K. Carpenter (Eds.) Tyrannosaurus rex: The Tyrant King. (pp. 1–55). Bloomington: Indiana University Press.

71. Larson, P. (1994). Tyrannosaurus sex. Paleontological Society Special Publication, 7, 139–155.

72. Larson, P. (2008). Variation and sexual dimorphism in *Tyrannosaurus rex*. In P. Larson and K. Carpenter (Eds.) Tyrannosaurus rex: The Tyrant King. (pp. 103–130). Bloomington: Indiana University Press.

73. Larson, P. (2013). The validity of Nanotyrannus lancensis. Society of Vertebrate Paleontology 2013 Annual Meeting Abstracts, 159.

74. Lawrence, I. (2022). Was there *Tyrannosaurus imperator* & a *regina*? *Terrible Lizards*, https://www.youtube.com/watch?v=bgZUE8tiLGs.

75. Lister, A. M. & Sher, A. V. (2015). Evolution and dispersal of mammoths across the northern hemisphere. Science, 350, 805–809.

76. Loewen, M. A. et al. (2013). Tyrant dinosaur evolution tracks the rise and fall of Late Cretaceous oceans. PLoS ONE, 8, e79420.

77. Long, K. L., Prothero, D. L., & Syverson, V. J. P. (2020). How do small birds evolve in response to climate change? Data from the long-term record at La Brea tar pits. Integrated Zoolology, 15, 249–261.

78. Lü, J., Yi, L., Brusatte, S. L., Yang, L. & Chen, L. (2014). A new clade of Asian Late Cretaceous tyrannosaurids. Nature Communications, 5, 3788.

79. Macdonald, I. & Currie, P. J. 2018, Description of a partial *Dromiceiomimus* skeleton with comments on the validity of the genus. Canadian Journal of Earth Sciences, 56, 129–157.

80. Mader, B. J. (2010). A species-level revision of the North American brontotheres *Eotitanops* and *Palaeosyops*. Zootaxa, 2339, 1–43

81. Maisch, M.W. (2008). Revision der Gattung *Stenopterygius* Jaekel, 1904 emend. von Huene, 1922 aus dem unteren Jura Westeuropas. Palaeodiversity, 1, 227–271.

82. Maleev, E. A. (1955). Giant carnivorous dinosaurs of Mongolia. Doklady Akademii Naul SSSR, 104, 634–637.

83. Maleev, E. A. (1974). Giant carnosaurs in the family Tyrannosauridae. *Sovmestnaa Sovetsko-Mongolskaa Paleontologicesaa Ekspedicia*, Trudy 1, 132–191.

84. Mallon, J. C. et al. (2022). The record of *Torosaurus* in Canada and its taxonomic implications. Zoological Journal of the Linnean Society, 195, 157–171.

85. Marghoub, A. et al. (2022). Unraveling the structural variation of lizard osteoderms. Acta Biomaterialia, 146, 306–316.

86. Maxwell, E. E. 2012. New metrics to differentiate species *of Stenopterygius* from the Lower Jurassic of southwestern Germany. Journal of Paleontology, 86, 105–115.

87. Mayr, E. (1982). The Growth of Biological Thought, Diversity, Evolution and Inheritance. Harvard University Press: Cambridge.

88. Mayr, G. (2018). A survey of casques, frontal humps, and other extravagant bony cranial protuberances in birds. Zoomorphology, 137, 457–472.

89. Mertens, R. 1942 Die Familie der Warane (Varanidae), Abhandlungen der Senckenbergischen Naturforschenden Gesellschaft: Abhandlung 462.

90. Mihlbachler, M. C. (2008). Species taxonomy, phylogeny, and biogeography of the Brontotheriidae. Bulletin of the American Museum of Natural History, 311, 1–475.

91. Molnar, Ralph E. (2004). Dragons in the Dust: the Paleobiology of the Giant Monitor Lizard Megalania. Bloomington: Indiana University Press.

92. Nowak, R. M. 1991. Walker’s Mammals of the World. Baltimore, The Johns Hopkins University Press.

93. Osborn, H. F. (1905). *Tyrannosaurus* and other Cretaceous carnivorous dinosaurs. Bulletin of the American Museum of Natural Hisory, 21, 259–265.

94. Osborn, H. F. (1917). Skeletal adaptations of Ornitholestes, Struthiomimus, Tyrannosaurus. Bulletin of the American Museum of Natural History, 35, 733–771.

95. Oyston, J. W., Wilkinson, M., Ruta, M. & Wills, M. A. (2020) Molecular phylogenies map to biogeography better than morphological ones. Communications Biology, 5. 521.

96. Padian, K. & Horner, J. (2011). The evolution of ‘bizarre structures’ in dinosaurs: biomechanics, sexual selection, social selection, or species recognition? Journal of Zoology, 283, 3–17.

97. Padian, K. & Horner, J. R. (2014). The species recognition hypothesis explains exaggerated structures in non-avialan dinosaurs better than sexual selection does. Comptes Rendus Palevol, 13, 97–107.

98. Paul, G. S. (1988). Predatory Dinosaurs of the World. New York: Simon & Schuster.

99. Paul, G. S. (1997). Dinosaur models: the good, the bad, and using them to estimate the mass of dinosaurs. In D. L. Wolberg, E. Stump and G. D. Rosenberg (Eds.) Dinofest International Symposium Proceedings, Academy of Natural Sciences, *Philadelphia*. (pp. 129–154). Philadelphia: Academy Sciences of Natural Sciences.

100. Paul, G. S. (2006). A revised taxonomy of the iguanodont dinosaur genera and species. Cretaceous Research, 29, 192–216.

101. Paul, G. S. (2008). The extreme lifestyles and habits of the gigantic tyrannosaurid superpredators of the Late Creaceous of North America and Asia. In P. Larson and K. Carpenter (Eds.) Tyrannosaurus rex: The Tyrant King. (pp. 307–352). Bloomington: Indiana University Press.

102. Paul, G. S. (2016). Princeton Field Guide to Dinosaurs. Princeton: Princeton University Press.

103. Paul, G. S. (2018). Nonornithischian dinosaurs did too have lips, probably big lips, here’s why. Prehistoric Times, 127, 44–49.

104. Paul, G. S. (2019). Determing the largest known land animal: A critical comparison of differing methods for restoring the volume and mass of extinct animals. Annals of Carnegie Museum, 85, 335–358.

105. Paul, G. S. (2022a). The Princeton Field Guide to Pterosaurs. Princeton: Princeton University Press.

106. Paul, G. S. (2022b). Splitting *Tyrannosaurus*. Prehistoric Times, 141, 32-33, 44–48.

107. Paul, G. S., Persons, W. S. & Van Raalte, J. 2022. The tyrant lizard king, queen and emperor: Multiple lines of evidence support subtle evolution and probable speciation within the North American genus *Tyrannosaurus*. Evolutionary Biology, 49, 156–179.

108. Perri, A. R. et al. (2021). Dire wolves were the last of an ancient New World canid lineage. Nature, 591, 87–91.

109. Rinehart, L. F. et al. (2009). The paleobiology of *Coelophysis bauri* from the Upper Triassic Whitaker quarry, New Mexico, with a detailed analysis of a single quarry block. The New Mexico Museum of Natural History & Science, 45, 1–260.

110. Russell, D. A. (1970). Tyrannosaurs from the Late Cretaceous of western Canada. National Museum of Natural Sciences Publications in Paleontology, 1, 1–34.

111. Scannella, J. B, Fowler, D. W., Goodwin, M. B. & Horner, J. R. (2014). Evolutionary trends in *Triceratops* from the Hell Creek Formation, Montana. PNAS, 111, 10245–10250.

112. Schweitzer, M. H., et al. (2016). Chemistry supports the identification of gender-specific reproductive tissue in *Tyrannosaurus rex*. Science Reports, 6, 23099.

113. Sellers, W. I. et al. (2017). Investigating the running abilities of Tyrannosaurus rex using stress-constrained multibody dynamic analysis. PeerJ, 5, e3420.

114. Sereno, Paul C. (2010). Taxonomy, cranial morphology, and relationships of parrot- beaked dinosaurs. In Ryan, Michael J.; Chinnery-Allgeier, Brenda J.; Eberth, David A. (eds.). New Perspectives on Horned Dinosaurs: The Royal Tyrrell Museum Ceratopsian Symposium. (pp. 21–58). Bloomington: Indiana University Press.

115. Snively, E., Henderson, D. M., Philips, D. S. (2006). Fused and vaulted nasals of tyrannosaurid dinosaurs: Implications for cranial strength and feeding mechanics. Acta Palaeontologica Polonica, 51, 435–454.

116. Tschopp, E., Mateus, O. V., & Benson, R. B. (2015). A specimen-level phylogenetic analysis and taxonomic revision of Diplodocidae. PeerJ, 3, e857.

117. Vickaryous, M. K., Meldrum, G. and Russell, A. P. (2015). Armored geckos: A histological investigation of osteoderm development in *Tarantola* and *Gekko* with comments on their regeneration and inferred function. Journal of Morphology, 276, 1345–1357.

118. Witton, M. P. (2022). Tyannouroboros: how everything old is new again in recent proposals of *Tyrannosaurus* taxonomy. Mark P. Witton’s blog, https://www.timescall.com/2022/03/06/scott-rochat-rochat-can-you-see-triple-your-t-rex-triple-your-fun.

119. Zeinio, K. (2012). Species Concepts. Scientific American, https://blogs.scientificamerican.com/evo-eco-lab/species-concepts.

